# Validating Conditionally Essential Targets: Discovery of the First Orally Effective Biotin Inhibitor against *Mycobacterium Tuberculosis*

**DOI:** 10.1101/2025.09.24.678246

**Authors:** Qiang Liu, Joshua B. Wallach, Yahani P. Jayasinghe, Mark R. Sullivan, Julianna Proietto, Suyapa Rodriguez, Sang Vo, Helena I. M. Boshoff, Ziyi Jia, Camilla Folvar, Betelhem Tatek, Firat Kaya, Lev Ostrer, Kritee Mehdiratta, Rui Shi, Véronique Dartois, Anthony D. Baughn, Eric J. Rubin, Donald R. Ronning, Matthew D. Zimmerman, Dirk Schnappinger, Courtney C. Aldrich

**Author notes:** Correspondence: Courtney C. Aldrich, Mailing address: Department of Medicinal Chemistry, University of Minnesota, 308 Harvard Street, Weaver-Densford Hall, Room 8-101, Minneapolis, Minnesota 55455, United States, Dirk Schnappinger, Mailing address: Department of Microbiology and Immunology, Weill Cornell Medical College, New York 10021, United States, Matthew D. Zimmerman, Mailing address: Center for Discovery and 1820 Innovation, Hackensack Meridian Health, Nutley, New Jersey 1821 07110, United States. contributed equally (Qiang Liu, Joshua B. Wallach).

## Abstract

Conditionally essential pathways - such as the biotin biosynthesis - represent promising targets for new antibiotics. However, the chemical interrogation of the biotin pathway with an orally effective lead remains elusive, and the preclinical development of biotin inhibitors for mycobacterial infections in vivo is challenging due to the unusually high concentration of biotin in standard mouse models. Structure-guided optimization was applied to develop the first oral lead targeting aminotransferase BioA, a key enzyme in bacterial biotin biosynthesis, resulting in **C48**, a picomolar inhibitor displaying sub-micromolar MICs against *Mycobacterium tuberculosis* (Mtb). Mechanism of action was confirmed by biochemical, structural, and genetic studies. **C48** demonstrated favorable pharmacokinetics and excellent oral bioavailability resulting in over 39,000-fold improved exposure. We next developed an easy-to-operate, low-biotin mouse model that recapitulates human biotin physiology. **C48** significantly reduced Mtb burden in this low-biotin mouse model, providing the first in vivo proof-of-concept for targeting biotin biosynthesis in Mtb.

## INTRODUCTION

Antimicrobial resistance (AMR) is a rapidly intensifying global health crisis, fueled by overuse of antibiotics, incomplete treatment, environmental contamination, and inadequate sanitation^1^. The 2024 WHO Bacterial Priority Pathogen List highlights the severity of this threat by identifying 15 high-priority drug-resistant bacteria such as rifampicin-resistant *Mycobacterium tuberculosis* and carbapenem-resistant *Acinetobacter baumannii* and *Enterobacterales*^2^. Biotin is an essential cofactor for lipid metabolism and particularly critical for Mtb, which features a complex, lipid rich cell envelope that underpins its virulence, pathogenesis and drug resistance^3^. Genetic studies have confirmed that Mtb must synthesize biotin to grow and persist in mice^4–6^ and remarkably, a comprehensive study across 60 genetically diverse mouse strains found that biotin-deficient Mtb mutants suffered an average reduction in fitness of almost 400-fold, underscoring the essentiality of biotin synthesis for survival of Mtb in vivo^7^.

Despite advances in genetic validation of biotin synthesis as a drug target, attempts to develop an orally bioavailable lead compound have failed. Nevertheless, the natural products amiclenomycin and acidomycin, which selectively target BioA and BioB, respectively, identified the biotin synthesis pathway (Supplementary Fig. S1) as vulnerable to chemical inhibition^8, 9^. Further phenotypic and target-based screening identified several novel chemotypes including (nitrophenylthio)-acetohyrazide MAC13772^10^ and aryl hydrazine^11^. These compounds proved to be valuable chemical probes for a diverse of pathogens, including *Acinetobacter* spp., *E. coli*, *K. pneumoniae*, and *Mycobacterium* spp. However, further development of these antibiotic scaffolds has been halted due to their inherent instability or limited chemical tractability.

Historically, antibiotic development has focused on bacterial gene products that are essential for growth in nutrient-rich media or animal models that often fail to replicate the human infection environment^12^. This limitation has been increasingly recognized, as drug targets that were overlooked in conventional systems are now being identified in host-mimicking conditions, for example, serum-based or nutrient-limited condition to recapitulate the starvation in human^10, 13, 14^. Those efforts uncovered previously overlooked, conditionally essential pathways, revealing a promising new frontier for antibiotic development. However, the preclinical development of conditionally essential inhibitors for mycobacterial infections in vivo faces a key translational hurdle as illustrated by biotin in this report. In humans, where biotin is scarce (1 nM), Mtb is expected to be more vulnerable to inhibition of biotin synthesis than in conventional mouse models of TB, which exhibit ∼40-fold higher circulating biotin levels (∼40 nM). Consequently, Mtb can likely scavenge sufficient biotin from the murine host to offset the effects of partial biotin synthesis inhibition, leading to a false-negative result in efficacy studies^15^. Brown *et al* developed a mouse model mimicking human biotin content for ∼12 hour by single streptavidin i.v. injection and demonstrated biotin auxotrophs of *A. baumannii, Pseudomonas aeruginosa*, and *Klebsiella pneumoniae* were unable to proliferate without biotin supplementation^15^. Unfortunately, this model is not suited for studying slow-growing bacteria like Mtb, as the ∼12 hour period of biotin depletion window is too brief to monitor the chronic stages of infection which routinely takes weeks in animal studies.

Here we describe the development of the first orally available biotin inhibitor, **C48**, that specifically targets biotin synthesis exhibits potent activity against drug-sensitive and drug-resistant mycobacteria. Additionally, we established an easy-to-operate, low-biotin mouse model of TB infection that better recapitulates human biotin physiology with sustained duration of time. **C48** exhibits robust efficacy against Mtb in vivo. These studies provide the first proof-of-concept that effective inhibition of biotin biosynthesis in vivo is achievable with an orally bioavailable small-molecule. Furthermore, it emphasizes the untapped potential of conditionally essential targets and screening strategies in nutrient-limited condition that better mimic physiological conditions for developing novel antibiotics^10, 16–19^.

## RESULTS

### Structure-guided design of C48

Our initial hit **6** (Fig. 1A) was originally identified in a biochemical screen for inhibitors of the aminotransferase BioA and selected as a template based on its promising activity and modular structure^20^. Compound **6** demonstrated poor pharmacokinetic properties in CD-1 mice (Supplementary Table S1), exhibiting high clearance (30.6 mL/kg/min) and negligible oral exposure (*F* = 3.4%), highlighting critical need for further optimization. A co-crystal structure of **6** with BioA revealed opportunities to enhance potency and selectivity through π-stacking and van der Waals interactions since most of the side-chains were engaged with hydrogen-bonds with other protein atoms^21^. In parallel with the structure-guided design to boost potency, we also implemented rational design strategies by introducing fluorine atoms and reducing rotatable bonds to improve pharmacokinetics. Recognizing the benzodioxole as a metabolic liability^22^, we replaced it with a 3-bromo-4-fluorophenyl moiety, which also improved shape complementarity to the P1 pocket. Modification of the *N*-aryl fragment was performed to enhance potency and physicochemical properties by removal of a rotatable bond^23^ through annulation of the acetyl group onto the phenyl ring (Fig. 1A). These modifications yielded **C21**, which demonstrated superior potency with an IC_50_ of 48 nM and MIC_50_ of 1.9 μM against Mtb in biotin-free medium (Fig. 1A and Fig. 1B). Notably, the strategic incorporation of fluorine and indanone group significantly improved the pharmacokinetic profile of compound **C21** (Supplementary Table S1). These modifications reduced its clearance to 5.3 mL/kg/min, resulting in an excellent oral exposure (AUC = 108 μg·h/mL), reasonable half-life (0.82 h), and outstanding bioavailability (*F* > 100%). Consequently, the oral exposure of **C21** is represented by an AUC/MIC ratio of 57, an 1,475-fold improvement over the original hit compound **6**. Detailed synthetic procedures and compound characterizations are provided in the Supplementary Materials and Methods (Supplementary Scheme S1). In biological systems, the stacking interactions between heterocycles and aromatic amino acids (Phe, Tyr and Trp) are critical for molecular recognition and inspired by Wheeler’s report on the cooperative enhancement of π-π stacking by proximal electron-withdrawing groups (e.g., carbonyl and nitrogen atoms)^24^. Therefore, we next sought to install a nitrogen atom within the indanone ring to maximize π-π stacking interactions with Trp64 and Trp65 in the P2 pocket, and found the 6-position was optimal (Fig. 1A and Fig. 1C, 1D and 1E)^24^. The resulting compound **C48** demonstrated a profound increase in activity (MIC_50_ = 0.093 µM) over the parent compound establishing it as the most potent inhibitor of biotin biosynthesis reported to date (Supplementary Table S2). Moreover, **C48** also demonstrated favorable pharmacokinetics (Supplementary Table S1), with excellent oral bioavailability (*F* > 100%) and reduced drug clearance (3.2 mL/kg/min) in CD-1 mice. This improved oral exposure (AUC = 146 μg·h/mL) resulted in a remarkable AUC (p.o.)/MIC value of 1570, representing a 39,000-fold enhancement over hit **6**. The following section details the subsequent biochemical and mechanistic investigation of **C48**.

**Figure 1.**
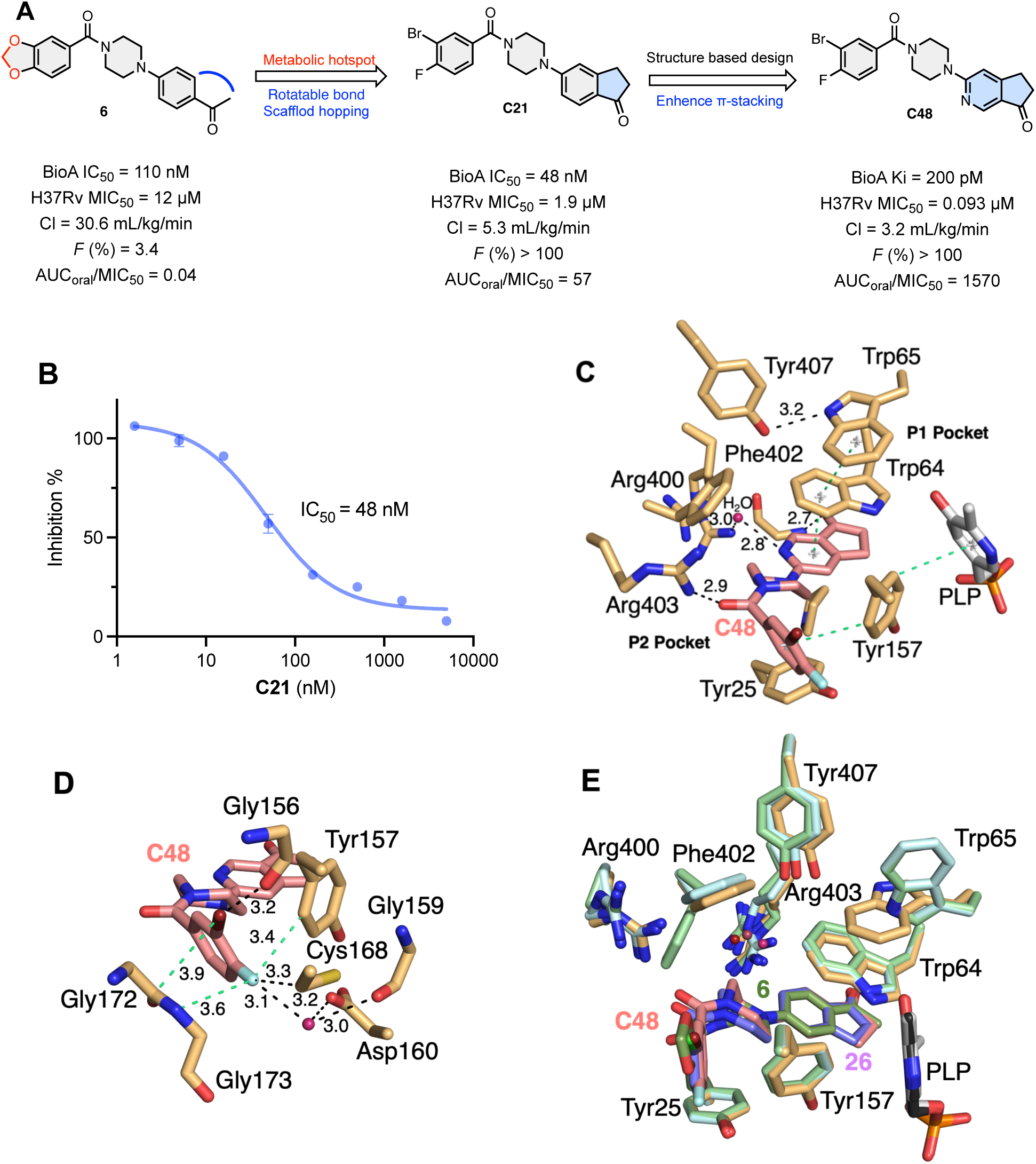
The discovery of potent biotin inhibitor C48. (**A**) Systematic in vitro and in vivo optimization campaign of compound **6**, derivatives **C21** and **C48** were identified as promising candidates: CL = drug clearance, *F* = oral bioavailability, AUC_oral_ = area under the plasma concentration−time curve administrated following oral administration (25 mg/kg). (**B**) The biochemical activity of **C21**. The assay was performed with **C21** (3-fold dilution ranging from 5000 to 1.6 nM in 50 nM BioA, 320 nM BioD, 3 µM KAPA, 1 mM S-adenosyl methionine, 20 nM Fluorescent-DTB tracer, 35 nM streptavidin, 5 mM ATP, 50 mM NaHCO_3_, 1 mM MgCl_2_, 0.1 mM PLP, 0.0025% Igepal CA630, and 100 mM Bicine [pH 8.6]. (**C**) Important π-π interactions that contribute to BioA binding of **C48**. The center of the π systems involved in important interactions are indicated by gray stars and connected by light green dashed bonds. Polar interactions are indicated by black dashed bonds with distances indicated. The carbon atoms of **C48** are pink and the carbon atoms of BioA are tan. Nitrogen, oxygen, phosphorous, fluorine, and bromine atoms are blue, red, orange, cyan, and maroon, respectively. (**D**) The additional halogen interactions formed in the P2 pocket as a result of halogen incorporation within the 3-bromo-4-fluorophenyl moiety. Colors are as indicated in panel B. (**E**) Superposition of BioA/PLP/**C48** (PDB:9D7M), BioA/PLP/**6** (PDB: 4XJP), and BioA/PLP/**26** complexes (PDB: 4XJO). The carbon atoms of 4XJP and 4XJO are in green and light blue, respectively. Compound **6** is in green and **26** is in purple. Heteroatom coloring is as indicated in panel B. Note the different conformation of Trp65 in the **C48** structure versus the other two ligand complexes.

To understand the molecular basis for **C48**’s increase in activity, we conducted detailed biochemical and structural analyses. In the BioA inhibition assay, **C48** exhibited an IC₅₀ of 34 nM (Fig. 2A), a value that approached half the total enzyme concentration (25 nM). We also observed a linear titration of activity with **C48,** a hallmark of tight binding, wherein the inhibitor quantitatively binds to the remaining free enzyme (Fig. 2B)^25^. By lowering the concentration of BioA to 5 nM and using the analysis of Morrison to account for inhibitor depletion, **C48** demonstrated an inhibition constant (*K*_i_) of 200 pM (Fig. 2C). Structural studies (Fig. 1C) further revealed that **C48** binds to the PLP-bound state of BioA rather than the pyridoxamine 5′-phosphate (PMP)-bound state, which indicates that **C48** is a competitive inhibitor with respect to KAPA (7-<u>k</u>eto-8-<u>a</u>mino-<u>p</u>elargonic <u>a</u>cid), rather than SAM (<u>S</u>-<u>a</u>denosyl <u>m</u>ethionine). This is consistent with a previous report that BioA inhibitor MAC13772 covalently attaches to PLP^15^.

**Figure 2.**
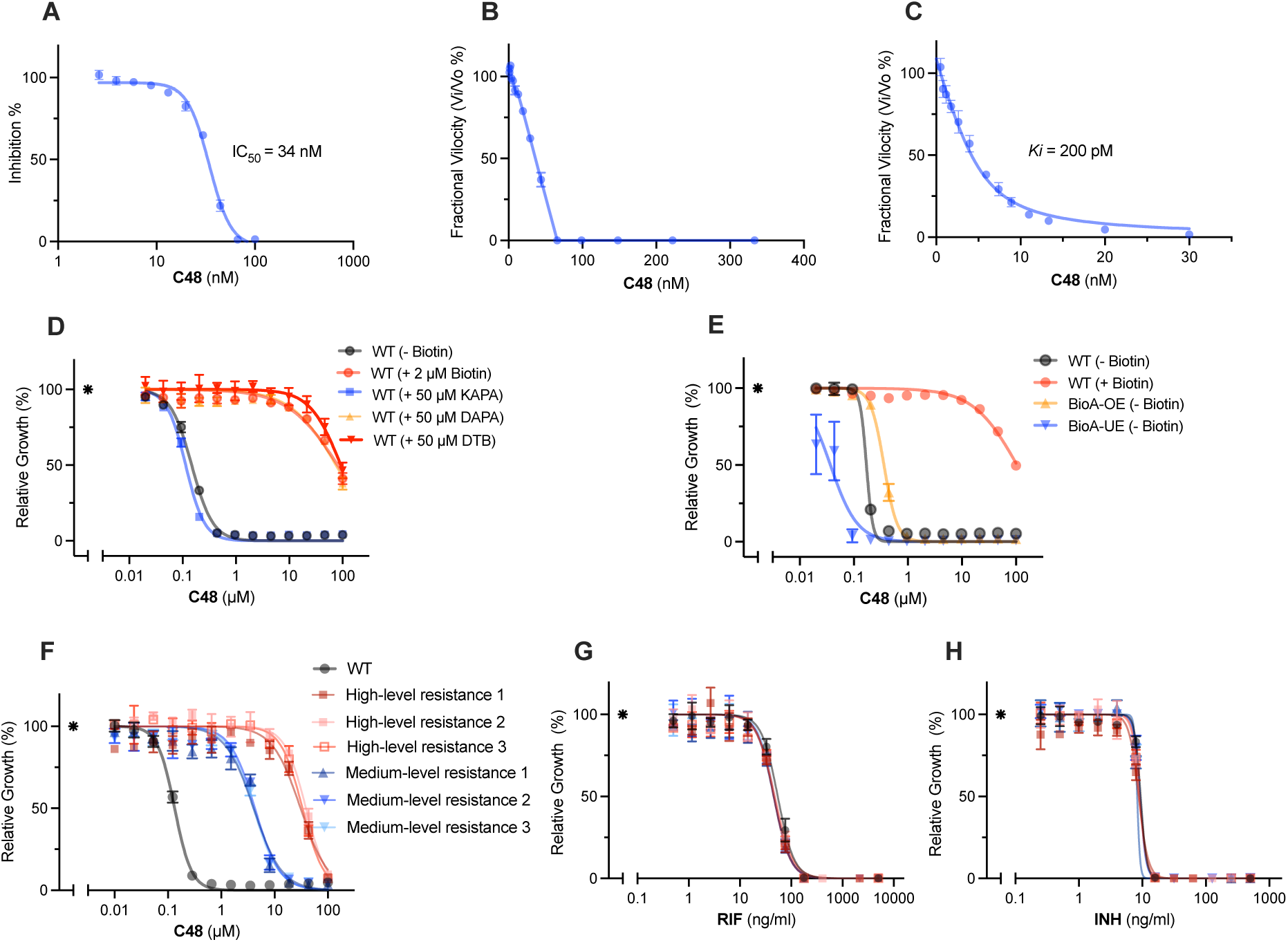
Biochemical and on-target validation of C48. **(A)** The biochemical activity of **C48**. The assay was performed with **C48** (1.5-fold dilution ranging from 100 to 2.6 nM) in 50 nM BioA, 320 nM BioD, 3 µM KAPA, 1 mM S-adenosyl methionine, 20 nM Fluorescent-DTB tracer, 35 nM streptavidin, 5 mM ATP, 50 mM NaHCO_3_, 1 mM MgCl_2_, 0.1 mM PLP, 0.0025% Igepal CA630, and 100 mM Bicine [pH 8.6]. (**B**) Linear titration of activity with **C48**. Initial *K*i was explored with **C48** (1.5-fold dilution ranging from 333 to 1.1 nM) in 50 nM BioA, 320 nM BioD, 3 µM KAPA, 1 mM S-adenosyl methionine, 20 nM Fluorescent-DTB tracer, 35 nM streptavidin, 5 mM ATP, 50 mM NaHCO_3_, 1 mM MgCl_2_, 0.1 mM PLP, 0.0025% Igepal CA630, and 100 mM Bicine [pH 8.6]. (**C**) The Ki of **C48**: The assay was performed with **C48** (30, 20, 13.33, 11.11, 8.89, 7.41, 5.93, 3.95, 2.63, 1.75, 1.17, 0.78 and 0.52 nM) in 5 nM BioA, 50 nM BioD, 10 µM KAPA, 7 mM SAM, 5 nM Fluorescent-DTB tracer, 9 nM streptavidin, 0.5 mM ATP, 50 mM NaHCO_3_, 1 mM MgCl_2_, 0.1 mM PLP, 0.0025% Igepal CA630, and 100 mM Bicine [pH 8.6]. (**D**). The activity of **C48** against Mtb was rescued by DAPA, DTB, Biotin, but not KAPA. (**E**) Sensitivity of **C48** against Mtb was subjected to the expression level of BioA, BioA-OE = BioA protein over-expressed strain and Bio-UE = BioA protein under-expressed strain compared to WT (H37Rv). (**F**) Isolated resistant mutants displayed MIC shifts against **C48**. (**G**) and (**H**). Isolated resistant mutants did not show susceptibility change against RIF (rifampicin) and INH (isoniazid). Asterisk in Fig. 2D-**H** represent no drug control. Biochemical assays were performed in triplicate and MIC experiments were performed at least twice.

The co-crystal structure of **C48** bound to BioA revealed that its tight-binding is driven by a combination of optimal shape complementarity and π-stacking interactions, extensive halogen bonded interactions, and additional hydrogen bonds that envelope the ligand (Fig. 1C, 1D, 1E and Supplementary Fig. S2, PDB: 9D7M, Diffraction data and summary refinement statistics are provided in Supplementary Table S3). Comparison of the BioA/**C48** complex to the published BioA/**6** (PDB: 4XJP) and BioA/**26** complex structures (PDB: 4XJO; has a similar chemical structure as **C21** but possesses a single chlorine in the benzoyl moiety) reveals important conformational differences and enhanced interactions along with the expected binding mode similarities described in our previous work^21^. The 3-bromo-4-fluorobenzoyl moiety occupies nearly the complete volume of the nonpolar P1 pocket and forms π-π interactions with Tyr157, which are enhanced by π-π interactions between Tyr 157 and the PLP cofactor (Fig. 1C). Replacement of the benzodioxole of **6** with a 3-bromo-4-fluorophenyl moiety in **C48** forms new and strengthened interactions with BioA. Indeed, the bromine and fluorine atoms are well positioned in the P1 pocket. The anisotropic electron distribution of heavy halogens like bromine produces a σ-hole, a partial positive charge on the bromine atom opposite the C-Br bond, as well as an electron-rich belt around the bromine atom perpendicular to the C-Br bond (Fig. 1D). This heterogenous electronic distribution affords diverse interactions with a variety of polar and non-polar functional groups that contributes to multiple interactions between **C48** and the BioA active site^21^. Specifically, this likely strengthens the interactions with the conjugated π system of the peptide bond between residues Gly172 and Gly173 as well as forming an interaction with the backbone carbonyl of Gly156. In contrast to bromine, electron density around the fluorine atom is more uniform. This affords polar interactions with the electron-depleted edge of the Tyr157 aromatic side chain moiety and a strongly-ordered water molecule that is coordinated by the side chains of Asp160, Cys168, and the backbone carbonyl of Gly159. In addition, the fluorine forms van der Waals interactions with Cys168 and Gly173.

The conformationally flexible central piperazine linker positioned between Tyr25 and Phe402 is oriented differently when compared to **6** and **26** complex structures but still forms hydrophobic interactions with those neighboring residues^21, 23^. The altered conformation repositions the carbonyl carbon linking the piperazinyl ring to the benzylhalogen ring by placing it closer to the side chain of Arg403 and thereby shortening the hydrogen bond between those groups. The azaindanone moiety projects into the P2 pocket of the BioA active site. Similar to the interactions observed in the BioA/**26** complex, the carbonyl oxygen of the cyclopentanone of the 3,4-bicyclic moiety forms a hydrogen bonded interaction with the backbone amide of Gly93. On one face of the azaindanone moiety, **C48** forms a hydrophobic interaction with Pro24. On the other face, the azaindanone forms a parallel displaced π-stacking interaction with Trp64 and Trp65. When compared with the BioA/**6** and BioA/**26** complex structures, the Trp65 side chain in the BioA/**C48** complex shows a significant conformational change affording increased π-π stacking interactions with Trp64 (Fig. 1E)^21^. This conformational change also forms an additional hydrogen bond between Trp65 and the side chain of Tyr407 found in the aforementioned π-π stacking network. In addition, the pyridine nitrogen of the bicyclic ring forms an important water mediated hydrogen bond with Arg400. Although Arg400 exhibits an alternative conformation in both monomers of the asymmetric unit, which suggests a relatively structurally dynamic character to this residue, both BioA molecules form the same water mediated interactions. As the incorporation of the aryl amine at this position appears to profoundly improve the potency of **C48**, this suggests that the formation of that specific hydrogen bond with this structured water molecule is a major contributor to binding affinity and specificity.

Overall, the strategic incorporation of a nitrogen atom on the same side as the carbonyl is the major driver of the enhancement in potency but that enhancement is aided by other novel interactions affording increased π-stacking energy and additional hydrogen bonds of the indanone fragment. This leads to positive cooperativity that may be further amplified by the dual tryptophan residues, which provide 40% stronger interactions than phenylalanine^24^. The incorporation of halogen to make the 3-bromo-4-florobenzoyl moiety also appear to contribute to the enhanced binding through strengthening the π-π interactions plus additional halogen bonded interactions. Taken together, this structural study elucidates the molecular basis of the enhancement in activity of **C48** relative to **6**.

### Mechanistic characterization of C48’s whole cell activity

The growth inhibitory effect of **C48** was antagonized by biotin, desthiobiotin (DTB) and 7, 9-<u>d</u>i<u>a</u>mino<u>p</u>elargonic <u>a</u>cid (DAPA), but not by 7-<u>k</u>eto-8-<u>a</u>mino-<u>p</u>elargonic <u>a</u>cid (KAPA)-the substrate of BioA-supporting BioA inhibition as the mechanism of **C48**’s antitubercular activity (Fig. 2D and Supplementary Fig. S1). To further confirm engagement of BioA, we evaluated **C48** against an engineered strain of Mtb H37Rv with ATc-inducible BioA overexpression. **C48** showed the expected monotonic changes in susceptibility to increasing BioA expression (Fig. 2E). **C48** was subsequently evaluated against a diverse panel of 70 Mtb isolates, including drug-resistant strains, representing a broad range of mutations covering known resistant mechanisms against first- and second-line TB drugs (Table 1, Supplementary Table S4 and S5). As a result, **C48** displayed remarkable sensitivity with MIC values of < 0.07 µM against a panel of susceptible and drug-resistant Mtb. The lack of cross-resistance of **C48** demonstrates the importance of developing new drugs with distinct modes of action as rifampicin-resistant Mtb has been included in the 2024 WHO Bacterial Priority Pathogen List for the first time^2^. For comparison, we tested the second- line TB drug linezolid to better calibrate the interpretation of our data and **C48** displayed superior potency than linezolid across the strains tested (Table 1, Supplementary Table S4 and S5). Nontuberculous mycobacteria (NTM) were also susceptible, exhibiting MICs ranging from 0.5-1 µM. In contrast, **C48** was inactive against the entire ESKAPE panel of pathogens (Supplementary Table S5), likely due to the low sequence similarity of BioA in mycobacteria and ESKAPE pathogens (Supplementary Fig. S3, Table S6 and Table S7).

**Table 1.**
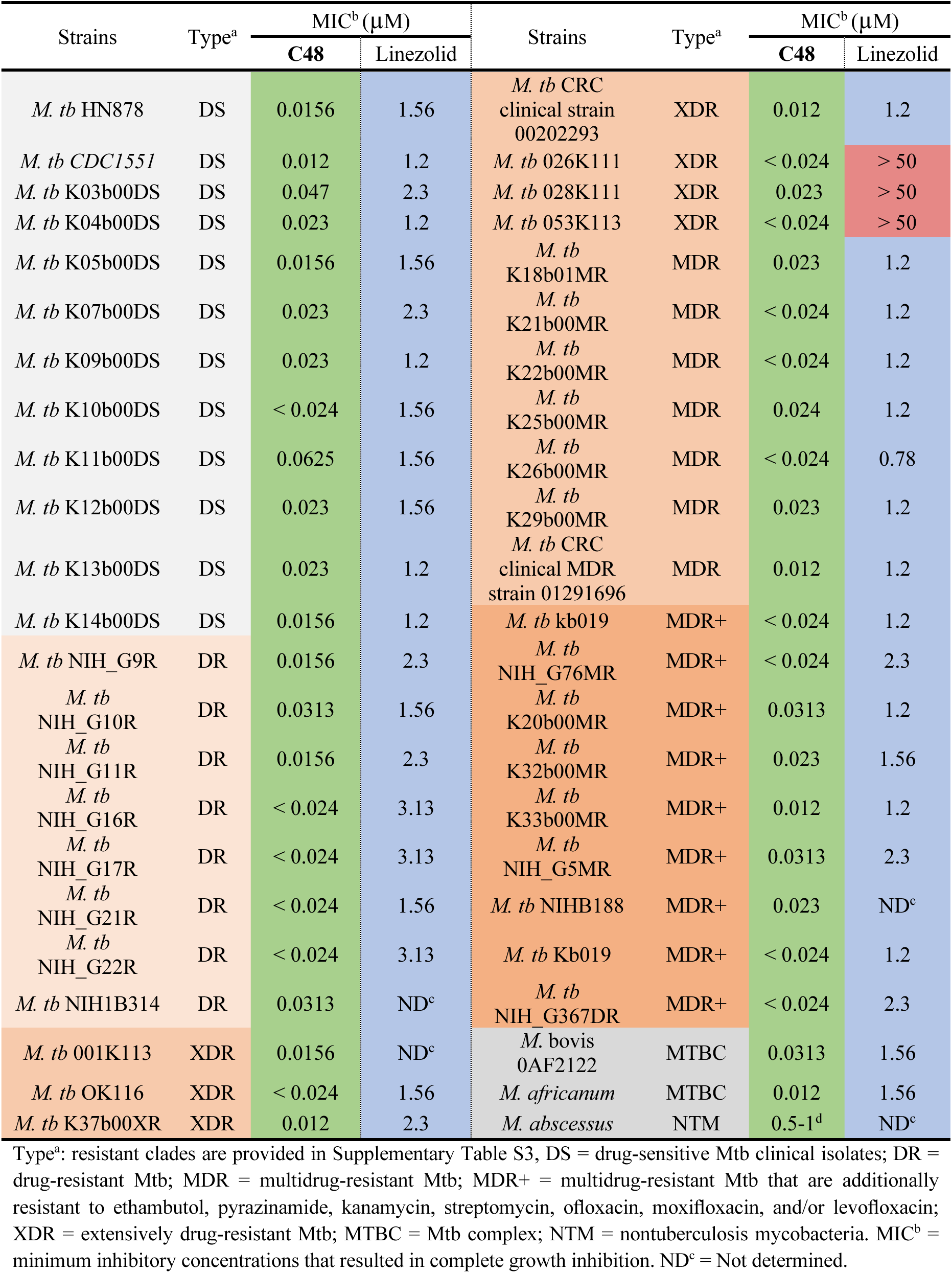
MIC screening of C48 against a panel of pathogens and strains.

| Strains | Type<sup>a</sup> | MIC<sup>b</sup> (μM) |  | Strains | Type<sup>a</sup> | MIC<sup>b</sup> (μM) |  |
| --- | --- | --- | --- | --- | --- | --- | --- |
|  |  | C48 | Linezolid |  |  | C48 | Linezolid |
| *M. tb* HN878 | DS | 0.0156 | 1.56 | *M. tb* CRC clinical strain 00202293 | XDR | 0.012 | 1.2 |
| *M. tb* CDC1551 | DS | 0.012 | 1.2 | *M. tb* 026K111 | XDR | < 0.024 | > 50 |
| *M. tb* K03b00DS | DS | 0.047 | 2.3 | *M. tb* 028K111 | XDR | 0.023 | > 50 |
| *M. tb* K04b00DS | DS | 0.023 | 1.2 | *M. tb* 053K113 | XDR | < 0.024 | > 50 |
| *M. tb* K05b00DS | DS | 0.0156 | 1.56 | *M. tb* K18b01MR | MDR | 0.023 | 1.2 |
| *M. tb* K07b00DS | DS | 0.023 | 2.3 | *M. tb* K21b00MR | MDR | < 0.024 | 1.2 |
| *M. tb* K09b00DS | DS | 0.023 | 1.2 | *M. tb* K22b00MR | MDR | < 0.024 | 1.2 |
| *M. tb* K10b00DS | DS | < 0.024 | 1.56 | *M. tb* K25b00MR | MDR | 0.024 | 1.2 |
| *M. tb* K11b00DS | DS | 0.0625 | 1.56 | *M. tb* K26b00MR | MDR | < 0.024 | 0.78 |
| *M. tb* K12b00DS | DS | 0.023 | 1.56 | *M. tb* K29b00MR | MDR | 0.023 | 1.2 |
| *M. tb* K13b00DS | DS | 0.023 | 1.2 | *M. tb* CRC clinical MDR strain 01291696 | MDR | 0.012 | 1.2 |
| *M. tb* K14b00DS | DS | 0.0156 | 1.2 | *M. tb* kb019 | MDR+ | < 0.024 | 1.2 |
| *M. tb* NIH_G9R | DR | 0.0156 | 2.3 | *M. tb* NIH_G76MR | MDR+ | < 0.024 | 2.3 |
| *M. tb* NIH_G10R | DR | 0.0313 | 1.56 | *M. tb* K20b00MR | MDR+ | 0.0313 | 1.2 |
| *M. tb* NIH_G11R | DR | 0.0156 | 2.3 | *M. tb* K32b00MR | MDR+ | 0.023 | 1.56 |
| *M. tb* NIH_G16R | DR | < 0.024 | 3.13 | *M. tb* K33b00MR | MDR+ | 0.012 | 1.2 |
| *M. tb* NIH_G17R | DR | < 0.024 | 3.13 | *M. tb* NIH_G5MR | MDR+ | 0.0313 | 2.3 |
| *M. tb* NIH_G21R | DR | < 0.024 | 1.56 | *M. tb* NIHB188 | MDR+ | 0.023 | ND<sup>c</sup> |
| *M. tb* NIH_G22R | DR | < 0.024 | 3.13 | *M. tb* Kb019 | MDR+ | < 0.024 | 1.2 |
| *M. tb* NIH1B314 | DR | 0.0313 | ND<sup>c</sup> | *M. tb* NIH_G367DR | MDR+ | < 0.024 | 2.3 |
| *M. tb* 001K113 | XDR | 0.0156 | ND<sup>c</sup> | *M. bovis* 0AF2122 | MTBC | 0.0313 | 1.56 |
| *M. tb* OK116 | XDR | < 0.024 | 1.56 | *M. africanum* | MTBC | 0.012 | 1.56 |
| *M. tb* K37b00XR | XDR | 0.012 | 2.3 | *M. abscessus* | NTM | 0.5-1<sup>d</sup> | ND<sup>c</sup> |

On biotin-free agar plates supplemented with **C48** at four times the minimum inhibitory concentration (MIC), spontaneously resistant mutants emerged at a frequency of approximately 3×10^−8^. Whole-genome sequencing of 16 independently isolated mutants revealed that each harbored a single nonsynonymous mutation in *bioA*, leading to changes at either position Met91 (15 out of 16 clones) or Cys168 (1 out of 16 clones). These mutations conferred either medium-level resistance (Met91Val, Met91Thr, and Cys168Tyr) or high-level resistance (Met91Ile) to **C48 (**Supplementary Table S8). All selected mutants retained susceptibility to RIF and INH that were like that of the parent strain (Fig. 2F, 2G and 2H). Cys168 is juxtaposed between the 3-bromo and 4-fluoro substituents of **C48** in the P1 binding pocket while Met91 is adjacent to the carbonyl group of the azaindanone fragment in the P2 binding pocket. Both pockets are nearly fully occupied, explaining why mutations at Cys168 and Met91 result in a loss of potency. Notably, the bulky Met91I substitution likely strongly disrupts critical π-π stacking interactions with Trp64 and Trp65 in the P2 pocket, contributing to the observed high-level resistance. Collectively, these studies confirm that the antibacterial activity of **C48** is due to inhibition of BioA. Additionally, **C48** exhibited no cytotoxicity toward mammalian cell lines (HepG2 and HT-29) at concentrations up to 100 µM, resulting in a therapeutic index (IC_50_/MIC_50_) greater than 10,000 (Fig. 3A and Supplementary Fig. S4A). The pronounced susceptibility of NTM pathogens and both drug-sensitive and drug-resistant Mtb clinical strains to **C48**, combined with lack of activity against ESKAPE pathogens and mammalian cells, highlights the compounds selectivity and promising safety profile.

**Figure 3.**
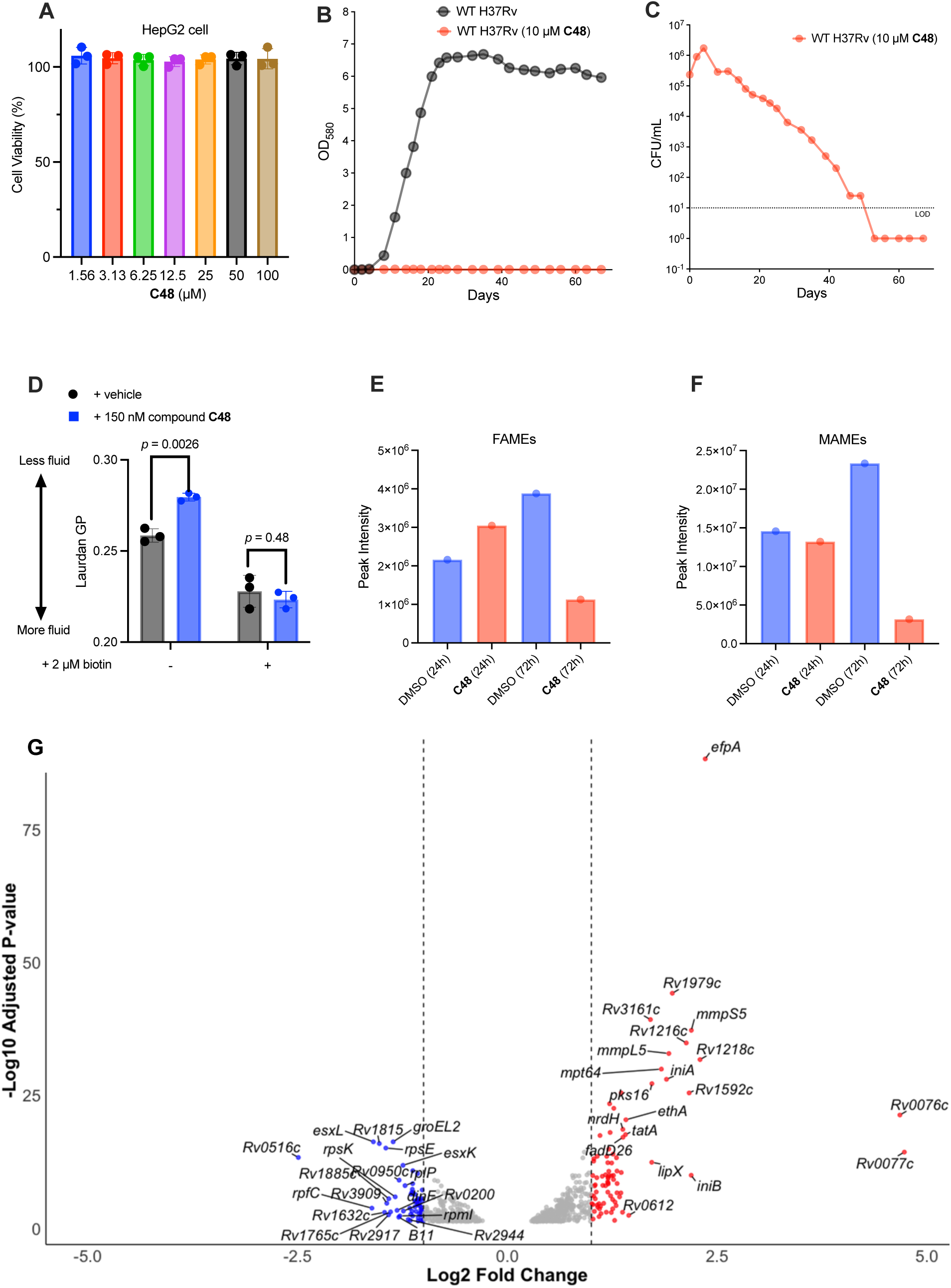
C48 recapitulates biotin starvation and induces cell envelope stress. (**A**) Viability of HepG2 after 48 h treatment with the indicated concentrations of **C48**. Values are normalized to vehicle treated condition (DMSO). The assay was performed as triplicate. (**B**) OD readings of **C48** kill kinetics, the assay was performed in GAST medium against wild-type (WT) *Mtb* (H37Rv) and the OD_580_ was taken at the indicated time. (**C**) Time to kill kinetics of **C48**, the assay was performed in GAST medium against wild-type (WT) *Mtb* (H37Rv) and the CFU was counted at the indicated time. (**D**) Mtb membrane fluidity in GAST medium upon treatment with **C48**. (**E**) and (**F**) Densitometry analysis of [^14^C]acetate-labeled FAMEs and MAMEs upon the exposure to **C48** (100 nM) or control (DMSO). (**G**) Transcription profile of **C48**. Differential expression of Mtb genes in response to **C48** exposure, with top 20 most up (red) and down (blue) regulated genes listed (*p*-value cutoff of 0.05). Vertical dashed lines representing log2-fold change cutoff of ± 1.

### C48 treatment induces cell envelope stress

To determine whether **C48** recapitulated the bactericidal phenotype of a *bioA* deletion mutant-which exhibits slow sterilization when starved for biotin^9^ - we incubated Mtb with 10 µM **C48** in a biotin-free medium. The growth of Mtb was arrested (Fig. 3B) and Mtb underwent approximately three doublings before CFU’s declined at a rate of one log_10_ CFU per week until the culture was sterilized, mirroring the phenotype observed for the Δ*bioA* mutant^5^ (Fig. 3C). Given biotin’s central role in regulating lipid metabolism and enabling bacterial membrane remodeling in response to environmental changes, we hypothesized that altered membrane fluidity could underlie the phenotype of Δ*bioA*^20, 21^. To test this, we quantified membrane fluidity with laurdan, a well-established fluorescent probe for studying membrane fluidity in bacteria. Laurdan generalized polarization (GP) was calculated as described in the methods, with blue fluorescence (λ_em_ = 440 nm) indicating ordered domains and red-shifted fluorescence (λ_em_ = 490 nm) reflecting disordered regions. Treatment of Mtb cultures with 150 nM **C48** resulted in an increase in laurdan GP, signifying enhanced membrane order, tighter lipid packing, and a shift away from unsaturated or branched fatty acids (Fig. 3D).^26^ In contrast, controls supplemented with biotin showed no significant change. The changes in membrane fluidity prompted us to investigate the lipids composition by [^14^C]acetate-labeled fatty acid methyl esters (FAMEs) and mycolic acid methyl esters (MAMEs). TLC analysis revealed that treatment with 100 nM **C48** reduced both FAMEs and MAMEs in a time-dependent manner (Fig. 3E and 3F). Notably, keto-mycolates – which are essential for pellicle growth and Mtb biofilm formation^27^ - were significantly decreased to negligible levels (Supplementary Fig. S4B) following **C48** exposure, highlighting the potential of biotin inhibitors for the treatment of chronic tuberculosis.

We further hypothesized that bactericidal activity caused by biotin depletion might result from downstream perturbations in lipid metabolism and thus sought to investigate the transcriptional response. RNA sequencing identified changes in mRNA levels for 154 genes upon **C48** exposure. As expected for a cell-wall targeting antibiotic^28, 29^ genes involved in cell envelope synthesis were among those most upregulated (Fig. 3G). Notably, enzymes essential for mycolic acid and surface lipid (PDIM) production-including polyketide synthases (*pks3*, *pks16*) and *fadD26*-were upregulated. **C48** also induced the expression of *iniA* and *iniB*, genes known to be activated by the cell wall-targeting INH^30^. Additionally, we observed the increased expression of several efflux pumps, particularly the siderophore export systems (mmpL5/mmpS5). This upregulation may indicate disruption of siderophore recycling in response to **C48** treatment^31^.

### Pharmacokinetic profile and initial efficacy study of C48 in conventional mice model

To determine its potential to reach therapeutically relevant concentrations in vivo, we assessed the pharmacokinetic properties of **C48** in CD-1 mice. **C48** exhibited excellent oral bioavailability (*F* > 100%) and a half-life of 0.91 hours. **C48**’s improved oral exposure resulted in an AUC (p.o.)/MIC value of 1570, which represents a 39,000-fold improvement compared to hit **6** (Fig. 4A and Supplementary Table S1). In a dose-escalation study conducted in BALB/c, **C48** demonstrated linear oral absorption up to 200 mg/kg and was well tolerated, with no adverse effects observed (Fig. 4B). Distribution studies confirmed that **C48** is efficiently delivered into the lungs throughout the observation period (Fig. 4C), despite **C48** demonstrated low volume of distribution. These findings supported advancing **C48** to in vivo efficacy studies.

**Figure 4.**
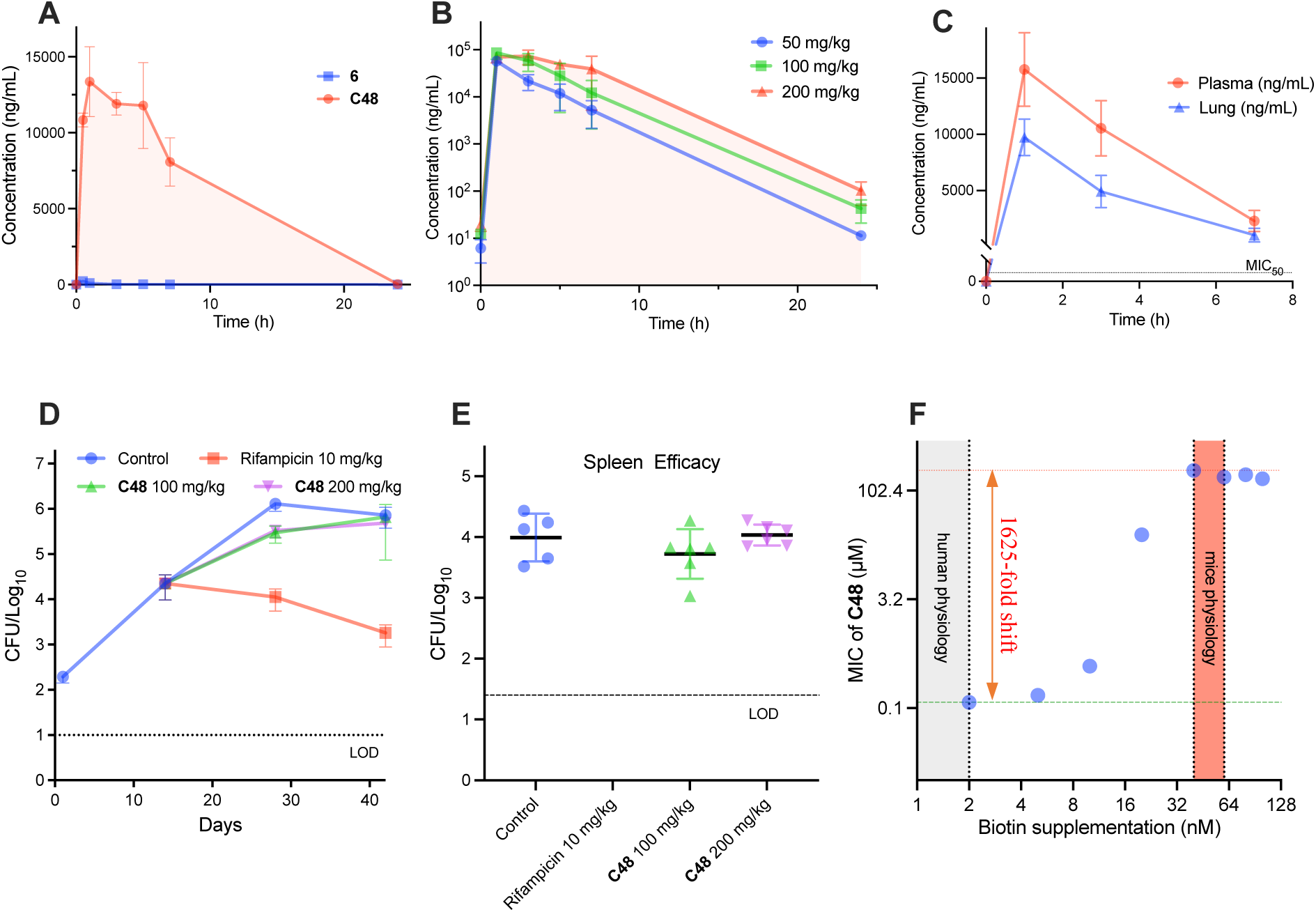
Pharmacokinetic and initial efficacy study of C48 in conventional mouse model. (**A**) Oral pharmacokinetic profile of **C48** and **6** in CD-1 mice (25 mg/kg, n = 3). (**B**) **C48** dose escalation and tolerability study in CD-1 mice (n = 3) at 50 mg/kg, 100 mg/kg and 200 mg/kg administered orally for 4 days. Plasma concentrations were measured by HPLC coupled with tandem mass spectrometry (LC-MS/MS). Black dotted line: MIC_50_ or minimum concentration that inhibits 50% growth. (**C**) Tissue distribution of **C48** in CD-1 mice following oral administration of 25 mg/kg (n = 3). Lung and plasma concentrations were quantified by LC-MS/MS. (**D**) and (**E**) **C48** was evaluated in a standard model of acute TB infection in BALB/c mice. The mice (n = 5 to 6) were infected on day 0, drugs were administrated 14 days post infection via oral gavage 7 days/week, and the treatment lasted for 28 days. Lung data was collected and CFU were enumerated on Day 28 and 42. Rifampicin treatment group in **E** is below the LOD. LOD = limit of detection. At Day 28 100 mg/kg **C48** showed a *p* < 0.0001 and 200 mg/kg **C48** showed a *p* = 0.0001. (**F**) The potency of **C48** against WT Mtb was antagonized in the presence of elevated biotin. The MIC against WT Mtb was performed and the MIC_50_ was plot in the function of biotin supplementation.

The initial efficacy study of **C48** was conducted in BALB/c mice, representing a conventional TB mouse model. Following aerosol infection with ∼10² CFU of Mtb H37Rv, mice received a 28-day oral treatment of **C48** or Rifampicin, and the CFU of lung and spleen were enumerated on Day 28 and 42. However, **C48** failed to display efficacy in this conventional mouse model of TB infection (Fig. 4D and 4E) showing no significant difference between the **C48** treated and vehicle control mice at the ending point. This lack of efficacy was likely due to high endogenous biotin levels in mice, as mouse plasma contains approximately 40 nM biotin, approximately 40 times the concentration found in humans^15, 32^. Consistent with this hypothesis, in vitro analysis confirmed that **C48**’s anti-TB activity is biotin-dependent. While **C48** is potent in a biotin environment mimicking human physiology (MIC_50_ = 1.2 µM), its efficacy was severely compromised under mouse-level biotin conditions (MIC_50_ = 195 µM), with activity shifting 1625-fold to a negligible level (Fig. 4F and Supplementary Fig. S4C).

### Establishing a biotin-deficient mouse model for TB

Brown *et al*. established a transient mouse model of biotin deficiency through single intravenous streptavidin injection, which reduces biotin levels for ∼12 hours^15^. This short window complicates studying chronic infections by slow-growing bacteria such as Mtb, where animal study requires monitoring over several weeks. Consequently, there is an unmet need for an easy-to-operate, sustained mouse model that recapitulates human biotin physiology for extended durations. To address this, we developed a low biotin mouse model by formulating mouse chow with egg whites containing avidin to deplete biotin during food taken. In this model, the serum biotin levels were reduced from 40 to 1 nM in 24 h and maintained these low, steady-state biotin levels during the time frame tested (Fig. 5A)^33, 34^. We next sought to determine if this new biotin-deficient model could be challenged by WT Mtb, and, as expected, this reduction in biotin did not affect TB infection and the growth of Mtb is the same as those in a conventional mice TB model (Fig. 5B and 5C).

**Figure 5.**
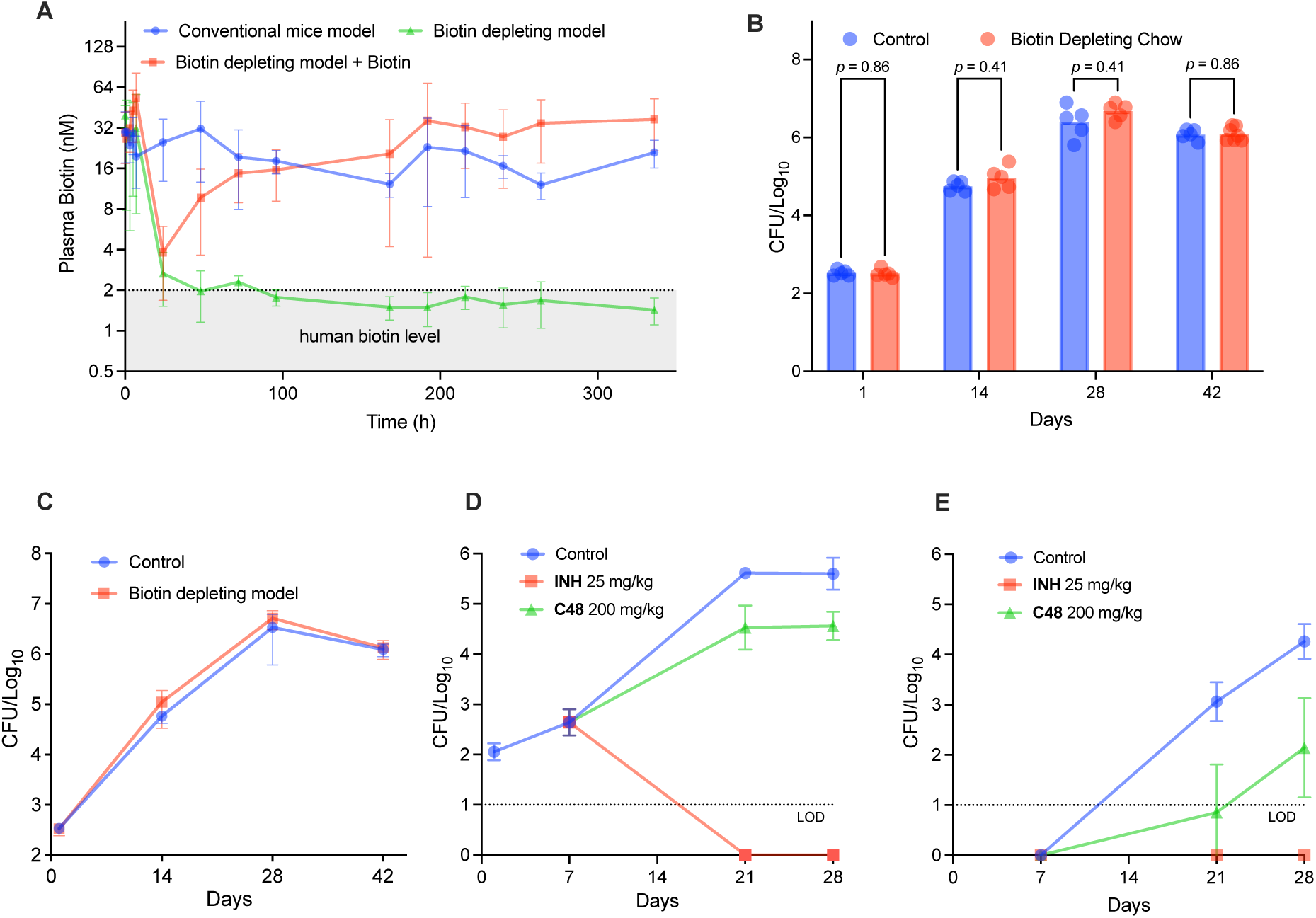
In vivo efficacy of C48 in biotin-deficient mouse model. (**A**) Quantification of biotin levels in mice plasma following different mice chow administrated, Conventional mice model : regular mice chow with biotin supplemented (PicoLab^®^ Rodent Diet 20), Biotin depletion model (biotin-free, Inotiv TD.81079). 6 mice were tracked for biotin levels for 2 weeks and due to IACUC guidelines, timepoints at 1, 3, 5, 7, 48, 72 and 96 h contained 3 mice samples and the rest timepoints contained 6 mice samples. (**B**) and (**C**) Biotin depletion has no effect on the ability of WT Mtb to cause infection in vivo. Mice (n = 5) were infected with WT Mtb under regular chow and biotin depletion chow. The lungs were collected, homogenized and the CFU was counted at the indicated time. (**D**) and (**E**) **C48** was evaluated in a biotin-deficient model of acute TB infection in BALB/c mice. The mice (n = 5 to 7) were infected on day 0, **C48** was administrated 7 days post infection via oral gavage 7 days/week, and the treatment lasted for 21 days. Lung and spleen samples were collected and CFU were enumerated on Day 28. LOD = limit of detection. All comparisons showed *p* < 0.0001 except for the lung CFU at Day 28 which showed a *p* = 0.0003.

Lastly, we conducted efficacy studies in this model, BALB/c mice were placed on the biotin-depletion diet seven days prior to infection, resulting in serum biotin levels of 1-2 nM. Mice were then aerosol-infected with 10² CFU of Mtb H37Rv. Treatment was initiated one week after infection with once-daily oral dosing of **C48** (at 200 mg/kg) or isoniazid (INH, 25 mg/kg) and continued for three weeks. **C48**-treated mice demonstrated significantly reduced bacterial burden in lungs (*p* = 0.0003) and spleen (*p* < 0.0001) by 1.0 log_10_ CFU and 2.1 log_10_ CFU, respectively (Fig. 5D and 5E). The activity of **C48** in lungs is thus consistent with the in vitro time-kill studies showing a one log_10_ CFU decrease over three weeks. The robust efficacy observed in this new low-biotin mouse model underscores the critical role of an animal model that more accurately mirrors human physiology and these results thus provide proof-of-concept for the therapeutic potential of inhibiting the biotin synthesis pathway of Mtb.

### C48 shows strong synergy with inhibition of BioB

Combination therapy is a cornerstone for TB treatment, particularly, regimens with synergistic effects may shorten treatment duration, lower relapse rates, and prevent the emergence of drug resistance. We therefore explored the potential synergy resulting from simultaneous inhibition of BioA and BioB, which mediated adjacent steps in biotin synthesis. For a BioB inhibitor, we developed acidomycin-amide, a potent synthetic prodrug (MIC_50_ = 0.5 µM) of the natural product Acidomycin^9, 35^, by introducing a neutral amide to mask its charged carboxylic acid group (Detailed synthetic procedures and compound characterizations are provided in the Supplementary Materials and Supplementary Scheme S1). We confirmed its on-target activity by showing that a BioB-overexpressing Mtb strain was significantly less susceptible to acidomycin-amide (Fig. 6A) but not to isoniazid (Fig. 6B). We next tested **C48** in combination with acidomycin-amide in vitro by determining the checkerboard fractional inhibitory concentration index (FIC_i_) wherein FIC_i_ ≤ 0.5, 0.5 < FIC_i_ < 1, and FIC_i_ >1 indicate synergistic, additive, and antagonistic interactions, respectively. **C48** and acidomycin-amide demonstrated strong synergy with an FIC_i_ of 0.28 (Fig. 6C). Given this pronounced synergy, we evaluated the combination’s bactericidal activity starting from sublethal concentrations. Individually, **C48** and acidomycin-amide showed no bactericidal activity at 1.5-, 3-, or 6-fold the MIC. In contrast, the combination displayed superior killing with nearly 2-log reduction at 21 days and 4-log reduction at 42 days at 1.5-, 3- or 6-fold MIC concentrations (Fig. 6D-F). We interpret the interaction between **C48** and acidomycin-amide (Fig. 6C) is derived from mutual potentiation wherein **C48** inhibits formation of DTB enhancing the interaction of acidomycin-amide with its target BioB while acidomycin-amide potentiates **C48**’s activity by depleting BioA’s substrate KAPA through inhibition of the precursor pimelate (Supplementary Fig S1), similar to the mechanism of mutual potentiation reported for trimethoprim and sulfamethoxazole^36^. The potent synergy between BioA and BioB may indicates a previously unrecognized metabolic feedback loop in the biotin pathway and the cyclic mutual potentiation of BioA and BioB amplifies depletion of biotin.

**Figure 6.**
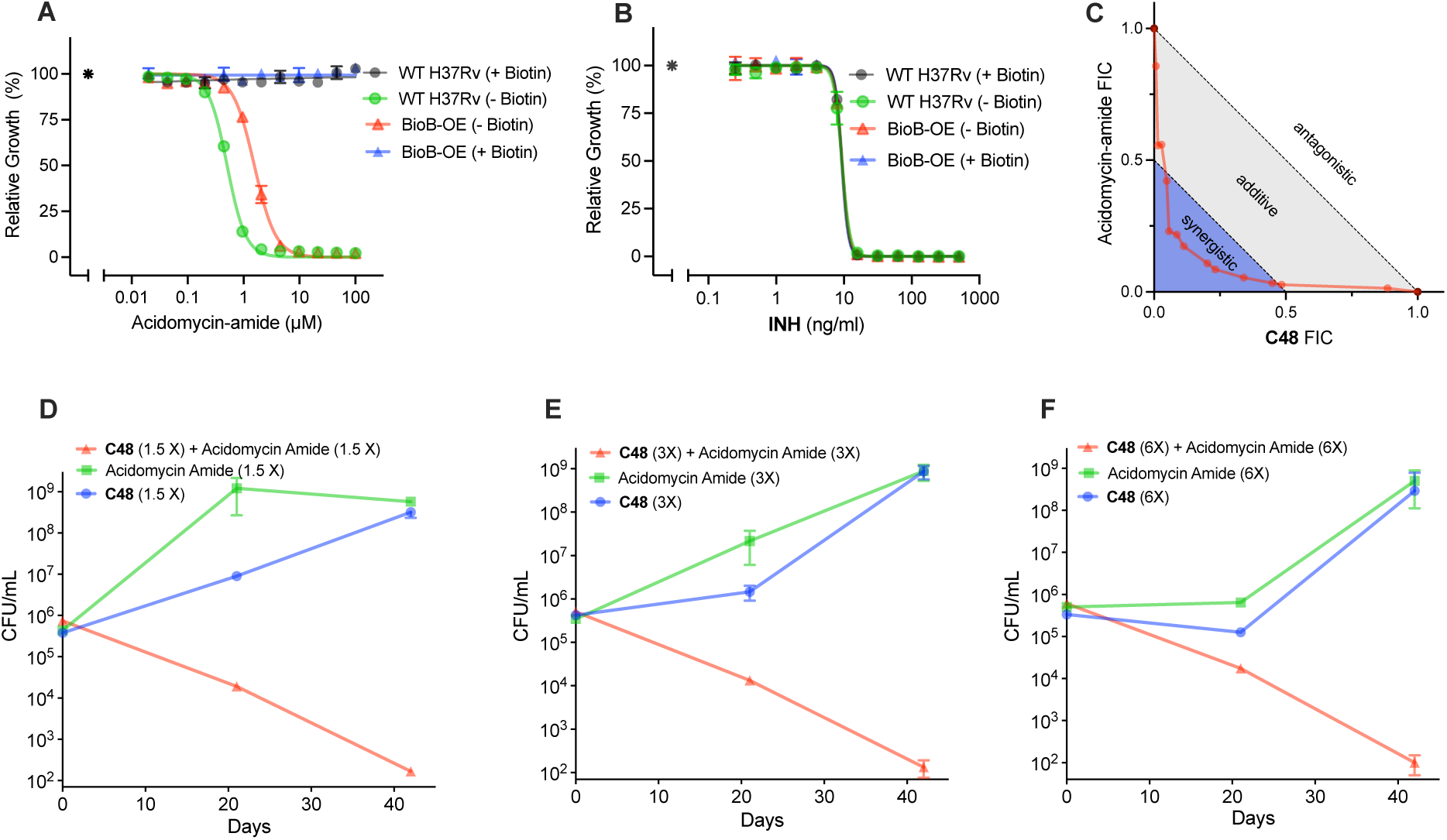
C48 and Acidomycin-amide synergy study. (**A**) Sensitivity of Acidomycin-amide against Mtb was subjected to the expression level of BioB, BioB-OE = BioB protein over-expressed strain compared to WT (H37Rv). (**B**) The susceptibility of Isoniazid remains unchanged against Mtb BioB mutant. (**C**) Chequerboard assays of **C48** and acidomycin-amide against Mtb, FIC_i_ = FIC**_C48_** + FIC_acidomycin-amide_. (**D**), (**E**) and (**F**) Time to kill of **C48** and acidomycin-amide, the assay was performed in GAST medium against WT Mtb, and the CFU was counted at the indicated time. 1.5, 3 and 6X represent 1.5, 3 and 6-fold MIC_50_. MIC experiments were performed at least twice.

## DISCUSSION

It is a challenge for target-based and phenotypic drug discovery approaches to recreate the conditions pathogens encounter in humans to evaluated small-molecules in vitro and, especially, in animal models. The standard culture media, optimized to maximize growth, often induce adaptation in bacterial pathogens that are be fundamentally different from those required to flourish in the nutrient-limited, host-specific environments they encounter in the body^12^. For example, Dick et al reported a lead compound displayed strong in vitro activity but fail to show efficacy in vivo, and the discrepancy was traced to major differences in carbon metabolism between bacteria growing in standard laboratory broth and those replicating in infected lung tissue^37^. Increasing evidence shows that the host relevant environment can profoundly influence antibiotics potency and a number of new inhibitors have been identified in clinic relevant, nutrient-limited conditions that better mimic host-like conditions^12^. This issue extends beyond tuberculosis drug discovery as recent work demonstrating nutrient transport as anti-malaria drug target in physiological relevance conditions^38^.

Biotin is an essential cofactor for Mtb, however, the development of an orally bioavailable lead compound targeting biotin biosynthesis remains elusive. In this report, **C48** was developed as a potent inhibitor targeting BioA, exhibiting a *K*i of 200 pM (Fig. 1A and 2C). Its potent biochemical activity translated into robust whole cell activity with MICs ranging from 0.012–0.093 µM against a panel of 70 Mtb strains, including drug-resistant isolates (Table 1 and Supplementary Table S5). Several lines of evidence support that **C48**’s impact on Mtb is due to inhibition of BioA: (i) **C48** is antagonized in a concentration-dependent manner by biotin, DTB and DAPA, but not by KAPA (Fig. 2D); (ii) potency of **C48** decreases monotonically with increasing expression of BioA (Fig. 2E); (iii) all spontaneously **C48** resistant mutants analyzed by whole-genome sequencing carried mutations in *bioA* (Supplementary Table S8).

Structural studies revealed a remarkable array of parallel-displaced, sandwich, and T-shaped π-interactions in the cocrystal structure of **C48** bound to BioA (Fig. 1C). These interactions elucidate the molecular basis for **C48**’s high binding affinity, explain the resistance of the isolated mutants, and provide insight into the lack of activity against organisms with divergent BioA orthologs. **C48** possesses excellent bioavailability (*F* > 100%) and dramatically improved oral exposure (Fig. 4A and Supplementary Table S1). Conventional mouse models are confounded by elevated biotin levels compared to the biotin levels present in humans^15^. To overcome this limitation, we developed an easy-to-operate, low-biotin mouse model for TB that recapitulates human biotin concentrations over the sustained duration of time needed to study slow growing bacteria like Mtb (Fig. 5A). In this model, oral administration of **C48** resulted in significant suppression of Mtb growth in vivo, with reductions of 1.0 log₁₀ CFU in the lungs and 2.1 log₁₀ CFU in the spleen (Fig 5D and 5E). Collectively, these findings establish in vivo proof-of-concept for targeting biotin biosynthesis as a therapeutic strategy against infections with Mtb.

We next explored the combination potential of biotin inhibitors. Similar to co-trimoxazole (trimethoprim and sulfamethoxazole, Bactrim^®^), inhibitors of BioA and BioB are targeting adjacent steps in cofactor biotin biosynthesis. Accordingly, **C48** and acidomycin-amide demonstrated strong synergy with an FIC_i_ of 0.28 (Fig. 6C) and pronounced bactericidal activity (Fig. 6D, E and F). Co-trimoxazole’s potency does not stem from “sequential blockade of a major biosynthetic pathway” as initially proposed, but rather arises from mutual potentiation. Sulfamethoxazole potentiates the activity of trimethoprim by depleting dihydrofolate, a competitive substrate, and trimethoprim potentiates sulfamethoxazole activity by depleting dihydropteroate pyrophosphate through inhibition of a folate-dependent precursor^36^. One interpretation for the synergy between **C48** and acidomycin-amide is that is also cause by mutual potentiation wherein **C48** potentiates acidomycin-amide by limiting downstream production of DTB, which is the substrate of BioB. However, this alone may not be sufficient to explain the observed mutual potentiation. It may instead require a metabolic feedback loop in the biotin pathway, in which BioB inhibition also constrains KAPA production, and this previously unrecognized metabolic feedback loop in the biotin pathway amplifies depletion of biotin.

Another unique feature of biotin depletion is the impact to Mtb cell envelope lipids, and the lipid composition of membranes is crucial for maintaining fluidity and optimal lipid-protein interactions, such as MmpL4/S4 and MmpL5/S5^26, 39^, and disruption of biotin biosynthesis inhibition triggers extensive lipid remodeling, profoundly altering membrane properties^26^. Indeed, we observed a significant decrease in membrane fluidity, as measured by an increased laurdan GP that was fully antagonized by biotin supplementation (Fig. 3D). This finding aligns with recent work by Sullivan and colleagues, who demonstrated that *Mycobacterium abscessus* requires biotin to support lipid remodeling and maintain membrane fluidity, particularly during adaptation to alkaline lung environments^26^. Inhibition of biotin biosynthesis selectively depleted both branched and unsaturated fatty acids, resulting in decreased membrane fluidity. Notably, tuberculostearic acid (10*R*-methylstearic acid), the most abundant mycobacterial fatty acid, was also significantly affected. Tuberculostearic acid plays a pivotal role in regulating this membrane heterogeneity, and its rapid depletion may disproportionately disrupt the function of membrane-associated proteins^40^. These findings underscore the need for further research to elucidate the organization and function of the mycobacterial inner membrane and its associated transporters. Evidence from this study and that of Sullivan *et al*. suggests that the strong, bactericidal impact biotin antimetabolites have on mycobacteria is ultimately caused by a breakdown of the inner membrane^26^. Complementary studies by Brown *et al*. have further shown that inhibition of biotin biosynthesis depletes phosphatidylethanolamines found in lipopolysaccharide (LPS), contributing to compromised cell envelope function^41^.

Together, this work presents a potent, oral effective lead targeting biotin biosynthesis - a conditionally essential pathway in *Mycobacterium tuberculosis* in a host-relevant model. Future studies should aim to realize the untapped potential of conditionally essential targets, using screening strategies in media or animal models that better mimic physiological conditions, thereby offering a new approach to discovering antibiotics with novel mechanism of action.

## Methods

All animal studies were ethically reviewed and approved (Approval Number: AUP-269.033) by the Institutional Animal Care and Use Committee of the Center for Discovery and Innovation, Hackensack Meridian Health.

### Bacterial strains, media and culture conditions

A complete list of bacterial strains and plasmids used in the study can be found in Supplementary Table S3. Mtb H37Rv was grown in GAST, 7H9, modified 7H9 medium, on 7H10 or 7H11 agar plates: GAST medium (0.3 g/L Bacto Casitone, 2 g/L ammonium chloride, 1 g/L L-alanine, 4 g/L dibasic potassium phosphate, 2 g/L citric acid, 50 mg/L ferric ammonium citrate, 1.2 g/L magnesium chloride hexahydrate, 0.6 g/L potassium sulfate, 1% [v/v] glycerol, 0.05% [v/v] Tyloxapol, pH 6.6). 7H9 medium (Middlebrook 7H9 Broth supplemented with 0.2% [v/v] glycerol, 0.05% [v/v] tyloxapol and 10% [v/v] ADNaCl (0.5% [w/v] BSA, 0.2% [w/v] dextrose, 0.085% [w/v] sodium chloride). Modified 7H9 medium without biotin (m7H9: 0.5 g/L ammonium sulfate, 0.5 g/L L-Glutamic acid, 0.1 g/L sodium citrate, 2.5 g/L disodium phosphate, 1.0 g/L monopotassium phosphate, 0.04 g/L ferric ammonium citrate, 0.05 g/L magnesium sulfate, 1.0 mg/L pyridoxine, 0.5 mg/L calcium chloride, 1.0 mg/L zinc sulfate, 1.0 mg/L copper sulfate) supplemented with 0.2% [v/v] glycerol, 0.05% [v/v] tyloxapol and 10% [v/v] ADNaCl. 7H10 agar [Middlebrook 7H10 supplemented with 10% OADC (Becton Dickinson) and 0.5% glycerol]. 7H11 agar [10.5 g 7H11 powder (Millipore Sigma) and 5 mL of 50% glycerol in 445 mL of Millipore water, 50 mL OADC (Fisher) was supplemented]. When required, hygromycin B, zeocin, or kanamycin was used at a concentration of 50 µg/ml, 25 µg/ml or 25 µg/ml. Preparation of competent cells, electroporations, and genomic DNA isolations were performed as described^42^. Where indicated, biotin (Sigma) was added at a concentration of 2 µM. All drug resistant Mtb isolates were grown in Dubos-based medium (6.5 g/L Dubos broth base, 0.81 g/L NaCl, 7.5 g/L glucose, and 5 g/L BSA fraction V).

### BioA protein expression and purification

The recombinant BioA protein was overexpressed in *Escherichia coli* and purified with Ni affinity and size exclusion chromatography as described previously^43, 44^. Codon-optimized *bioA* gene with a cleavable N-terminal His tag (Integrated DNA technology, Coralville, IA) was inserted into pET32 plasmid. *E. coli* BL-21 (DE3) cells (New England BioLabs) were transformed with the BioA construct. The cells were grown at 37 °C in Luria Broth (LB) medium containing 100 μg/mL ampicillin (Gold Biotechnology). After the culture reached an OD_600_ = 0.6, IPTG (Gold Biotechnology) was added to induce gene expression and the culture was incubated at 16 °C for 16 h. Cells were harvested by centrifugation at 3700 x g for 25 min and resuspended in lysis buffer containing 50 mM HEPES pH 7.5, 500 mM NaCl, and 400 μM PLP. The recombinant BioA protein was purified according to the procedure published by Dai *et al*., with some modifications^11^. The cells were incubated with hen egg white lysozyme (Hampton Research) and DNase I (Roche Applied Sciences) for 30 min followed by sonication (Sonicator 3000, Misonix) to enhance cell lysis. The crude cell lysate was clarified by centrifugation at 13,000 x *g* for 40 min (Fixed-angle rotor, 5810-R Centrifuge, Eppendorf). The clarified lysate was loaded onto a HiTrap TALON crude cobalt column (Cytiva) that had been equilibrated with a 50 mM HEPES buffer at pH 7.5 with 500 mM NaCl, and 0.1 mM TCEP. After washing with 15 column volumes of the same buffer, protein was eluted using 50 mM HEPES buffer at pH 7.5 with 500 mM NaCl, 0.1 mM TCEP, and 150 mM imidazole. The N-terminal poly-histidine tag was cleaved by adding recombinant human rhinovirus 3C protease to the eluted protein sample, followed by overnight dialysis at 4 °C. Application of the dialyzed sample to the cobalt column removed the cleaved poly-histidine tag and the protease. The purified protein was then applied to a Superdex 200 Increase 10/300 (Cytiva) size exclusion column using 50 mM HEPES pH 7.5, 50 mM NaCl, 0.5 mM TCEP as the mobile phase. An additional 1 mM PLP was added to the fractions containing BioA, and free PLP was removed by three rounds of ultrafiltration. Differential Scanning Fluorometry was used to ensure the protein was fully saturated with PLP as previously described^44^.

### Biochemical characterization

Compounds in 500 mM DMSO stock solution were robotically dispensed from Labcyte low dead volume 384-well source plates into Corning flat bottom black 96-well plates (Cat. 3991) using an Echo 550 liquid handler to achieve 8-point, 3-fold dilutions with concentrations ranging from 5 μM to 2.28 nM. The first and last columns contained positive (compound **6**)^20^ and negative (DMSO only) controls. Next, 50 µL of 2X reaction mixture (100 nM BioA, 640 nM BioD, 2 mM S-adenosyl methionine, 40 nM Fluorescent-DTB tracer, 70 nM streptavidin, 10 mM ATP, 50 mM NaHCO_3_, 1 mM MgCl_2_, 0.1 mM PLP, 0.0025% Igepal CA630, and 100 mM Bicine [pH 8.6]), containing all reaction components except KAPA, was dispensed into all wells of the plate. The reaction was started by the addition of 50 µL of freshly prepared 2X KAPA initiation solution (6 µM KAPA, 50 mM NaHCO_3_, 1 mM MgCl_2_, 0.1 mM PLP, 0.0025% Igepal CA630, and 100 mM Bicine [pH 8.6]) and incubated at 25 ℃ for 30 min. The reaction was terminated by the addition of 20 µL of 500 mM EDTA into each well of the assay plate. After 5 min equilibration, the plate was read on a CLARIO star microplate reader for fluorescence intensity with an excitation of 485 nM, an emission of 530 nM, and a cutoff of 530 nM. The data points were collected in triplicate and the averaged value was used to generate concentration-response plots. The IC_50_ value for each compound was obtained by nonlinear regression curve fitting of a four-parameter variable slope equation to the dose-response data using Prism software (9.5.1). The IC_50_ of best compound **C48** was manually repeated. 50 µL of assay buffer containing 3X **C48** was added to a 96-well plate to achieve 15-point, 1.5-fold dilutions with final concentrations ranging from 1 μM to 3.4 nM. The first and last columns contained positive (compound **6**)^21^ and negative (DMSO only) controls. Next, 50 µL of 3X reaction mixture (150 nM BioA, 960 nM BioD, 3 mM S-adenosyl methionine, 60 nM Fluorescent-DTB tracer, 105 nM streptavidin, 15 mM ATP, 50 mM NaHCO_3_, 1 mM MgCl_2_, 0.1 mM PLP, 0.0025% Igepal CA630, and 100 mM Bicine [pH 8.6]), containing all reaction components except KAPA, was dispensed into all wells of the plate. The reaction was started by the addition of 50 µL of freshly prepared 3X KAPA initiation solution (9 µM KAPA, 50 mM NaHCO_3_, 1 mM MgCl_2_, 0.1 mM PLP, 0.0025% Igepal CA630, and 100 mM Bicine [pH 8.6]) and incubated at 25 ℃ for 30 min. The reaction was terminated by the addition of 20 µL of 500 mM EDTA into each well of the assay plate. After 5 min equilibration, the plate was read on a CLARIO star microplate reader for fluorescence intensity with an excitation of 485 nM, an emission of 530 nM, and a cutoff of 530 nM. The assay was performed in triplicate. The *Ki* was further determined due to the tight binding nature of **C48**. 50 µL of assay buffer containing 3X **C48** was added to a 96-well plate to achieve 13-point dilution series with final concentrations 30, 20, 13.33, 11.11, 8.89, 7.41, 5.93, 3.95, 2.63, 1.75, 1.17, 0.78 and 0.52 nM. The first and last columns contained positive (EDTA) and negative (DMSO only) controls. Next, 50 µL of 3X reaction mixture (15 nM BioA, 150 nM BioD, 20 mM S-adenosyl methionine, 15 nM Fluorescent-DTB tracer, 26.3 nM streptavidin, 1.5 mM ATP, 50 mM NaHCO_3_, 1 mM MgCl_2_, 0.1 mM PLP, 0.0025% Igepal CA630, and 100 mM Bicine [pH 8.6]), containing all reaction components except KAPA, was dispensed into all wells of the plate. The reaction was started by the addition of 50 µL of freshly prepared 3X KAPA initiation solution (30 µM KAPA, 50 mM NaHCO_3_, 1 mM MgCl_2_, 0.1 mM PLP, 0.0025% Igepal CA630, and 100 mM Bicine [pH 8.6]). The plate was shaken for 5 min and incubated at 25 ℃ for 10 min, and the plate was read by kinetics model on a CLARIO star microplate reader for fluorescence intensity with an excitation of 485 nM, an emission of 530 nM, and a cutoff of 530 nM. The assay was performed in triplicate (n=3) and the normalized initial velocities (vi/v0) were fit by nonlinear regression analysis to the Morrison equation using Prism 9.5.1 to determine the *Ki* value.

### Co-Crystal X-ray structure of the BioA/C48 complex

BioA crystallization experiments were performed as previously described^11, 45^. BioA was concentrated to 10 mg/mL and co-crystallized with 1 mM of **C48** using the vapor diffusion hanging drop method. Protein crystals were obtained using microseeding and incubation against a reservoir solution containing 100 mM HEPES buffer pH 7.5, 100 mM MgCl_2_, and 9% w/v PEG 8000. Protein, reservoir, and seed solution were mixed in a 4:3:1 ratio. The crystals were cryoprotected with a solution containing 100 mM HEPES pH 7.5, 100 mM MgCl_2_, 15% PEG 400, and 15% PEG 8000 and immediately flash-cooled with liquid nitrogen. The X-ray diffraction data set was obtained at MX beamtime at a Diamond Light source (Oxfordshire, OX11 0DE), and the data set was processed using xia2-dials^46, 47^. Automated molecular replacement was performed using MrBUMP^48^. The structure was further refined in PHENIX, and manual model building using COOT was performed between the refinements^49, 50^. The restraints for the **C48** compound and PLP were obtained by eLBOW and geometries were further refined by REEL^51, 52^. The ligand was fitted to the difference density with COOT and refined by PHENIX. The quality of the model was validated using Molprobity^53^.

### Mtb Whole-Cell MIC assay

MIC measurements were based on a method using *bioA* mutants as previously described^20^. Wild-type (WT) H37Rv, BioA-UE, and BioA-OE were cultured in 10 mL of GAST medium + 2 µM biotin (with Hygromycin at 50 µg/mL and Kanamycin at 25 µg/mL for the *bioA* strains) in 25-cm^2^ tissue culture flasks with vented caps for approx. 7 d at 37 °C and 5% CO_2_ in a humidified incubator, growing to OD_580_ = ∼1.0. The cultures were then washed twice with 10 mL GAST without biotin and were diluted to OD_580_ = 0.005, +/- 2 µM biotin for WT H37Rv, and +200 ng/mL Anhydrotetracycline (ATc) but without biotin for the *bioA* strains. Compounds were solubilized in DMSO and dispensed into black, clear-bottom 384-well tissue culture plates using an HP D300e Digital Dispenser as 13-point, logarithmically distributed titrations from 0.01-100 µM in triplicate. 50 μL of OD_580_ = 0.005 suspension was pipetted using an Integra 16 Channel VIAFLO Electronic Pipette into each well, and plates were incubated for 14 days at 37 °C in the same conditions as above in stacks of no more than six plates wrapped with aluminum foil. Final OD_580_ values were normalized from 0–100% to the averages of no-drug (100%) and no-growth (0%) control wells. MIC values were calculated by fitting the log(inhibitor) vs. response data in GraphPad Prism to a Gompertz model provided by GraphPad (https://www.graphpad.com/support/faqid/1365/), with bottom and span best-fit values constrained to 0 and 100, respectively. When the calculated MIC was higher than the highest dose tested, it was reported to be greater than that dose. All MIC measurements were made at least twice, in independent replicate experiments.

### Supplementation experiments in Mtb

8-keto-7-aminopelargonic acid (KAPA), diaminopelargonic acid (DAPA), desthiobiotin (DTB) and biotin supplementation assays were performed otherwise identical with the whole cell MIC assay protocol, but adding 50 µM KAPA, 50 µM DAPA, 50 µM DTB or 2 µM biotin into the GAST media.

### Agar MIC of C48

To determine the agar MIC of **C48**, wild-type (WT) H37Rv was cultured in 10 mL of 7H9 medium in a 25-cm^2^ tissue culture flask with a vented cap for approx. 7 days at 37 °C and 5% CO_2_ in a humidified incubator, growing to OD_580_ = ∼1.0. The culture was then washed twice with modified 7H9 medium without biotin. From this washed culture, OD_580_ was measured, and 1 mL of culture was diluted to OD_580_ = 0.5. The remaining culture was pelleted and adjusted to OD_580_ = 2.0. 100µL of each culture was added and spread in each well of a set of two 6-well plates, with each well containing 6mL of modified 7H9 -biotin agar (m7H9, 1.5% Bacto agar) supplemented with 0.5% [v/v] glycerol and 10% [v/v] OADC. For each culture, one plate consisted of a No-Drug control well, a No-Growth control well containing 20µM rifampicin and 4 increasing concentrations of **C48**. The second plate consisted of 6 additional increasing concentrations of **C48**. The range of **C48** concentrations covered from ∼0.5x-32x the liquid IC_50_ (0.041-2.6 µM). The plates were incubated at 37 °C for 21 days. The agar MIC value was chosen based on the lowest concentration of **C48** to yield no colonies and determined to be 0.8 µM.

### Frequency of resistance for C48

To determine the Frequency of Resistance for **C48**, WT H37Rv was cultured in 10 mL of 7H9 medium in a 25-cm^2^ tissue culture flask with a vented cap for approx. 7 days at 37 °C and 5% CO_2_ in a humidified incubator, growing to OD_580_ = ∼1.0. After 7 days, the culture was diluted to OD_580_ = 0.01 in 20 mL of 7H9 medium in a 75-cm^2^ tissue culture flask with a vented cap and cultured for 7 d at 37 °C and 5% CO_2_ in a humidified incubator, growing to OD580 = ∼1.0. The culture was then washed twice with modified 7H9 medium without biotin. From this culture, serial dilutions (10^-4^, 10^-5^, 10^-6^) were plated on drug-free 7H10 agar plates supplemented with 0.5% [v/v] glycerol and 10% [v/v] OADC, to determine the total CFU input. The remaining culture was used to plate varying amounts on modified 7H9 -biotin agar plates supplemented with 0.5% [v/v] glycerol, 10% [v/v] OADC and 3.2 µM **C48**, which is equivalent to 4x the determined agar MIC. The plates were incubated at 37 °C for 21 days. Frequency of resistance was calculated as the fraction of resistant mutants isolated over the number of bacteria plated. To validate resistance, 30 resistance mutants were inoculated into 7H9 medium in 24-well plates, 800 µL per well, and incubated for 10 days at 37 °C and 5% CO_2_ in a humidified incubator. After 10 days, these cultures were then passaged into 10 mL of 7H9 medium in 25-cm^2^ tissue culture flasks with vented caps and incubated for approx. 7 days at 37 °C and 5% CO_2_ in a humidified incubator to OD_580_ = ∼1.0. A small portion of each culture was washed twice in GAST medium without biotin and used to determine the MIC against **C48**, including WT H37Rv as a control, in the manner as previously described. All 30 resistant mutants displayed an approximately 300-fold increase in MIC compared to WT H37Rv. The remainder of each culture was used to extract genomic DNA and 2 of these samples were later subjected to whole-genome sequencing to check for mutations that would confer resistance to **C48**.

### Fluctuation assay for C48

To confirm the fraction of resistant mutants and generate additional resistant mutants for **C48**, a small-scale Fluctuation Assay was performed. WT H37Rv was cultured in 10 mL of 7H9 medium in a 25-cm^2^ tissue culture flask with a vented cap for approx. 7 days at 37 °C and 5% CO_2_ in a humidified incubator, growing to OD_580_ = ∼1.0. After 7 days, the culture was diluted to OD_580_ = 0.00001 in 60 mL of 7H9 medium, which was then split into 14 separate 25-cm^2^ tissue culture flasks with vented caps and cultured for 14 days at 37 °C and 5% CO2 in a humidified incubator, growing to OD_580_ = ∼1.5-1.6. Four of the cultures were used to plate serial dilutions (10^-5^, 10^-6^) on drug-free 7H10 agar plates supplemented with 0.5% [v/v] glycerol and 10% [v/v] OADC to determine the total CFU input. The remaining 10 cultures were washed twice with modified 7H9 medium without biotin. These were then pelleted and plated on modified 7H9 -biotin agar plates supplemented with 0.5% [v/v] glycerol, 10% [v/v] OADC and 3.2µM **C48**. The plates were incubated at 37 °C for 27 days. Frequency of resistance was calculated as the fraction of resistant mutants isolated over the number of bacteria plated. Additionally, the mutation rate given as the *mutation rate per cell per division corrected by the plating efficiency*, was determined using the calculator provided by the Laboratory of Computational and Quantitative Biology (LCQB) at Sorbonne Université (http://www.lcqb.upmc.fr/bzrates). To validate resistance, 5 resistant mutants per plate (50 total) were inoculated into 7H9 medium in 24-well plates, 800 µL per well, and incubated for 10 days at 37 °C and 5% CO_2_ in a humidified incubator. After 10 days, these cultures were then passaged into 10 mL of 7H9 medium in 25-cm^2^ tissue culture flasks with vented caps and incubated for approx. 7 days at 37 °C and 5% CO_2_ in a humidified incubator to OD_580_ = ∼1.0. A small portion of each culture was washed twice in GAST medium without biotin and used to determine the MIC against **C48**, including WT H37Rv as a control, in the manner as previously described. Among the 50 resistant mutants, we found a range of increased resistance from 12-fold to over 300-fold shift in MIC compared to WT H37Rv. The remainder of each culture was used to extract genomic DNA and 10 of these samples were later subjected to whole-genome sequencing to check for mutations that would confer resistance to **C48**.

### Time-kill kinetics for C48

To analyze the time-kill kinetics for **C48**, WT H37Rv was cultured in 10 mL of GAST medium supplemented with 2µM biotin in a 25-cm^2^ tissue culture flask with a vented cap for approx. 7 days at 37 °C and 5% CO_2_ in a humidified incubator, growing to OD_580_ = ∼1.0. After 7 days, the culture was washed twice with 10 mL GAST without biotin and subsequently diluted to OD_580_ = 0.001 in 40mL GAST. **C48** was added to the culture at a concentration of 10µM, approximately 100x the liquid IC_50_, and the culture was split into separate 75-cm^2^ tissue culture flasks with vented caps. The cultures were incubated for upwards of 67 days at 37 °C and 5% CO_2_ in a humidified incubator. Killing was monitored every 3-4 days by plating serial dilutions on 7H10 agar plates supplemented with 0.5% [v/v] glycerol and 10% [v/v] OADC. CFUs were counted after 21 days of incubation and all timepoints were plotted in GraphPad Prism as CFU/mL over Time (days).

### BioB over-expressed mutant MIC assay

The potency of the bioB inhibitor acidomycin-amide was assessed by measuring MICs against Mtb H37Rv and a BioB over-expressing strain. Briefly, Wild-type (WT) H37Rv and BioB-OE were cultured in 10 mL of GAST medium + 2 µM biotin (with Zeocin at 25 µg/mL for the BioB-OE strain) in 25-cm^2^ tissue culture flasks with vented caps for approx. 7 d at 37 °C and 5% CO_2_ in a humidified incubator, growing to OD_580_ = ∼1.0. The cultures were then washed twice with 10 mL GAST without biotin and were diluted to OD_580_ = 0.005, +/- 2 µM biotin. Compounds were solubilized in DMSO and dispensed into black, clear-bottom 384-well tissue culture plates using an HP D300e Digital Dispenser as 13-point, logarithmically distributed titrations from 0.01-100 µM in triplicate. 50 μL of OD_580_ = 0.005 suspension was pipetted using an Integra 16 Channel VIAFLO Electronic Pipette into each well, and plates were incubated for 14 days at 37 °C in the same conditions as above in stacks of no more than six plates wrapped with aluminum foil. Final OD_580_ values were normalized from 0–100% to the averages of no-drug (100%) and no-growth (0%) control wells. MIC values were calculated by fitting the log(inhibitor) vs. response data in GraphPad Prism to a Gompertz model provided by GraphPad (https://www.graphpad.com/support/faqid/1365/), with bottom and span best-fit values constrained to 0 and 100, respectively. When the calculated MIC was higher than the highest dose tested, it was reported to be greater than that dose. All MIC measurements were made at least twice, in independent replicate experiments.

### Chequerboard synergy assay

Combinational activity of **C48** and acidomycin-amide was evaluated against Mtb H37Rv in 384-well plates using a chequerboard method, with the combination of **C48** plus bedaquiline also included, as an experimental control. Briefly, WT H37Rv was cultured in 10 mL of GAST medium supplemented with 2µM biotin in a 25-cm^2^ tissue culture flask with a vented cap for approx. 7 d at 37 °C and 5% CO2 in a humidified incubator, growing to OD_580_ = ∼1.0. After 7 d, the culture was washed twice with 10 mL GAST without biotin and subsequently diluted to OD_580_ = 0.005 in 10 mL GAST medium. Compounds were solubilized in DMSO and dispensed into black, clear-bottom 384-well tissue culture plates using the Synergy feature of an HP D300e Digital Dispenser as 14-point, 2-fold dilution series for each compound. 50 μL of the OD_580_ = 0.005 suspension was pipetted or dispensed using a Thermo Fisher Multidrop Combi Reagent Dispenser into each well, and plates were incubated for 14 d at 37 °C in the same conditions as above in stacks wrapped with aluminum foil. Final OD_580_ values were normalized from 0–100% to the averages of no-drug (100%) and no-growth (0%) control wells. The fractional inhibitory concentration index (FICI) was used to access the results. The FICI was calculated for each drug combination by using the following formula, FIC_A_ + FIC_B_ = FICI, in which FIC_A_ is the MIC of Compound A in combination with Compound B divided by the MIC of Compound A alone, and FIC_B_ is the MIC of Compound B in combination with Compound A divided by the MIC of Compound B alone. Synergy was defined as a FICI ≤ 0.5, while a FICI > 0.5, but ≤ 4.0, was defined as no interaction, and a FICI > 4.0 was defined as antagonism. The combination of **C48** and acidomycin-amide was determined to be synergistic, while the combination of **C48** and bedaquiline yielded no interaction.

### C48 and acidomycin-amide combination killing assay

To analyze the effect of combining **C48** with acidomycin-amide and determine the minimum bactericidal concentration (MBC), briefly, WT H37Rv was cultured in 10 mL of GAST medium supplemented with 2 µM biotin in a 25-cm^2^ tissue culture flask with a vented cap for approx. 7 d at 37 °C and 5% CO_2_ in a humidified incubator, growing to OD_580_ = ∼1.0. After 7 d, the culture was washed twice with 10 mL GAST without biotin and subsequently diluted to OD_580_ = 0.001 in 600 mL GAST medium. In triplicates, 25-cm^2^ tissue culture flasks with vented caps were filled with 6 mL of culture and a concentration of **C48**, acidomycin-amide, or both, ranging from 0.188-48x the liquid IC50s. Additionally, a set of triplicate flasks containing no drugs were filled as controls. DMSO was added, as needed, to normalize all flasks to the highest percentage of DMSO [v/v] required, 0.24%. Killing was monitored by plating serial dilutions on 7H10 agar plates supplemented with 0.5% [v/v] glycerol and 10% [v/v] OADC, on Day 0 and ∼Day 40. CFUs were counted after 21 d of incubation and all timepoints were plotted in GraphPad Prism as CFU/mL over Fold of IC_50_.

### MIC screening against *M*. *bovis*, *M. africanum*, and all drug sensitive and resistant Mtb

*M*. *bovis*, *M*. *africanum*, and all drug sensitive and resistant Mtb isolates were grown in Dubos-based medium (6.5 g/L Dubos broth base, 0.81 g/L NaCl, 7.5 g/L glucose, and 5 g/L BSA fraction V) to an OD_650_ = 0.2. Cells were diluted 1000-fold in this medium, and 50 μL per well was dispensed into round-bottom clear sterile polypropylene plates containing either **C48** or linezolid serially diluted from 50 to 0.049 μM or from 0.5 to 0.00049 μM in the same medium at 50 μL per well. Plates were incubated for up to 2 weeks at 37 °C, and MIC was scored under an inverted enlarging mirror. The ability of biotin to rescue growth was confirmed by performing the MIC under identical conditions with the addition of biotin (final concentration 0.5 mg/mL) to the media. MIC determinations were performed in duplicate for each concentration and repeated independently at least two times.

### MIC screening against Gram-negative pathogens

MIC measurements were based on a method previously described^15^. Overnight liquid cultures of *E. coli* BW, *E. coli* C0244, *A. baumannii* ATCC 17978, *K. pneumoniae* ATCC 43816, *P. aeruginosa* PA01 and *E. faecium* ATCC 19434 strains were inoculated with a single colony from a freshly streaked agar plate of the corresponding bacterial strain and cultured in M9 minimal medium supplemented with amino acids^15^. The bacterial culture was prepared by washing 1 mL of overnight culture in PBS pH 7.4 (x3), then diluted to OD_600_ = 0.5 and used to inoculate 1:1000 in M9 minimal medium with amino acids in 96-well plates. Compounds were added to 96-well plates in two-fold serial dilutions. The plates were incubated stationary at 37 ℃ for 18 h, then OD_600_ was measured. The MIC values were determined to be the minimal concentration that fully inhibits bacterial growth. The assay was performed in duplicate.

### Membrane fluidity measurements

Mtb cultures were grown with the indicated concentration of compound **C48** in GAST medium with or without addition of 2 μM biotin to OD_600_ = 0.6. Then, laurdan (D250, Thermo Fisher) dissolved in dimethylformamide (DMF) was added to a final laurdan concentration of 10 µM and a final DMF concentration of 1% (v/v). Laurdan cultures were incubated for 2 h at 37 °C with shaking and then collected by centrifugation at 3,200 × g for 10 min at 20 °C. Samples were washed 4 times in GAST medium supplemented with 1% (v/v) DMF, then resuspended in 1/50 initial culture volume of GAST medium + 1% (v/v) DMF. Samples were transferred to black 96-well plates (3915, Corning), and fluorescence was measured in a BioTek Synergy H1 plate reader (Agilent) first at 20 °C, then at 37 °C after rapidly increasing the internal temperature of the plate reader. Laurdan was excited at 350 nm, and emission was monitored over a range of 440–490 nm. Fluorescence intensity (I) measurements were converted into the laurdan GP metric: Laurdan GP=*I*440−*I*490 / *I*440+*I*490. n=3 biological replicates, where each biological replicate was measured in technical duplicate and the technical duplicates were averaged. Data are represented as the mean +/- standard deviation of the biological triplicate measurements.

### Lipid labeling, extraction, and analysis

^14^C-acetate labeled fatty acid methyl esters and mycolic acid methyl esters were prepared as described earlier^54^. Mtb H37Rv was grown in Dubos-based medium to an OD_650_ = 0.38 and split into four 25-mL cultures in 250 mL roller bottles. To two cultures, **C48** was added to 0.1 mM while an equivalent volume of DMSO was added to the other 2 cultures. Cells were incubated (37 °C, 100 rpm) for 24 or 72 hours, after which cells were harvested and each pellet resuspended in 10 mL fresh Dubos-based medium with 0.1 mM **C48** or DMSO, respectively and transferred to 30 mL inkwell bottles. To each culture, 10 mL 0.5 mCi/mL 1-^14^C-acetate (Moravek Biochemicals #MC125) was added and cultures incubated for 14 hours (37 °C, 100 rpm) after which cells were harvested and washed twice with Dubos-based medium. Cell pellets were resuspended in 0.25 mL water and 0.25 mL 40% tetrabutylammonium hydroxide (Sigma Aldrich) in glass tubes and boiled for 3 hours. The samples were cooled after which 1 mL CH_2_Cl_2_ and 0.1 mL CH_3_I was added. Samples were mixed for 30 minutes and centrifuges at 750×g for 10 minutes. The top aqueous phase was removed and the organic layer washed once with 0.1 M HCl. The organic layer containing the FAMEs and MAMEs was dried down under argon. Lipids were resuspended in 75 mL diethyl ether and an equivalent amounts of counts spotted to Silica TLC plates (Sigma Aldrich, 250 mm, 20 x 20cm) which were developed three times in petroleum ethers: diethyl ether (85:15). Dried TLC plates were exposed to phosphor imager plates and analyzed with an Typhoon imager and image analysis with ImageQuant software (GE Healthcare).

### Pharmacokinetic, dose escalation and lung distribution studies

CD-1 female mice (22-25 g) were used in pharmacokinetic studies. Compounds administered by intravenous injection (IV) were dosed at 5 mg/kg dose and formulated in 5% Dimethylacetamide (DMA) and 95% - 4% Cremophor EL. IV formulations were filtered to 0.22 µm prior to dosing. The filtrate was quantified by LC-MS to adjust for losses during filtration. Compounds dosed by oral gavage (PO) at 25mg/kg were formulated in 5% DMA, 60% Polyethylene Glycol 300, and 35% of 5% dextrose in water (D5W). Aliquots of 50 μL of blood were taken by puncture of the lateral tail vein from each mouse (n = 3 per route and dose) at 5 min, 15 min, 1, 3, 7, and 24 h post-dose for IV and 30 min, 1, 3, 5, 7, and 24 h post-dose for PO. Blood was captured in CB300 blood collection tubes containing K_2_EDTA and stored on ice. Plasma was recovered after centrifugation and stored at -80 °C until analyzed by high pressure liquid chromatography coupled to tandem mass spectrometry (LC-MS/MS). Dose escalation pharmacokinetic profiling was performed at doses of 50, 100, and 200 mg/kg after 4 days of QD PO dosing to assess exposure linearity and tolerability. Compounds were formulated using 0.5% Carboxymethyl Cellulose and 0.5% Tween 80 (CMC/Tween) in water and stirred overnight to micronize particles. A tissue distribution study using **C48** formulated as previously described was performed after a single QD dose of 200 mg/kg. Terminal blood and lung samples were obtained at 1, 3, and 7 h, using n=3 mice at each time point. Samples were stored at -80 °C until analysis by LC-MS/MS. Pharmacokinetic parameters were determined with the PK Solver Excel add-in using non-compartmental pharmacokinetic analysis.

### LC-MS/MS analytical methods

Neat 1 mg/mL DMSO stocks of compounds were serial diluted in 50/50 acetonitrile (ACN)/ milli-Q water to create standard curve solutions. Drug free CD-1 mouse lung tissues collected in house were weighed and homogenized in 4 volumes of phosphate-buffered saline (PBS) to a final 5x dilution factor for use in standard curves. Homogenization was achieved using a FastPrep-24 instrument (MP Biomedicals) and 1.4 mm zirconium oxide beads (Bertin Corp.). Standards were created by adding 10 µL of spiking solutions to 90µL of drug free plasma (CD-1 K_2_EDTA Mouse, Bioreclamation IVT) or 90 µL of drug free mouse lung homogenate. 10 µL of control, standard, or study sample was added to 100 µL of ACN protein precipitation solvent containing 10 ng/mL of the internal standards Verapamil (Sigma Aldrich). Extracts were vortexed for 5 min and centrifuged at 4000 rpm for 5 min. 75 µL of supernatant was transferred for LC-MS/MS analysis and diluted with 75 µL of Milli-Q deionized water. LC-MS/MS analysis was performed on a Sciex Applied Biosystems Qtrap 6500^+^ triple-quadrupole mass spectrometer coupled to a Shimadzu Nexera X2 UHPLC system to quantify each drug in plasma. Chromatography was performed on an Agilent SB-C8 (2.1x30 mm; particle size, 3.5µm) using a reverse phase gradient. Milli-Q deionized water with 0.1% formic acid was used for the aqueous mobile phase and 0.1% formic acid in ACN for the organic mobile phase. Multiple-reaction monitoring of parent/daughter transitions in electrospray positive-ionization mode was used to quantify all the analytes. The following MRM transitions were used for detection of the analytes **C21** (416.91/200.90), **C48** (418.00/201.00) and verapamil (455.2/165.2). Sample analysis was accepted if the concentrations of the quality control samples were within 20% of the nominal concentration. Data processing was performed using Analyst software (version 1.6.2; Applied Biosystems Sciex).

### Efficacy studies in the BALB/c mouse model of acute TB infection (biotin-rich chow)

BALB/c mice (9-week-old females; weight range, 18-20 g, Charles River) were maintained under specific pathogen-free conditions and fed water and chow *ad libitum* (PicoLab^®^ Rodent Diet 20). Mouse body weights and behavior were monitored daily for signs of distress. Rifampicin (Sigma) and **C48** were prepared for oral gavage (PO) in 0.5% Carboxy Methyl Cellulose & 0.5% Tween 80 in water (CMC/Tween) and kept at 4 ℃ up to 7 days. RIF was dosed at 10 mg/kg. **C48** was dosed at 100 mg/kg and 200 mg/kg. All drugs and vehicle were dosed at 8mL/kg. Mice were infected with an inoculum of Mtb H37Rv mixed with 5 mL of PBS (3×10^6^ CFU/mL) using a Glas-Col whole-body aerosol unit. This resulted in an average measured lung implantation of 2.29 log10 (195) CFU per mouse at 1 day post infection. Groups of 5 mice were euthanized at the start of treatment (2 weeks post-infection) and after receiving test compound by oral gavage (PO) daily (QD) at specific time points up to 28 days of dosing. Whole lungs were homogenized in 5 mL of PBS containing 0.05% Tween 80 and CFUs were quantified by plating serial dilutions of homogenates onto Middlebrook 7H11 agar with 10% OADC. Colonies were counted after at least 21 days of incubation at 37 °C. Log CFU/lung data plots and statistical analysis was performed using GraphPad Prism 9.5.1. Statistical significance was determined using an ordinary one-way analysis of variance (ANOVA) and a Dunnet’s *post hoc* test for multiple comparisons. Log-transformed CFU were used to calculate means and standard deviations.

### Biotin deficient mouse chow TB infection pilot study

BALB/c mice (9-week-old females; weight range, 18-20 g, Charles River) were maintained under specific pathogen-free conditions and fed water. Biotin deficient chow (Inotiv TD.81079) was given *ad libitum* to the first group to reduce serum biotin to humanized levels, while a matching chow supplemented with biotin (Inotiv TD.97126) was given to a second group. Mouse body weights and behavior were monitored daily for signs of distress. Both groups of mice were infected simultaneously in the same infection chamber with an inoculum of Mtb H37Rv mixed with 5 mL of phosphate-buffered saline (PBS) (1×10^7^ CFU/mL) using a Glas-Col whole-body aerosol unit. At day 1, this resulted in an average measured lung implantation of 2.52 log10 (334) CFU per mouse (n = 5) for the group fed biotin deficient chow and 2.53 log10 (340) CFU per mouse (n=5) for the group fed biotin supplemented chow. Timepoints continued in the same manner at 14-, 28-, and 42-days post infection. Lung processing and CFU counting were performed as previously described.

### C48 biotin deficient TB efficacy study

BALB/c mice (9-week-old females; weight range, 18-20 g, Charles River) were maintained under specific pathogen-free conditions and fed water. Biotin deficient chow (Inotiv TD.81079) was given *ad libitum*. Isoniazid (Sigma) and **C48** were prepared for oral gavage (PO) in CMC and kept at 4 ℃ up to 7 days. Isoniazid was dosed at 25 mg/kg. **C48** was dosed at 200 mg/kg All drugs and vehicle were dosed at 8 mL/kg. Mice were infected as previously described in the pilot study. At day 1, this resulted in an average measured lung implantation of 2.04 log10 (110) CFU per mouse (n = 5 to 7). Groups of 6 mice were sacrificed by cervical dislocation at the start of treatment (1 week post-infection) and after receiving test compound by oral gavage (PO) daily (QD) at specific time points up to 21 days of dosing. Lung processing, CFU counting, and statistical analysis were performed as previously described.

### Quantification of biotin in mouse plasma

For determination of Biotin in plasma, IDK Biotin ELISA Kit (Kit # KR 8141 Immundiagnostik AG, Germany) was used. Plasma samples were diluted 10X (1:9) with sample dilution buffer. 50 µl of standards, samples and controls were added into respective wells that are pre-coated with streptavidine, covered with foil and incubated for 30 min at room temperature. After incubation, the wells were washed 5 times with 250 µL of wash buffer and firmly tapped on absorbent paper on the final washing step. 50 µL of conjugate (enzyme-labelled biotin) was added to each well, which competed against the biotin in the samples, standards and controls for streptavidin on the microtiter plate. After adding the conjugate, the microtiter plate was covered with foil and incubated for 30 min at room temperature. After incubating with the conjugate, wells were washed 5 times with 250 µL of wash buffer, which washed away unbound enzyme-labeled biotin, and firmly tapped on absorbent paper after the final wash. 100 µL of substrate was added in each well, resulting in a color reaction. The plate was incubated for 10-15 min until good color differentiation was achieved, at which point 100 µL of ELISA stop solution was added to all wells. The microplate was read at 450nm against 620 nm as a reference. Results were generated using 4 parameter algorithms. Linear ordinate for the optical density and a logarithmic abscissa for the concentration was used to calculate the absolute biotin levels in plasma.

### Cell cytotoxicity-MTT assay

Cells (HepG2 and HT-29) were cultured in medium (Eagle’s Minimum Essential Medium for HepG2 and McCoy’s 5A Medium for HT-29) supplemented with 10% fetal bovine serum (FBS), 100 I.U./mL penicillin, and 100 μg/mL streptomycin. Cells were plated in 96-well plates at (2.5−5.0) × 10^4^ cells per well. After incubation at 37 °C for 24 h, cells were treated with **C48** at 2-fold serial diluted concentrations ranging from 100 μM to 3.125 μM, in addition to 1% DMSO as negative control and 1% Triton X-100 as positive control. Treated cells were incubated for 48 h at 37 °C in a 5% CO_2_/95% air humidified atmosphere. Measurement of cell viability was carried out using CyQUANT XTT cell viability assay kit (Invitrogen Cat. X12223). Briefly, 6:1 of XTT reagent and electron coupling reagent was combined as the working solution prior to testing. 70 μL of the working solution was added directly to 96-well plates. After incubation at 37 °C for 4 h, the plate was read on a M5e spectrophotometer (Molecular Devices) at 450 nm and 660 nm for background subtraction. Cell viability was estimated as the percentage absorbance of sample relative to the DMSO control. n = 3 biological replicates, where each biological replicates was measured in technical duplicate and the technical duplicates were averaged.

### RNA isolation and sequencing

(note: all experiments were done at 4 °C until samples were applied to the kit, after which extraction was done at room temperature) RNA was harvested from triplicate Mtb H37Rv cultures. Log-phase Mtb H37Rv grown in Sauton’s media was diluted to an OD_600_ = ∼0.1 in 250 mL inkwell bottles containing at least 30 mL of the same medium and subjected to 10 μM **C48**, or equal-volume DMSO, in triplicates. Cells were incubated with shaking at 37 °C for 64 h. Cells were centrifuged at 3,900 *x g* for 10 min at 4 ℃, and the resulting pellets were resuspended in 500 μL TRIzol reagent (Invitrogen) containing 1% polyacryl carrier (Molecular Research Center). Next, cells were lysed by two 1 min rounds of bead beating (Biospec) at maximum speed. Samples were centrifuged at 10,000 rpm for 30 s to remove beads. 50 μL of 1-bromo-3-chloropropane (Sigma-Aldrich) was added to the lysed samples, which were subsequently centrifuged at 10,000 rpm for 15 min to separate phases. An equal volume of ethanol (Fisher Bioreagent) was added to the aqueous phase. After that, total RNA was isolated, and DNA was removed using the Direct-zol RNA MiniPrep Plus Kit (Zymo Research) as outlined by the manufacturer. Around 1 μg of total bacterial RNA was depleted of rRNA, and cDNA libraries were generated using the Stranded Total RNA Prep with Ribo-Zero PlusMicrobiome kit (Illumina Inc) by SeqCenter. Final libraries were sequenced on an Illumina Novaseq platform with 150 bp paired-end reads, generating a total of 12 million reads.

### RNA-seq data analysis

Raw reads were pre-processed using an established pipeline (*MDHowe4/RNAseq-Pipeline: RNA-Sequencing Pipeline for Analysis of Prokaryotic Paired-End Read Expression Data Utilizing STAR*, n.d.). Briefly, FASTQ files were quality-checked using fastqc (v0.11.7), quality trimmed using Cutadapt (v2.4)^55^, and aligned to the Mtb H37Rv genome (NC_000962.3) using Spliced Transcripts Alignment to a Reference (STAR) (v2.7.1a)^56^. Total reads for each gene were computed using feature Counts^57^. Genes with less than 10 reads were omitted, and differential expression analysis was conducted on the resulting feature counts files using the R (v4.3.2) package DESeq2 (v1.34.0)^58^. Genes that met log_2_-fold change cut-off of > 1 or < -1 and adjusted *p*-value cut-off of < 0.05 were considered to be significantly differentially expressed. The outputs were visualized using R packages tidyverse, ggpubr, and Enhanced Volcano.

### Statistical analyses

Experiments were conducted on at least two independent occasions and the resulting data are presented as the arithmetic mean of these biological repeats unless stated otherwise. GraphPad Prism 9.5.1 was used for data analysis and figure generation. Data are shown as means ± s.e.m. *p* < 0.05 was considered statistically significant. In this study, no statistical methods were used to predetermine sample size. The investigators were not blinded to allocation during the experiments and outcome assessments.

### Reporting summary

Further information on research design is available in the Nature Portfolio Reporting Summary linked to this article.

### Materials availability

All reagents generated in this study are available upon request from the corresponding author.

### Data availability

All relevant data generated in this study are present within the manuscript and its Supplementary Information. The crystal structure of *Mt*BioA with ligand **C48** has been deposited to the Protein Data Bank with accession code 9D7M. The genome sequences of resistant mutants are available at BioProject database under accession code PRJNA1288510. The RNA-sequencing data have been deposited to the BioProject database under accession code PRJNA1291460. The source data has been provided for Fig. 1B, Fig. 2A-H, Fig. 3A-F, Fig. 4A-F, Fig. 5A-E, Fig. 6A-F, Supplemental Fig. S4A and S4C. The data supporting the findings of this study are available from the corresponding author on reasonable request.

## Acknowledgements

We thank Aldrich lab members Tsung-Yun Wong and Peng Ge for input on this project, Adhar Manna and Ambrose Cheung from Geisel School of Medicine at Dartmouth for screening against S. aureus, Rodion Gordzevich, Megan M. Tu and Eric D. Brown from University of McMaster for screening against gram-negative pathogens, Camilla Folvar, Betelhem Tatek and Firat Kaya from Center for Discovery and Innovation, Hackensack Meridian Health for assisting in vivo studies. We also thank Curtis A. Engelhart from Weill Cornell Medical College for performing MIC study against WT Mtb. This work was supported by a grant (AI143784 to C.C.A., D.S., and M.D.Z.) from the National Institutes of Health. Mass spectrometry was carried out in the Analytical Biochemistry Shared Resource of the Masonic Cancer Center, University of Minnesota, funded in part by Cancer Center Support Grant CA-77598. This research was supported in part by the Intramural Research Program of the National Institutes of Health (NIH). The contributions of the NIH author(s) were made as part of their official duties as NIH federal employees, are in compliance with agency policy requirements, and are considered Works of the United States Government. However, the findings and conclusions presented in this paper are those of the author(s) and do not necessarily reflect the views of the NIH or the U.S. Department of Health and Human Services. The X-ray diffraction experiment was supported by an agreement between the Advanced Photon Source, a U.S. Department of Energy (DOE) Office of Science user facility operated for the DOE Office of Science by Argonne National Laboratory under Contract No. DE-AC02-06CH11357, and the Diamond Light Source, the U.K.’s national synchrotron science facility, located at the Harwell Science and Innovation Campus in Oxfordshire, where the work was performed under proposal AU38824.

## Author contributions

Q.L., D.S. and C.C.A. conceptualized the study, Q.L. designed and carried out experiments and data analysis. J.B.W. performed the microbiology assays. Y.P.J. and D.R.R. obtained and solved the X-ray crystal structure. M.R.S., K.M. performed and E.J.R. supervised the membrane fluidity assay and NTM screening. J.P., C.F. and B.T. performed the in vivo studies. S.R. performed LC-MASS analysis. S.V., Z.J., L.O. and A.D.B. performed microbiology and RNA-sequencing. H.I.M.B. performed drug-resistant potency screening. V.D. and M.D.Z. supervised the PK and animal study. Q.L. and C.C.A. wrote the initial manuscript. All authors have given approval to the final manuscript.

