## Supplemental Information for "Validating Conditionally Essential Targets: Discovery of the First Orally Effective Biotin Inhibitor against *Mycobacterium Tuberculosis*"

^†^contributed equally

**General materials and methods.**

Chemicals and solvents were purchased from Alfa Aesar, Oakwood chemicals, Chem-Impex International, Acros Organics, TCI America and Sigma-Aldrich, and were directly used as received. An anhydrous solvent dispensing system using two packed columns of molecular sieves were used for drying DMF, while two packed columns of neutral alumina were applied to dry CH_2_Cl_2_, and the solvents were dispensed under nitrogen gas (N_2_). Anhydrous grade dioxane was purchased from Sigma-Aldrich. EtOAc and hexanes were purchased from Fisher Scientific. All reactions were performed in oven-dried (150 °C) glassware under an inert atmosphere of dry nitrogen gas (N_2_). TLC analyses were carried out on TLC silica gel plates 60F_254_ purchased from Sigma-Aldrich and were visualized by UV light lamp. Purification by flash chromatography was performed using a medium-pressure flash chromatography system (Buchi) equipped with flash column silica cartridges with the indicated solvent system. Analytical reversed-phase HPLC purity was performed on a Waters XSelect 5 μm C18 150 × 4.6 mm column operating at 1 mL/min with detection at 254 nm employing a linear gradient from 5% to 95% MeCN (0.1% FA) in H_2_O (0.1% FA) for 10 min. ^1^H, ^13^C and ^19^F spectrums were acquired on 400 or 600 MHz NMR spectrometers (Agilent Scientific Instruments). Proton chemical shifts are recorded in ppm by an internal standard of residual dimethyl sulfoxide (2.50), methanol (3.31) and chloroform (7.26); carbon chemical shifts are recorded in ppm from an internal standard of residual dimethyl sulfoxide (39.5), methanol (49.1), or chloroform (77.2). Proton chemical data are reported as follows: chemical shift, multiplicity (s = singlet, d = doublet, dt = doublet of triplets, t = triplet, q = quartet, m = multiplet, ap = apparent, br = broad, ovlp = overlapping), coupling constant (s), integration. Melting points of final compounds were determined by Thomas Hoover capillary melting point apparatus. High-resolution mass spectra were obtained on an LTQ Orbitrap Velos (Thermo Scientific, Waltham, MA). All final compounds were determined to be > 95% purity by analytical reverse-phase HPLC. All animal studies were ethically reviewed and carried out in compliance with the Institutional Animal Care and Use Committee of Hackensack Meridian Health.

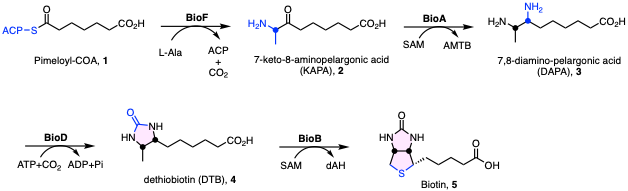

### **Supplementary Figure S1 Biotin biosynthesis in Mtb**

(**A**) Biotin biosynthesis in *Mtb*. Pimeloyl-ACP (**1**) is converted to 7- keto-8-aminopelargonic acid (KAPA, **2**) by BioF (KAPA synthetase). Transamination by BioA (DAPA synthetase) converts 2 to 7,8-diaminopelargonic acid (DAPA, **3**), followed by insertion of a carbonyl by BioD (dethiobiotin synthetase) gives rise to dethiobiotin (DTB, **4**). Finally, BioB (biotin synthase) is responsible for the conversion of 4 to biotin (5).

### **Supplemental Table S1. Pharmacokinetics of 6, C21 and C48 in CD-1 mice**

| Compd | i.v. PK parameters^a^ | | | | p.o. PK parameters^b^ | | |
| --- | --- | --- | --- | --- | --- | --- | --- |
|  | AUC_0–∞_ (*μ*g·hr/mL) | *V*d (L/kg) | CL (mL/kg·min) | *t*_1/2_ (h) | AUC_0–∞_ (*μ*g·hr/mL) | *F* (%) | AUC (p.o.)/MIC |
| **6** | 0.3 | 0.54 | 30.6 | 0.2 | 0.46 | 3.4 | 0.04 |
| **C21** | 4.0 | 0.37 | 5.3 | 0.82 | 108 | 100 | 57 |
| **C48^c^** | 3.3 | 0.26 | 3.2 | 0.91 | 146 | 100 | 1570 |

^a^i.v. dose (D_iv_) = 5 mg/kg, ^b^p.o. dose (D_po_) = 25 mg/kg. AUC_0–∞_, area under the plasma concentration−time curve from time 0 to infinity; *V*d, volume of distribution; CL, clearance; *t*_1/2_, terminal elimination half-life; *F*, relative oral bioavailability calculated as follows: *F* = 100 × [(AUC_po_ × D_iv_)/(AUC_iv_ × D_po_); AUC (p.o.)/MIC = AUC_0–∞_(p.o.)/MIC_50_]. ^c^The i.v. dose of **C48** was reduced to 0.648 mg/kg due to solubility issue. Each group contains 3 mice.

**Supplemental Scheme S1.** Synthesis of **C21**, **C48** and acidomycin-amide

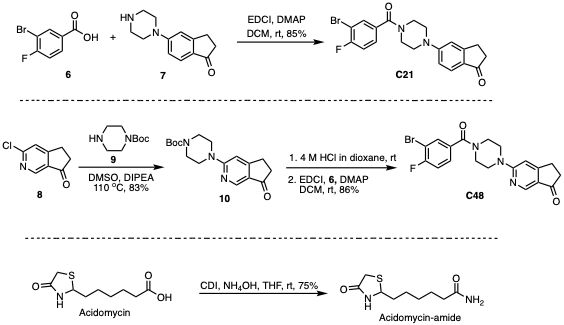

### *Synthesis of* ***C21***.

To a solution of carboxylic acid **6** (114 mg, 0.50 mmol, 1.0 equiv) and known amine **7**^1^ (130 mg, 0.60 mmol, 1.2 equiv) in CH_2_Cl_2_ (10 mL) EDCI (116 mg, 0.75 mmol, 1.5 equiv) and DMAP (6 mg, 0.050 mmol, 0.10 equiv) were added and the reaction was stirred for 16 h at 23 °C, quenched with addition of water (25 mL) and extracted by CH_2_Cl_2_ (3 × 20 mL). The combined organic layers were washed with brine, dried over Na_2_SO_4_ and concentrated under reduced pressure. Purification by flash chromatography on silica gel (1:3 hexanes/EtOAc) afforded the product **C21** (155 mg, 85%) as a white solid; *R*_f_ = 0.4 (1:3 hexanes/EtOAc); ^1^H NMR (400 MHz, CDCl_3_) δ 7.68 (dd, *J* = 6.5, 2.1 Hz, 1H), 7.65 (d, *J* = 8.6 Hz, 1H), 7.39 (ddd, *J* = 8.4, 4.6, 2.1 Hz, 1H), 7.19 (t, *J* = 8.3 Hz, 1H), 6.88 (dd, *J* = 8.7, 2.2 Hz, 1H), 6.83 (s, 1H), 4.12–3.51 (br, 4H), 3.42 (s, 4H), 3.11–2.98 (m, 2H), 2.72–2.60 (m, 2H); ^13^C NMR (101 MHz, CDCl_3_) δ 205.2, 168.2, 160.2 (d, *J*_C-F_ = 253.0 Hz), 158.0, 155.4, 133.0, 132.8 (d, *J*_C-F_ = 4.2 Hz), 129.1, 128.4 (d, *J*_C-F_ = 7.8 Hz), 125.4, 116.9 (d, *J*_C-F_ = 22.9 Hz), 115.1, 110.9, 109.8 (d, *J*_C-F_ = 21.8 Hz), 48.1, 36.5, 26.1; HRMS (ESI+) *m/z* calcd for C_20_H_19_BrFN_2_O_2_ [M+H]^+^ 417.0608, found 417.0604 (error 1 ppm). Melting point: 168 ℃.

### *Synthesis of Piperazine* ***10***.

To a 100 mL flask charged with N_2_, commercially available **8** (500 mg, 3.0 mmol, 1.0 equiv), amine **9** (670 mg, 3.6 mmol, 1.2 equiv) and DMSO (15 mL) were added, followed by the addition of DIPEA (780 mg, 6 mmol, 2.0 equiv). The reaction was stirred at 110 °C for 3 h, quenched with water (20 mL) and extracted by EtOAc (3 × 30 mL). The combined organic layers were washed with brine, dried with Na_2_SO_4_ and concentrated under reduced pressure. Purification by flash chromatography on silica gel (1:1 hexanes/EtOAc) afforded the Buchwald coupling product **10** (789 mg, 83%) as a grey solid. *R*_f_ = 0.3 (1:1 hexanes/EtOAc); ^1^H NMR (400 MHz, CDCl_3_) δ 8.61 (s, 1H), 6.53 (s, 1H), 3.72 (t, *J* = 5.2 Hz, 4H), 3.55 (t, *J* = 5.4 Hz, 4H), 3.16–2.91 (m, 2H), 2.78–2.49 (m, 2H), 1.49 (s, 9H); ^13^C NMR (101 MHz, CDCl_3_) δ 203.64, 164.71, 161.63, 154.93, 146.66, 123.99, 101.63, 81.49, 44.87, 36.42, 28.60, 25.80; HRMS (ESI+) *m/z* calcd for C_17_H_24_N_3_O_3_ [M+H]^+^ 318.1818, found 318.1820 (error 0.6 ppm).

### *Synthesis of* ***C48***.

To a solution of compound **10** (500 mg, 1.5 mmol, 1.0 equiv) in CH_2_Cl_2_ (10 mL) was added 4 M HCl in dioxane (5 mL) at 0 °C and the solution was stirred at 23 °C for 4 h. The solvent was removed and the crude product amine was dried under vacuum and used for the next step without further purification.

The above deprotection product was dissolved in CH_2_Cl_2_ (15 mL) and subsequently acid **6** (501 mg, 2.3 mmol, 1.5 equiv), EDCI (100 mg, 2.3 mmol, 1.5 equiv) and DMAP (18 mg, 0.15 mmol, 0.10 equiv) were added and the mixture was stirred at 23 °C for 8 h. The reaction was quenched with water (25 mL) and extracted by CH_2_Cl_2_ (3 × 20 mL). The combined organic layers were washed with brine, dried with Na_2_SO_4_ and concentrated under reduced pressure. Purification by flash chromatography on silica gel (EtOAc) afforded the product **C48** (538 mg, 86%) as a white solid, *R*_f_ = 0.3 (1:2 hexanes/EtOAc); ^1^H NMR (400 MHz, CDCl_3_) δ 8.54 (s, 1H), 7.90–7.57 (m, 1H), 7.41–7.31 (m, 1H), 7.15 (t, *J* = 8.3 Hz, 1H), 6.55 (s, 1H), 3.99–3.44 (m, 8H), 2.99 (t, *J* = 6.2 Hz, 2H), 2.69–2.30 (m, 2H); ^13^C NMR (101 MHz, CDCl_3_) δ 203.4, 168.2, 164.7, 161.2, 160.0 (d, *J*_C-F_ = 252.9 Hz), 146.23, 132.97, 132.76 (d, *J* = 4.1 Hz), 128.29 (d, *J* = 7.7 Hz), 124.29, 116.81 (d, *J* = 22.9 Hz), 109.65 (d, *J* = 21.4 Hz), 101.92, 44.9, 36.2, 25.7; ^19^F NMR (376 MHz, CDCl_3_) *δ* –103.7 (q, *J* = 6.2 Hz); HRMS (ESI+) *m/z* calcd for C_19_H_18_BrFN_3_O_2_ [M+H]^+^ 418.0566, found 418.0553 (error 0.7 ppm). Melting point: 182 ℃.

### *Synthesis of acidomycine-amide*.

The synthesis of acidomycin-amide was following reported procedure^2^. Briefly, to a solution of acidomycin (500 mg, 2.3 mmol, 1.0 equiv) in THF (20 mL) was added 1,1'-Carbonyldiimidazole (750 mg, 4.6 mmol, 2.0 equiv) and stirred at 23 °C. After 2 h, saturated aqueous ammonium hydroxide (0.7 mL, 16 mmol, 7.0 equiv) was added in one portion. After 2 h, the reaction was concentrated in vacuo and re-suspended in H2O (20 mL). The mixture was acidified to pH ~2 with 2 N HCl, and organic products were extracted with EtOAc (3 × 25 mL). Combined organic extracts were washed with saturated aqueous NaCl (50 mL), dried (MgSO4) concentrated in vacuo, and triturated with cold H_2_O (15 mL) to afford the title compound (375 mg, 76 %) as a white solid. *R*_f_ = 0.3 (EtOAc); ^1^H NMR (400 MHz, CD_3_OD) δ 4.76 (t, *J* = 6.4 Hz, 1H), 3.58–3.41 (m, 2H), 2.21 (td, *J* = 7.5, 1.7 Hz, 2H), 1.92–1.78 (m, 1H), 1.76–1.55 (m, 3H), 1.52–1.30 (m, 4H); ^13^C NMR (101 MHz, CD_3_OD) δ 179.11, 176.92, 59.06, 39.55, 36.29, 32.71, 29.82, 26.65, 25.88; HRMS (ESI+) *m/z* calcd for C_9_H_17_N_2_O_2_ [M+H]^+^ 217.1011, found 217.1013 (error 0.9 ppm).

### **Supplemental Table S2.** Reported biotin biosynthesis inhibitors in literature^3^

| Structure | Target Enzyme | Activity |
| --- | --- | --- |
| 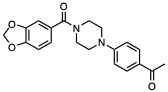 | BioA | IC_50_ of 155 nM against *Mt*BioA and MIC of 26 µM againt *Mtb*^1^ |
| 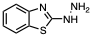 | BioA | Ki: 10.4 µM against *Mt*BioA^4^ |
| 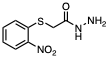 | BioA | IC_50_: 250 nM against *E. coli* BioA and MIC: 8 µg/mL against *E. coli*^5^ |
| 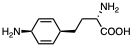 | BioA | Ki: 12 µM against *Mt*BioA^6^ |
| 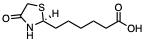 | BioB | Ki: 1 µM against *Mt*BioB and MIC: 0.6 µg/mL against *Mtb*^7^ |
| 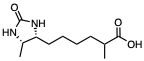 | BioB | MIC: 8 µg/mL against *E. coli*  0.5 µg/mL against *B. subtilis*  0.2 µg/mL against *M. avium*^8^ |
| 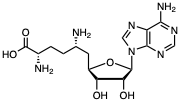 | BioC | 0.1 µM reduced 60% *B. cereus*  BioC activity, complete inhibition of BioC at 10 µM^9^ |
| 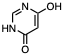 | BioD | Ki: 11 mM against *E. coli* BioD^10^ |
| 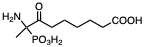 | BioF | *K*i = 7 µM against *Bacillus sphaericus* BioF^11^ |

### **Supplemental Figure S2.** Cocrystal of **C48/**BioA

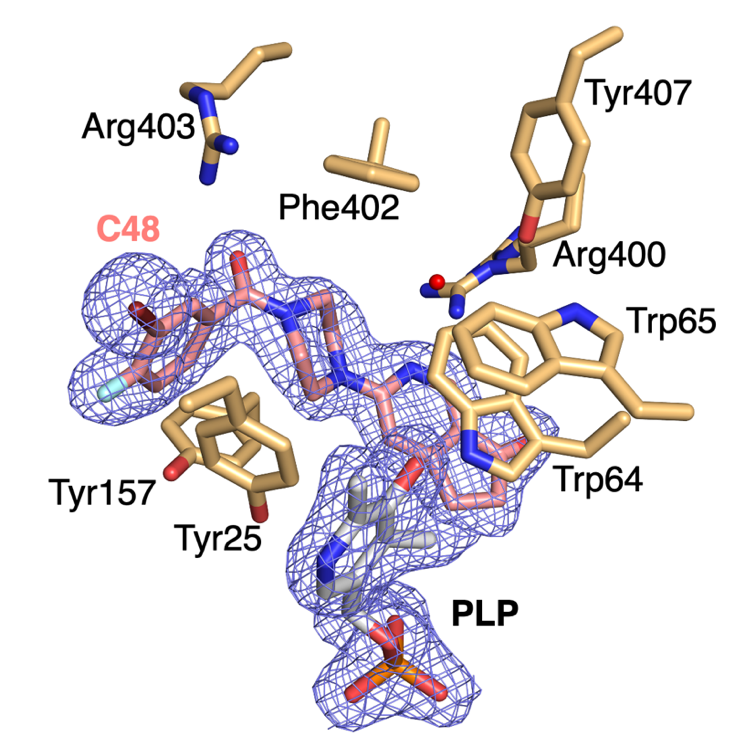

**Supplemental Figure S2.** The calculated 2Fo-Fc omit map highlighting **C48** and PLP in the BioA active site. The map is contoured at 1σ in purple. The carbon atoms of **C48** are pink, the carbon atoms of the BioA residues are tan, and the carbon atoms of PLP are gray. The nitrogen, oxygen, phosphorous, fluorine, and bromine atoms are blue, red, orange, cyan, and maroon, respectively.

### **Supplemental Table S3.** Crystallographic Data of C**48**

|  | **BioA_C48 (PDB: 9D7M)** |
| --- | --- |
| Wavelength | 0.97 |
| Resolution range | 62.83 - 1.97 (2.04 - 1.97) |
| Space group | P 21 21 21 |
| Unit cell |  |
| a, b, c (Å) | 63.1373 66.2907 204.841 |
| α, β, γ (°) | 90 90 90 |
| Total reflections | 614033 (16106) |
| Unique reflections | 61199 (5713) |
| Multiplicity | 11.3 (5.5) |
| Completeness (%) | 96.22 (80.60) |
| Mean I/sigma(I) | 37.85 (8.29) |
| Wilson B-factor | 20.37 |
| R-merge | 0.053 (0.16) |
| R-meas | 0.055 (0.17) |
| R-pim | 0.016 (0.07) |
| CC1/2 | 0.999 (0.97) |
| CC* | 1 (0.99) |
| Reflections used in refinement | 59446 (4904) |
| Reflections used for R-free | 1846 (112) |
| R-work | 0.1536 (0.19) |
| R-free | 0.1913 (0.24) |
| CC(work) | 0.962 (0.909) |
| CC(free) | 0.947 (0.869) |
| Number of non-hydrogen atoms | 7496 |
| macromolecules | 6488 |
| ligands | 82 |
| solvent | 926 |
| Protein residues | 860 |
| RMS(bonds) | 0.004 |
| RMS(angles) | 0.84 |
| Ramachandran favored (%) | 97.2 |
| Ramachandran allowed (%) | 2.1 |
| Ramachandran outliers (%) | 0.7 |
| Rotamer outliers (%) | 0 |
| Clashscore | 3.45 |
| Average B-factor | 21.68 |
| macromolecules | 20.37 |
| ligands | 16.95 |
| solvent | 31.31 |
| Statistics for the highest-resolution shell are shown in parentheses. | |

### **Supplemental Table S4.** Bacterial strains and cell lines used in this study

| Category | Genotype | Reference/Source |
| --- | --- | --- |
| *M.* tuberculosis (M. *tb*) | Wild-type (H37Rv) | ^12^ |
| *H37Rv ΔbioA TetON-1 (BioA-OE)* | BioA-Overexpress | ^13^ |
| *H37Rv ΔbioA TetON-5 (BioA-UE)* | BioA-Underexpress | ^13^ |
| BioB OE | BioB-Overexpress | ^7^ |
| *M. tb* HN878 | DS | ^14-16^ |
| *M. tb* Erdman | DS | ^14-16^ |
| *M. tb* *CDC1551* | DS | ^14-16^ |
| *M. tb* K03b00DS | DS | ^14-16^ |
| *M. tb* K04b00DS | DS | ^14-16^ |
| *M. tb* K05b00DS | DS | ^14-16^ |
| *M. tb* K07b00DS | DS | ^14-16^ |
| *M. tb* K08b00DS | DS | ^14-16^ |
| *M. tb* K09b00DS | DS | ^14-16^ |
| *M. tb* K10b00DS | DS | ^14-16^ |
| *M. tb* K11b00DS | DS | ^14-16^ |
| *M. tb* K12b00DS | DS | ^14-16^ |
| *M. tb* K13b00DS | DS | ^14-16^ |
| *M. tb* K14b00DS | DS | ^14-16^ |
| *M. tb* K15b00DS | DS | ^14-16^ |
| *M. tb* K16b00DS | DS | ^14-16^ |
| *M. tb* K17b00DS | DS | ^14-16^ |
| *M. tb* K35b00DS | DS | ^14-16^ |
| *M. tb* NIH_G13 | DS | ^14-16^ |
| *M. tb* NIH_G1DS | DS | ^14-16^ |
| *M. tb* NIH_G36 | DS | ^14-16^ |
| *M. tb* NIH_GA36 | DS | ^14-16^ |
| *M. tb* NIH_SA1 | DS | ^14-16^ |
| *M. tb* NIH_SA3 | DS | ^14-16^ |
| *M. tb* NIH_SA39 | DS | ^14-16^ |
| *M. tb* NIH_SA23 | DS | ^14-16^ |
| *M. tb* NIH_SA27 | DS | ^14-16^ |
| *M. tb* NIH_SA43 | DS | ^14-16^ |
| *M. tb* NIH_SA125 | DS | ^14-16^ |
| *M. tb* NIH_G12 | DS | ^14-16^ |
| CRC clinical strain K5072429 | DS | ^14-16^ |
| *M. tb* NIH_G9R | MCs | ^14-16^ |
| *M. tb* NIH_G10R | HRESRb | ^14-16^ |
| *M. tb* NIH_G11R | ECs | ^14-16^ |
| *M. tb* NIH_G16R | SCs | ^14-16^ |
| *M. tb* NIH_G17R | HEtCs | ^14-16^ |
| *M. tb* NIH_G21R | HRECsRb | ^14-16^ |
| *M. tb* NIH_G22R | SCs | ^14-16^ |
| *M. tb* NIH_G76MR | HROML | ^14-16^ |
| *M. tb* NIH1B314 | HEtCsOThLZ | ^14-16^ |
| *M. tb* 001K113 | HRSKPCpAkPtOMLRbZ | ^14-16^ |
| *M. tb* 28K111 | HRESKPCpAkPtCsOMLRbZLz | ^14-16^ |
| *M. tb* OK116 | HRESOMLPtCsRb | ^14-16^ |
| *M. tb* K37b00XR | HEKOPRM | ^14-16^ |
| *M. tb* CRC clinical strain 00202293 | HRESKPEtML | ^14-16^ |
| *M. tb* CRC clinical strain 01291696 | HRS | ^14-16^ |
| *M. tb* 026K111 | HRESKPCpAkPtCsOMLRbZLz | ^14-16^ |
| *M. tb* 053K113 | HRSKPCpAkPtOMLRbZLz | ^14-16^ |
| *M. tb* K21b00MR | HRES | ^14-16^ |
| *M. tb* K22b00MR | HRERb | ^14-16^ |
| *M. tb* K25b00MR | HREZRbTh | ^14-16^ |
| *M. tb* K26b00MR | HREZRb | ^14-16^ |
| *M. tb* CRC clinical MDR strain 01291696 | HRS | ^14-16^ |
| *M. tb* K18b01MR | HRERb | ^14-16^ |
| *M. tb* K29b00MR | HRSPO | ^14-16^ |
| *M. tb* NIH_G269DR | HRERb | ^14-16^ |
| *M. tb* NIH_G367DR | HROM | ^14-16^ |
| *M. tb* K20b00MR | HREZSKP | ^14-16^ |
| *M. tb* K32b00MR | HREKP | ^14-16^ |
| *M. tb* K33b00MR | HREZSKPTh | ^14-16^ |
| *M. tb* NIH_G5MR | HREKO | ^14-16^ |
| *M. tb* NIHB188 | HRZThRbLOM | ^14-16^ |
| *M. tb* Kb019 | HREPKOTh | ^14-16^ |
| *M*. bovis 0AF2122 | DS | ^14-16^ |
| *M. africanum* | DS | ^14-16^ |
| *M. abscessus* (ATCC19977) | Reference strain | ^17^ |
| *M. abscessus* T35 | Clinical strain | ^17^ |
| *M. abscessus* T37 | Clinical strain | ^17^ |
| *M. abscessus* T38 | Clinical strain | ^17^ |
| *M. abscessus* T49 | Clinical strain | ^17^ |
| *M. abscessus* BWH-B | Clinical strain | ^17^ |
| *M. abscessus* BWH-D | Clinical strain | ^17^ |
| *E. coli* C0244 | Gram-negative | IIDR clinical isolate collection |
| *E. coli* BW | Gram-negative | ^3^ |
| *A. baumannii* ATCC 17978 | Gram-negative | https://www.atcc.org/products/17978 |
| *K. pneumoniae* ATCC 43816 | Gram-negative | ^3^ |
| *P. aeruginosa* PA01 | Gram-negative | ^3^ |
| *E. faecium* ATCC 19434 | Gram-negative | https://genomes.atcc.org/genomes/b6  890c9e9268452d |
| *S. aureus* | Gram-positive | ^18^ |
| HepG2 | Mammalian | from ATCC |
| HT-29 | Mammalian | from ATCC |
| DS = Drug Susceptible Strains. Letter codes represent drug(s) to which that strain is resistant. Ak = Amikacin, Cp = Capreomycin, Cs = D-Cycloserine, E = Ethambutol, Et = Ethionamide, H = Isoniazid, K = Kanamycin, L = Levofloxacin, Lz = Linezolid, M = Moxifloxacin, O = Ofloxacin, P = *p*-Aminosalicylic acid, R = Rifampicin, Rb = Rifabutin, S = Streptomycin, Pt = Prothionamide, Z = Pyrazinamide, Th = Thiacetazone. | | |

### **Supplemental Table S5.** MIC screening of **C48** against a panel of pathogens

| Strains | Type | **C48** MIC (µM)^b^ | Linezolid MIC (µM)^b^ |
| --- | --- | --- | --- |
| *M. tb* Erdman | DS^a^ | 0.024 | 1.6 |
| *M. tb* K08b00DS | DS^a^ | < 0.024 | 1.56 |
| *M. tb* K15b00DS | DS^a^ | < 0.024 | 1.56 |
| *M. tb* K16b00DS | DS^a^ | < 0.024 | 1.56 |
| *M. tb* K17b00DS | DS^a^ | < 0.024 | 1.56 |
| *M. tb* K35b00DS | DS^a^ | < 0.024 | 1.2 |
| *M. tb* CRC clinical strain K5072429 | DS^a^ | < 0.024 | 1.2 |
| *M. tb* NIH_G12 | DS^a^ | < 0.024 | 2.3 |
| *M. tb* NIH_G1DS | DS^a^ | < 0.024 | 2.3 |
| *M. tb* NIH_GA36 | DS^a^ | < 0.024 | 1.6 |
| *M. tb* NIH_SA3 | DS^a^ | < 0.024 | ND^c^ |
| *M. tb* NIH_SA23 | DS^a^ | < 0.024 | ND^c^ |
| *M. tb* NIH_SA27 | DS^a^ | < 0.024 | ND^c^ |
| *M. tb* NIH_SA39 | DS^a^ | < 0.024 | ND^c^ |
| *M. tb* NIH_SA43 | DS^a^ | < 0.024 | ND^c^ |
| *M. tb* NIH_SA125 | DS^a^ | < 0.024 | ND^c^ |
| *M. abscessus* (ATCC19977) | Reference strain | 0.5^d^ | ND^c^ |
| *M. abscessus* T35 | Clinical strain | 0.5^d^ | ND^c^ |
| *M. abscessus* T37 | Clinical strain | 1^d^ | ND^c^ |
| *M. abscessus* T38 | Clinical strain | 1^d^ | ND^c^ |
| *M. abscessus* T49 | Clinical strain | 0.5^d^ | ND^c^ |
| *M. abscessus* BWH-B | Clinical strain | 0.5^d^ | ND^c^ |
| *M. abscessus* BWH-D | Clinical strain | 0.5^d^ | ND^c^ |
| *E. coli* C0244 | Gram-negative | inactive | ND^c^ |
| *E. coli* BW | Gram-negative | inactive | ND^c^ |
| *A. baumannii* ATCC 17978 | Gram-negative | inactive | ND^c^ |
| *K. pneumoniae* ATCC 43816 | Gram-negative | inactive | ND^c^ |
| *P. aeruginosa* PA01 | Gram-negative | inactive | ND^c^ |
| *E. faecium* ATCC 19434 | Gram-negative | inactive | ND^c^ |
| *S. aureus* | Gram-positive | inactive | ND^c^ |
| Type: DS^a^ = drug-sensitive Mtb clinical isolates. MIC^b^ = minimum inhibitory concentrations that resulted in complete growth inhibition. ND^c^ = Not determined. MIC^d^ = minimum inhibitory concentrations that resulted in 50% growth inhibition. | | | |

### **Structure Alignment of *Mt*BioA and ESKAPE BioA (Figure S3, Table S6 and S7)**

**>NP_216084.1 *Mycobacterium tuberculosis***

MAAATGGLTPEQIIAVDGAHLWHPYSSIGREAVSPVVAVAAHGAWLTLIRDGQPIEVLDAMSSWWTAIHGHGHPALDQALTTQLRVMNHVMFGGLTHEPAARLAKLLVDITPAGLDTVFFSDSGSVSVEVAAKMALQYWRGRGLPGKRRLMTWRGGYHGDTFLAMSICDPHGGMHSLWTDVLAAQVFAPQVPRDYDPAYSAAFEAQLAQHAGELAAVVVEPVVQGAGGMRFHDPRYLHDLRDICRRYEVLLIFDEIATGFGRTGALFAADHAGVSPDIMCVGKALTGGYLSLAATLCTADVAHTISAGAAGALMHGPTFMANPLACAVSVASVELLLGQDWRTRITELAAGLTAGLDTARALPAVTDVRVCGAIGVIECDRPVDLAVATPAALDRGVWLRPFRNLVYAMPPYICTPAEITQITSAMVEVARLVGSLP

**>WP_000131427.1 *Acinetobacter baumannii***

MTDNFDLEHIWHPYTSMTQPLPTFKVKRAYGATIELDDGRTLIDGMSSWWCAIHGYNHPELNQAVTDQLQNMSHIMFGGLTHDPAIELGKILLKITPPSLDKIFYADSGSVAVEVALKMAVQFWTAQGQPQKTNFITTRSGYHGDTWNAMSVCDPVTGMHQIFGTSLPNRLFVAAPQTKFHEEWNQEDIAELEQAIQQNHENLAALIIEPIVQGAGGMRFYHPEYLRQAKALCEKYHLLLIFDEIATGFGRTGKLFAWEHAQVEPDIMCLGKGLTGGYMTLSATLTTKHVAETISRGEAGVFMHGPTFMANPLACAVALKSTQLLIEQDWQANIKRIEQQLSQYLMPLNQLDYVADVRVLGAIGVVELTFNVDMKTLQQQFVERGIWIRPFGKLVYVMPPYVITQQELSDLLEHLVEVVKTMQGAH

**>WP_003687694.1 *Neisseria gonorrhoeae***

MPSEHQHTSSLLNFDRTHLLHPYTSMTDPLPVYPVKRAEGVFIELADGTRLIDGMSSWWCAIHGYNHPVLNQAVENQMKQMAHVMFGGLTHEPAVELGKLLVGILPQGLDRIFYADSGSVSVEVALKMAVQYQQARGLTAKQNIATVRRGYHGDTWNAMSVCDPETGMHHIFGSALPQRYFVDNPKNRFDDEWDGADLQPVRALFEAHHVDIAAFILEPVVQGAGGMYFYHPQYLRGLRDLCDEFDIVLIFDEIATGFGRTGKMFACEHAEVVPDIMCIGKGLSGGYMTLAAAITSQKVTETISRGEAGVFMHGPTFMANPLACAVACASVKLLLSQDWQANIRRIESILKGRLKAAWDIRGVKDVRVLGAIGVIELEKGVDMARFQADCVAQGIWVRPFGRLVYLMPPYIISDGILTKLADKTVQILKEHSK

**>WP_015367618.1 *Enterobacter aerogenes***

MTLDDLAFDRRHIWHPYTSMTSPLPVYPVVSAHGCELSLAGGEQLIDGMSSWWAAIHGYNHPRLNAAMKAQIDQMSHVMFGGITHPSAVALCRQLVAMTPESLECVFLADSGSVAVEVAMKMALQYWQAKGQPRRRFLTFRNGYHGDTFGAMSVCDPQNSMHSLWQGYLPDNLFAPAPQSRFDGEWDEMDMVPFARLMAAHRHEIAAVILEPVVQGAGGMRMYHPEWLKRVRKMCDREGILLIADEIATGFGRTGKLFACEHAGISADILCLGKALTGGTMTLSAAITTRTVAETISNGEAGCFMHGPTFMGNPLACAVASESLRLLESGEWQQQVAAIEAQLKAELAPARESEWVADVRVLGAIGVVETRQPVNMAALQRFFVEQGVWIRPFGRLIYLMPPYIISPQQLTRLTRAVNMAVQEETFFSE

**>**[**WP_002895578.1**](https://www.ncbi.nlm.nih.gov/protein/488984790) ***Klebsiella pneumoniae***

MTLDDLAFDRRHIWHPYTSMTSPLPVYPVVSAHGCELSLAGGEQLVDGMSSWWAAIHGYNHPRLNAALKGQIDQMSHVMFGGITHPPAVALCRQLVAMTPASLECVFLADSGSVAVEVAMKMALQYWQAKGEPRRRFLTFRNGYHGDTFGAMSVCDPQNSMHSLWQGYLPDNLFAPAPQSRFDGEWDEMDMVPFARLMAAHRHEIAAVILEPIVQGAGGMRMYHPEWLKRVRKMCDREGILLIADEIATGFGRTGKLFACEHAGITADILCLGKALTGGTMTLSAAITTRTVAETISNGEAGCFMHGPTFMGNPLACAVAGESLRLLESGEWQPQVTAIEAQLQAELAPARGSALVADVRVLGAIGVVETRRPVNMAALQRFFVEQGVWIRPFGRLIYLMPPYIITPEQLTRLTRAVNQAVQDETFFSE

**>WP_003117900.1 *Pseudomonas aeruginosa***

MGLNADWMQRDLNVLWHPCTQMKDHERLPVIPIRRGEGVWLEDFEGKRYIDAVSSWWVNVFGHANPRINQRIKDQVDQLEHVILAGFSHQPVIELSERLVKITPPGLDRVFYADSGSAGIEVALKMSYHFWLNSGRPRKKRFVTLTNSYHGETIAAMSVGDVALFTETYKSLLLDTIKVPSPDCFLRPDGMCWEEHSRNMFAHMERTLAEGHDEIAAVIVEPLIQGAGGMRMYHPVYLKLLREACDRYGVHLIHDEIAVGFGRTGTMFACEQAGIAPDFLCLSKALTGGYLPMSAVLTSETVYRGFYDDYQTLRAFLHSHTYTGNPLACAAALATLDIFEEDKVIEANRALSTHMARATAHLADHPHVAEVRQTGMVLAIEMVQDKASRTPYPWQERRGLKVFQHGLERGALLRPLGSVVYFLPPYVITPEQIDFLAEVASEGIDIATRDAVSVAVSDFHPDHRDPG

**>WP_002370098.1 *Enterococcus faecalis***

MKYNYLVPMGDITKVNEHKTTIVRAEEEYVFDEEQKRFVDLRSGLWNTNLGYKKELYEIIRQRFTEQLSKSLTYLDIHSFHHPVYQEYAKKLATFADKEGFYEQVIYTNSGSECTELALKISRQINKSNQKILAFSQGYHGTFWGGMSISGLDQEVTDIYSPKLSNMEFIKSPENDIEEKNFFKHIEYHHHEYSAMIIEPVLGSAGIKMPSIRFLNKLGSLLKKYGIIVIFDEVATGFYRTGKPFYFHYLDFKPDIINLSKGINNGMLPFGVVLLSNDIVCELKKEKLEHFSTQNGNLLGVISAYETLNYYRQHEVEIVQNIQNLNELILTELNVYGISFRGIGCMFAIPIDDKQALPLIIQSLKQTGILCYQYFNSDEDNGLTLMPSFYTNYQKMLQIIRRIAKVVNAYA

**>WP_001110064.1 *Staphylococcus aureus***

MNYTQQLKQKDSEYVWHPFTQMGVYSKEEAIIIEKGKGSYLYDTNGNKYLDGYASLWVNVHGHNNKYLNKVIKKQLNKIAHSTLLGSSNIPSIELAEKLIEITPSNLRKVFYSDTGSASVEIAIKMAYQYWKNIDREKYAKKNKFITLNHGYHGDTIGAVSVGGIKTFHKIFKDLIFENIQVESPSFYRSNYDTENEMMTAILTNIEQILIERNDEIAGFILEPLIQGATGLFVHPKGFLKEVEKLCKKYDVLLICDEVAVGFGRTGKMFACNHEDVQPDIMCLGKAITGGYLPLAATLTSKKIYNAFLSDSHGVNTFFHGHTYTGNQIVCTVALENIRLYEKRKLLSHIETTSSTLEKQLHALKRHRNVGDVRGRGLMFGVELVTDKDSKTPLEIEKVERIVRNCKENGLMIRNLENVITFVPVLSMSNKEVKTMVRIFKKAVHNILDRKC

**Supplemental Figure S3A**. NCBI Reference Sequence for Mtb and ESKAPE (*Enterobacter aerogenes, Staphylococcus aureus, Klebsiella pneumoniae, Acinetobacter baumannii, Pseudomonas aeruginosa and Enterococcus faecalis*) BioA protein.

```
Query: NP_216084.1 M. TUBERCULOSIS Query ID: lcl|Query_308311 Length: 437


Query range 1: 1 to 90

Query         11   EQIIAVDGAHLWHPYSSIGRE-AVSPVVAVA-A-HGAWLTLIRDGQPIEVLDAMSSWWTAIHGHG---HPALDQALTTQL-RVMNHVMFG  93
Query_308313  6          DLEHIWHPYTSMTQP-L--PTFKVKRA-YGATIEL-DDGRTL--IDGMSSWWCAIHGYN---HPELNQAVTDQL-QNMSHIMFG  78
Query_308316  6       LAFDRRHIWHPYTSMTSPLPVYPVVS---A-HGCELSLAGGEQ---LVDGMSSWWAAIHGYN---HPRLNAALKGQI-DQMSHVMFG  81
Query_308314  11     LLNFDRTHLLHPYTSMTDP-L--PVYPVKRA-EGVFIELA-DG--TRLIDGMSSWWCAIHGYN---HPVLNQAVENQM-KQMAHVMFG  87
Query_308315  6       LAFDRRHIWHPYTSMTSPLPVYPVVS---A-HGCELSLAGGEQ---LIDGMSSWWAAIHGYN---HPRLNAAMKAQI-DQMSHVMFG  81
Query_308317  15             LWHPCTQMKDH-ERLPVIPIRRG-EGVWL---EDFEGKRYIDAVSSWWVNVFGHA---NPRINQRIKDQV-DQLEHVILA  85
Query_308319  5    QQLKQKDSEYVWHPFTQMGVY-SKEEAIIIE-KGKGSYLY---DTNGNKYLDGYASLWVNVHGHN---NKYLNKVIKKQL-NKIAHSTLL  85
Query_308318  32                                              DEEQKRFVDLRSGLWNTNLGYKKELYEIIRQRFTEQLSKSLTYLDIH  78


Query range 2: 91 to 180

Query         94   GLTHEPAARLAKLLVDI--TPAGLDTVFFSDSGSVSVEVAAKMALQYWRG---RGLPGKRRLMTWRGGYHGDTFLAMSICDPHGGM---H  175
Query_308313  79   GLTHDPAIELGKILLKI--TPPSLDKIFYADSGSVAVEVALKMAVQFWTA---QGQPQKTNFITTRSGYHGDTWNAMSVCDPVTGM---H  160
Query_308316  82   GITHPPAVALCRQLVAM--TPASLECVFLADSGSVAVEVAMKMALQYWQA---KGEP-RRRFLTFRNGYHGDTFGAMSVCDPQNSM---H  162
Query_308314  88   GLTHEPAVELGKLLVGI--LPQGLDRIFYADSGSVSVEVALKMAVQYQQA---RGLTAKQNIATVRRGYHGDTWNAMSVCDPETGM---H  169
Query_308315  82   GITHPSAVALCRQLVAM--TPESLECVFLADSGSVAVEVAMKMALQYWQA---KGQP-RRRFLTFRNGYHGDTFGAMSVCDPQNSM---H  162
Query_308317  86   GFSHQPVIELSERLVKI--TPPGLDRVFYADSGSAGIEVALKMSYHFWLN---SGRPRKKRFVTLTNSYHGETIAAMSVGDV--------  162
Query_308319  86   GSSNIPSIELAEKLIEI--TPSNLRKVFYSDTGSASVEIAIKMAYQYWKNIDREKYAKKNKFITLNHGYHGDTIGAVSV----GGIKTFH  169
Query_308318  79   SFHHPVYQEYAKKLATFADKEGFYEQVIYTNSGSECTELALKISRQINKS---N-----QKILAFSQGYHGTFWGGMSISGLDQEV---T  157

Query range 3: 181 to 270

Query         176  SLWTDVL--AAQVF--AP-QVP-------------R-DYDP------AYSAA----FEAQLAQHAGELAAVVVEPVVQGAGGMRFHDPRY  236
Query_308313  161  QIFGTSL--PNRLFVAAP-QTKFH-----------E-EWNQ------EDIAE----LEQAIQQNHENLAALIIEPIVQGAGGMRFYHPEY  225
Query_308316  163  SLWQGYL--PDNLF--AP-APQ-------------S-RFDG------EWDEMDMVPFARLMAAHRHEIAAVILEPIVQGAGGMRMYHPEW  227
Query_308314  170  HIFGSAL--PQRYF--VD-NPK-------------N-RFDD------EWDGADLQPVRALFEAHHVDIAAFILEPVVQGAGGMYFYHPQY  234
Query_308315  163  SLWQGYL--PDNLF--AP-APQ-------------S-RFDG------EWDEMDMVPFARLMAAHRHEIAAVILEPVVQGAGGMRMYHPEW  227
Query_308317  163  ALFTETY--KSLLL--DTIKVPSPDCFLRPDGMCWE-EHSR------NMFAH----MERTLAEGHDEIAAVIVEPLIQGAGGMRMYHPVY  237
Query_308319  170  KIFKDLIFENIQVE--SP-SFY-------------RSNYDTENEMMTAILTN----IEQILIERNDEIAGFILEPLIQGATGLFVHPKGF  239
Query_308318  158  DIYSPKL--SNMEF--IK-SPE-------------N-DIE---------EKN----FFKHIEYHHHEYSAMIIEPVL-GSAGIKMPSIRF  214

Query range 4: 271 to 360

Query         237  LHDLRDICRRYEVLLIFDEIATGFGRTGALFAADHAGVSPDIMCVGKALTGGYLSLAATLCTADVAHTI--SAGAAG--ALMHGPTFMAN  322
Query_308313  226  LRQAKALCEKYHLLLIFDEIATGFGRTGKLFAWEHAQVEPDIMCLGKGLTGGYMTLSATLTTKHVAETI--SRGEAG--VFMHGPTFMAN  311
Query_308316  228  LKRVRKMCDREGILLIADEIATGFGRTGKLFACEHAGITADILCLGKALTGGTMTLSAAITTRTVAETI--SNGEAG--CFMHGPTFMGN  313
Query_308314  235  LRGLRDLCDEFDIVLIFDEIATGFGRTGKMFACEHAEVVPDIMCIGKGLSGGYMTLAAAITSQKVTETI--SRGEAG--VFMHGPTFMAN  320
Query_308315  228  LKRVRKMCDREGILLIADEIATGFGRTGKLFACEHAGISADILCLGKALTGGTMTLSAAITTRTVAETI--SNGEAG--CFMHGPTFMGN  313
Query_308317  238  LKLLREACDRYGVHLIHDEIAVGFGRTGTMFACEQAGIAPDFLCLSKALTGGYLPMSAVLTSETVYRGFYDDYQTLR--AFLHSHTYTGN  325
Query_308319  240  LKEVEKLCKKYDVLLICDEVAVGFGRTGKMFACNHEDVQPDIMCLGKAITGGYLPLAATLTSKKIYNAF--LSDSHGVNTFFHGHTYTGN  327
Query_308318  215  LNKLGSLLKKYGIIVIFDEVATGFYRTGKPFYFHYLDFKPDIINLSKGINNGMLPFGVVLLSNDIV--------------------------280

Query range 5: 361 to 450

Query         323  PLACAVSVASVELLLGQDWRTRITELAAGLTAGLDTARA----LPAVTDVRVCGAIGVIECDRPVDLAVATPA-------------ALDR  395
Query_308313  312  PLACAVALKSTQLLIEQDWQANIKRIEQQLSQYLMPLNQ----LDYVADVRVLGAIGVVELTFNVDMKTLQQQ-------------FVER  384
Query_308316  314  PLACAVAGESLRLLESGEWQPQVTAIEAQLQAELAPARG----SALVADVRVLGAIGVVETRRPVNMAALQRF-------------FVEQ  386
Query_308314  321  PLACAVACASVKLLLSQDWQANIRRIESILKGRLKAAWD----IRGVKDVRVLGAIGVIELEKGVDMARFQAD-------------CVAQ  393
Query_308315  314  PLACAVASESLRLLESGEWQQQVAAIEAQLKAELAPARE----SEWVADVRVLGAIGVVETRQPVNMAALQRF-------------FVEQ  386
Query_308317  326  PLACAAALATLDIFE----EDKVIEANRALSTHMARATAHLADHPHVAEVRQTGMVLAIEMVQ--DKASRTPYPWQERRGLKVFQHGLER  409
Query_308319  328  QIVCTVALENIRLYEKRKLLSHIETTSSTLEKQLHALKR----HRNVGDVRGRGLMFGVEL--VTDKDSKTPL-------------EIEK  398
Query_308318       ------------------------------------------------------------------------------------------

Query range 6: 451 to 500

Query         396  -----------GVWLRPFRNLVYAMPPYICTPAEI---TQITSAMVEVAR  431
Query_308313  385  -----------GIWIRPFGKLVYVMPPYVITQQEL---SDLLEHLVEVVK  420
Query_308316  387  -----------GVWIRPFGRLIYLMPPYIITPEQL---TRLTRAVNQAVQ  422
Query_308314  394  -----------GIWVRPFGRLVYLMPPYIISDGIL---TKLADKTVQILK  429
Query_308315  387  -----------GVWIRPFGRLIYLMPPYIISPQQL---TRLTRAV-----  417
Query_308317  410  -----------GALLRPLGSVVYFLPPYVITPEQIDFLAEVASEGIDIA-  447
Query_308319  399  VERIVRNCKENGLMIRNLENVITFVPVLSMSNKEV---------------  433
Query_308318        -----------------------------------------------------
```

**Supplemental Figure S3B**. Sequence alignment between *Mt*BioA and ESKAPE BioA

**Supplemental Table S6**. Sequence alignment between *Mt*BioA and ESKAPE BioA

\# blastp

\# Iteration: 0

\# Query: **NP_216084.1 *M. tuberculosis***

\# RID: PEJY8RH2114

\# Database: n/a

\# Fields:

| Query | Subject acc.vear. | % Identity | length | Mismatches | Gap opens | q. start | q. end | s. start | s. end | Evalue | Bit score | Positives |
| --- | --- | --- | --- | --- | --- | --- | --- | --- | --- | --- | --- | --- |
| *A. baumannii* | WP_000131427.1 | 51.429 | 420 | 194 | 6 | 17 | 431 | 6 | 420 | 4.02e-157 | 441 | 68.10 |
| *K. pneumoniae* | WP_002895578.1 | 52.009 | 423 | 192 | 5 | 14 | 431 | 6 | 422 | 1.09e-152 | 430 | 68.09 |
| *N. gonorrhoeae* | WP_003687694.1 | 50.708 | 424 | 199 | 5 | 13 | 431 | 11 | 429 | 2.20e-150 | 424 | 66.27 |
| *E. aerogenes* | WP_015367618.1 | 51.675 | 418 | 191 | 5 | 14 | 426 | 6 | 417 | 2.32e-150 | 424 | 67.70 |
| *P. aeruginosa* | WP_003117900.1 | 36.242 | 447 | 234 | 11 | 21 | 430 | 15 | 447 | 1.45e-85 | 259 | 53.69 |
| *S. aureus* | WP_001110064.1 | 31.963 | 438 | 260 | 11 | 11 | 419 | 5 | 433 | 2.50e-81 | 248 | 52.28 |
| *E.* *faecalis* | WP_002370098.1 | 24.806 | 258 | 179 | 6 | 51 | 302 | 32 | 280 | 4.64e-25 | 95.5 | 45.35 |

**Supplemental Table S7**. Analysis of active site between *Mt*BioA and ESKAPE BioA

| **Query number** | **Organism/accession** | **Identity of active site residues** | **Number of mutations and deletions among the 31 residues with 5Å of ligand** |
| --- | --- | --- | --- |
| **Query** | *M. tuberculosis/*  WP_003687694.1 | n.a. | n.a. |
| **Query_308313** | *A. baumannii/*  WP_015367618.1 | G155S, G172T, R403G | 3 nonsynonymous mutations (all and non-conservative) |
| **Query_308316** | *Klebsiella pneumoniae/*  WP_002895578.1 | G155N, G172N, G173S, R403G | 4 nonsynonymous mutations (all and non-conservative) |
| **Query_308314** | *Neisseria gonorrhoeae/*  WP_003687694.1 | G155R, G172T, R403G | 3 nonsynonymous mutations (all and non-conservative) |
| **Query_308315** | *Enterobacter aerogenes/* WP_015367618.1 | G155N, G172N, G173S, R403G | 4 nonsynonymous mutations |
| **Query_308317** | *Pseudomonas aeruginosa/* WP_003117900.1 | Y25C, M91I, F92L, G93A, G155N, G156S, C168G, P170V, G172-, G173-, M174-, M175-, M314L, G316S, P317H, M320T, F402L, R403G | 14 nonsynonymous mutations (3 conservative and 11 non-conservative),  4 deletions |
| **Query_308319** | *Staphylococcus aureus/*  WP_001110064.1 | Y21F, W64L, M91T, F92L, G93L, G155H, M165V, C168-, D169-, P170-, M174I, G227T, I256V, M314F, P317H, M320T, F402L, R403E, | 15 nonsynonymous mutations (3 conservative and 12 non-conservative),  3 deletions |
| **Query_308318** | *Enterococcus faecalis/*  WP_002370098.1 | P24-, Y25-, W64L, M91D, F92I, G93H, G155Q, C168S, D169G, P170L, G172Q, G173E, M174V, H175T, A226S, G227A, I256V, M314-, G316-, P317-, T318-, M320-, R400-, F402-, R403-, Y407- | 15 nonsynonymous mutations (3 conservative and 12 non-conservative),  11 deletions |

Amino acid residues within 5 Å of *M. tuberculosis* BioA active site

- Red is the bromo-fluorophenyl
- orange is the piperazine
- green is the azaindolone
- blue interacts with PLP cofactor

P24, **Y25**, **W64, W65**, M91, F92, G93; G155, G156, **Y157,** M165, C168, D169, P170, G172, G173, M174, H175, A226, G227, **I256**, **K283,** M314, G316, P317, T318, M320, R400, **F402**, R403, **Y407**

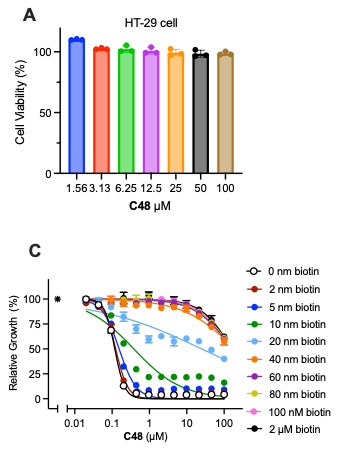

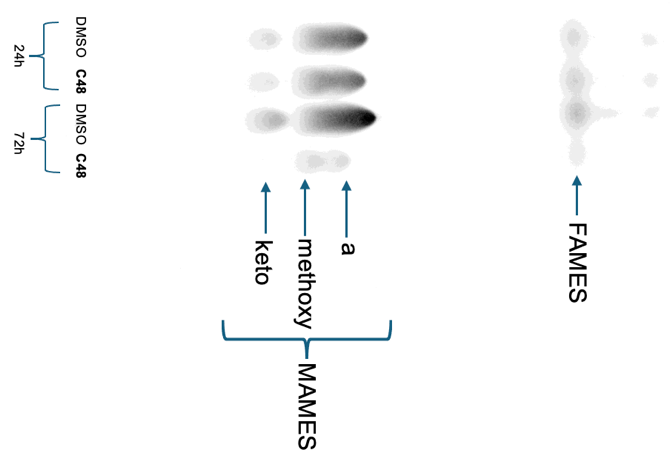

**B**

### **Supplemental Figure S4. Cytotoxicity of C48**

(**A**) Viability of HT-29 after 48 h treatment with the indicated concentrations of **C48**. Values are normalized to vehicle treated condition (DMSO). The assay was performed as triplicate. (**B**) TLC analysis of [^14^C]acetate-labeled FAMEs and MAMEs after **C48** (100 nM) or control (DMSO) treatment at indicated time. a = α -mycolic esters, methoxy = methoxy mycolic esters, keto = Keto-mycolic esters. (**C**) The potency of **C48** against WT Mtb was antagonized in the presence of biotin. Asterisk represents no drug control.

### **Supplemental Table S8.** **C48** Resistant Mutants and AA change

| Sample | Class | AA change | MIC shift |
| --- | --- | --- | --- |
| 1 | High-level resistance 1 (FOR10) | Met91Ile | Large |
| 2 | High-level resistance 2 (FOR22) | Met91Ile | Large |
| 3 | High-level resistance 4 (KK1) | Met91Ile | Large |
| 4 | Medium-level resistance 4 (KK2) | Met91Val | Medium |
| 5 | High-level resistance 5 (KK3) | Met91Ile | Large |
| 6 | High-level resistance 6 (KK4) | Met91Ile | Large |
| 7 | Medium-level resistance 1 (FA5) | Met91Val | Medium |
| 8 | High-level resistance 7 (FA9) | Met91Ile | Large |
| 9 | Medium-level resistance 5 (FA13) | Met91Thr | Medium |
| 10 | High-level resistance 3 (FA19) | Met91Ile | Large |
| 11 | High-level resistance 8 (FA23) | Met91Ile | Large |
| 12 | Medium-level resistance 6 (FA30) | Met91Thr | Medium |
| 13 | Medium-level resistance 2 (FA35) | Met91Val | Medium |
| 14 | High-level resistance 9 (FA37) | Met91Ile | Large |
| 15 | High-level resistance 10 (FA44) | Met91Ile | Large |
| 16 | Medium-level resistance 3 (FA49) | Cys168Tyr | Medium |
| FOR = resistance mutants were generated from frequency of resistance assay. FA = resistance mutants were generated from fluctuation assay. KK = resistance mutants were generated from kill kinetics study. AA change = amino acid change. MIC shift = MIC shift compared to WT. | | | |

### **Chemistry**

#### **^1^H and ^13^C NMR Spectra**

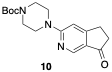

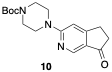

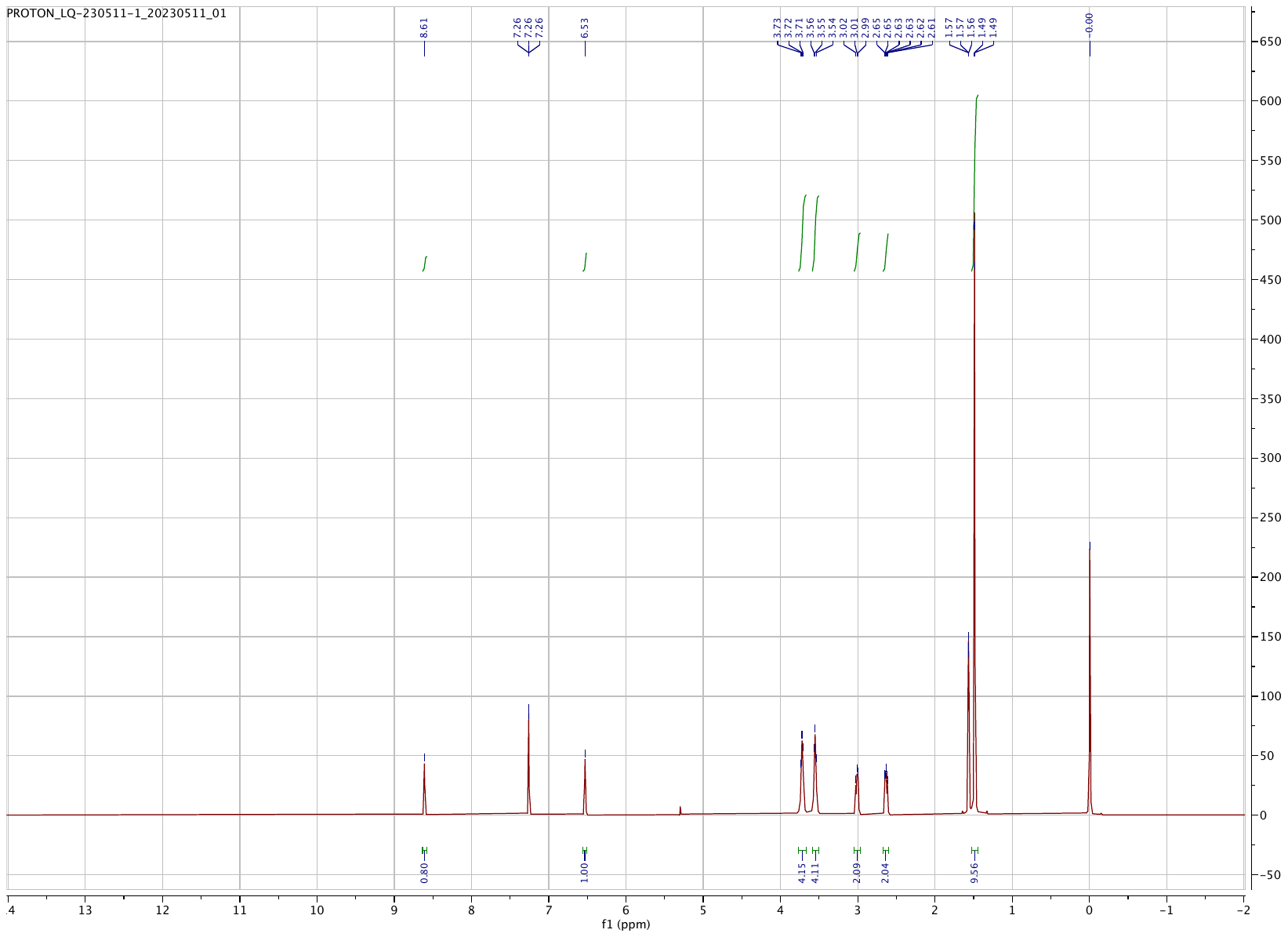

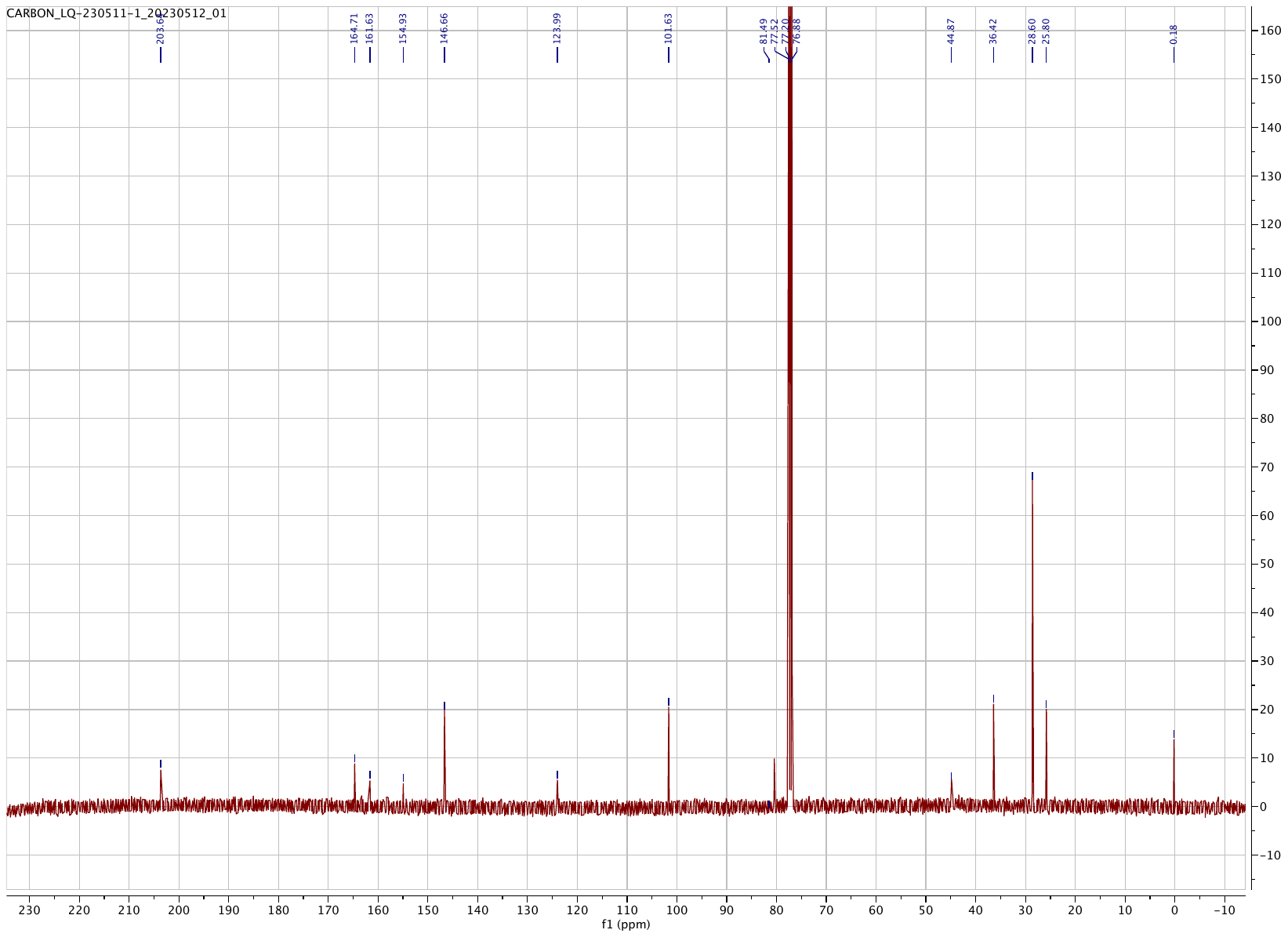

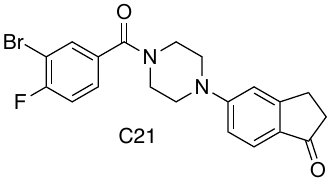

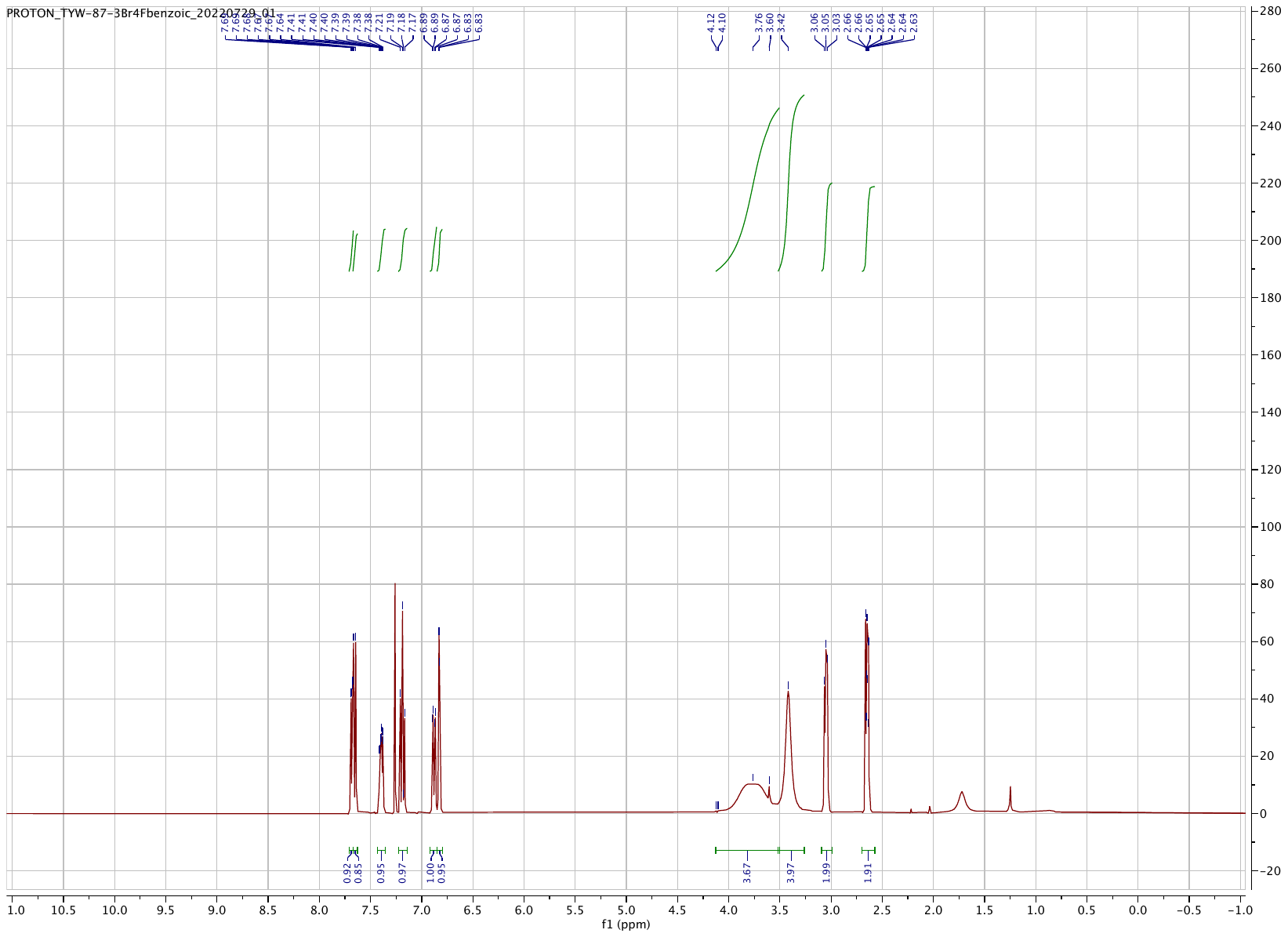

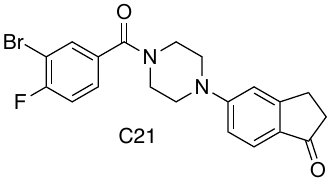

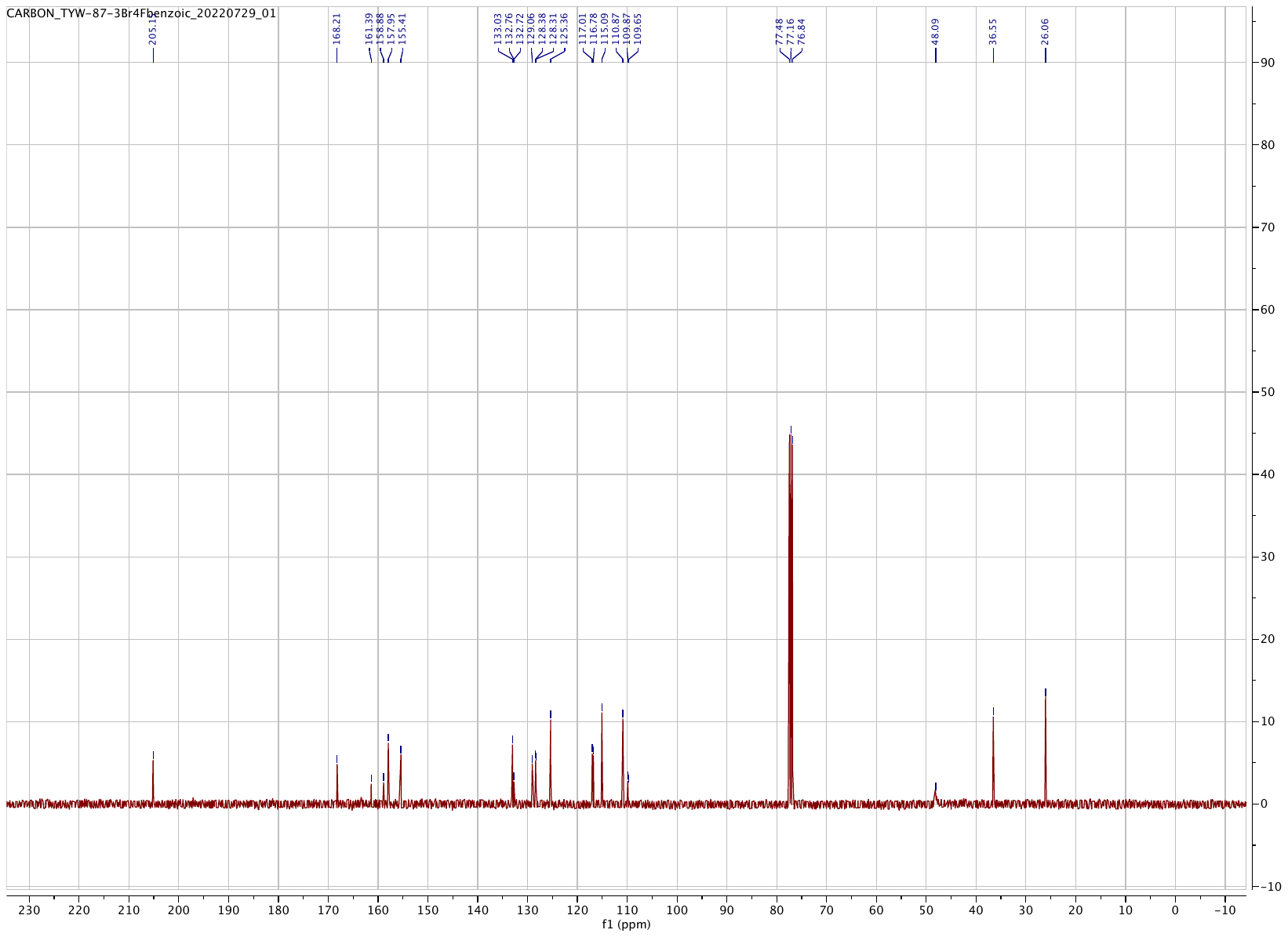

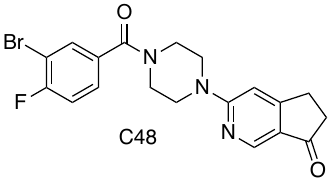

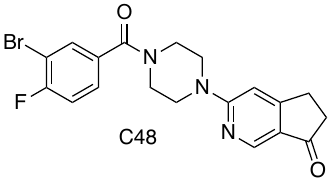

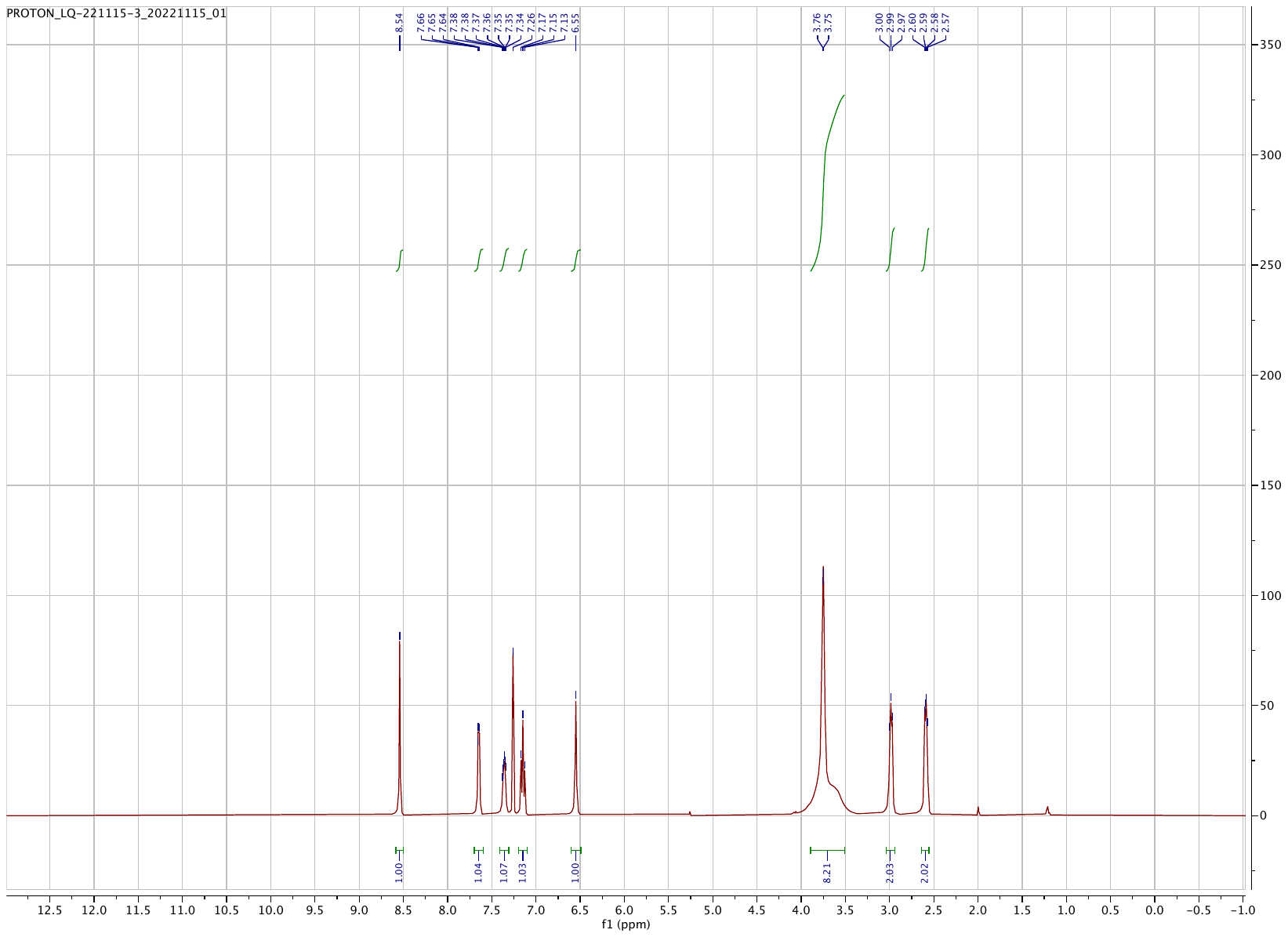

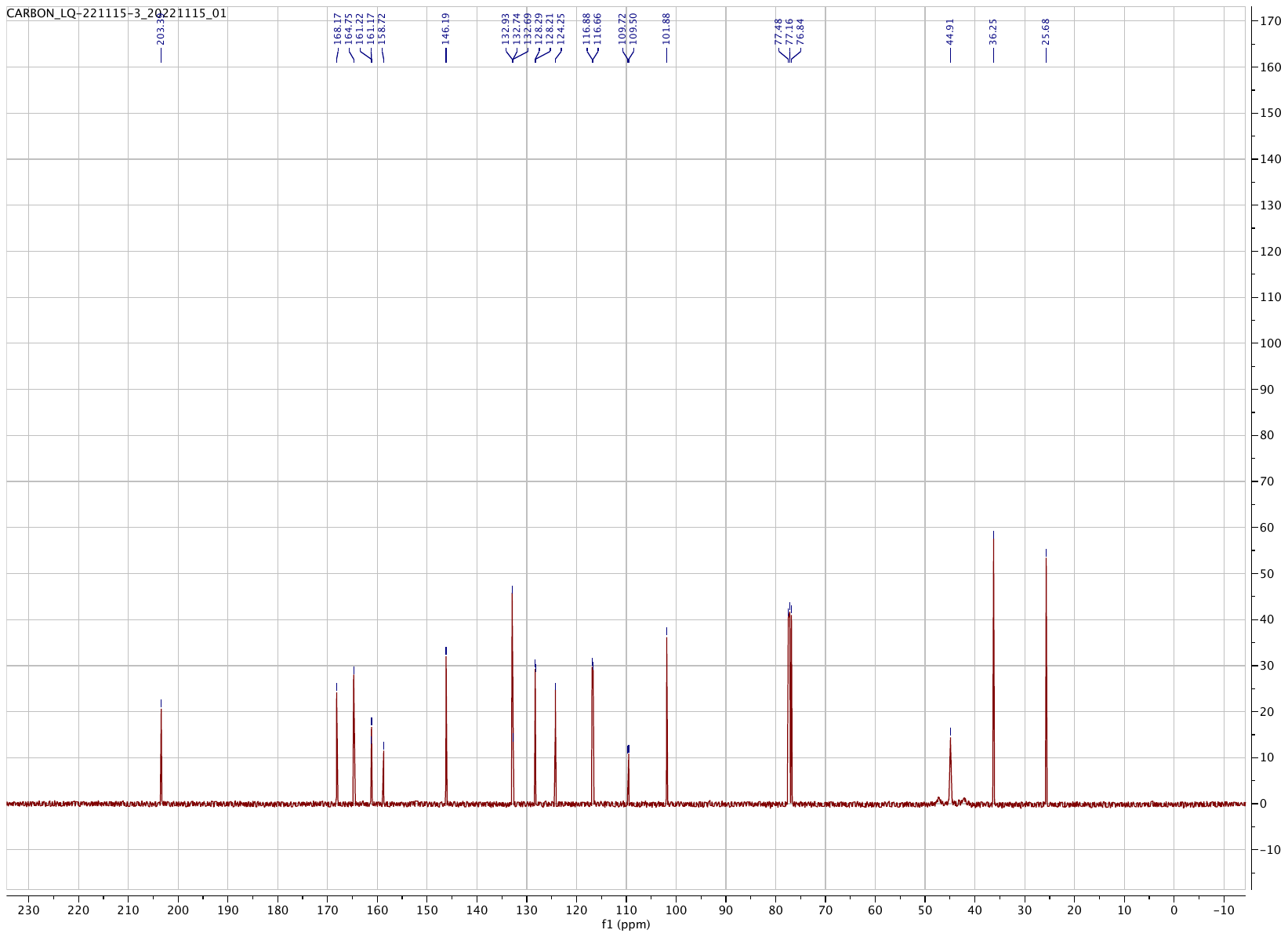

#### **HSQC of C48**

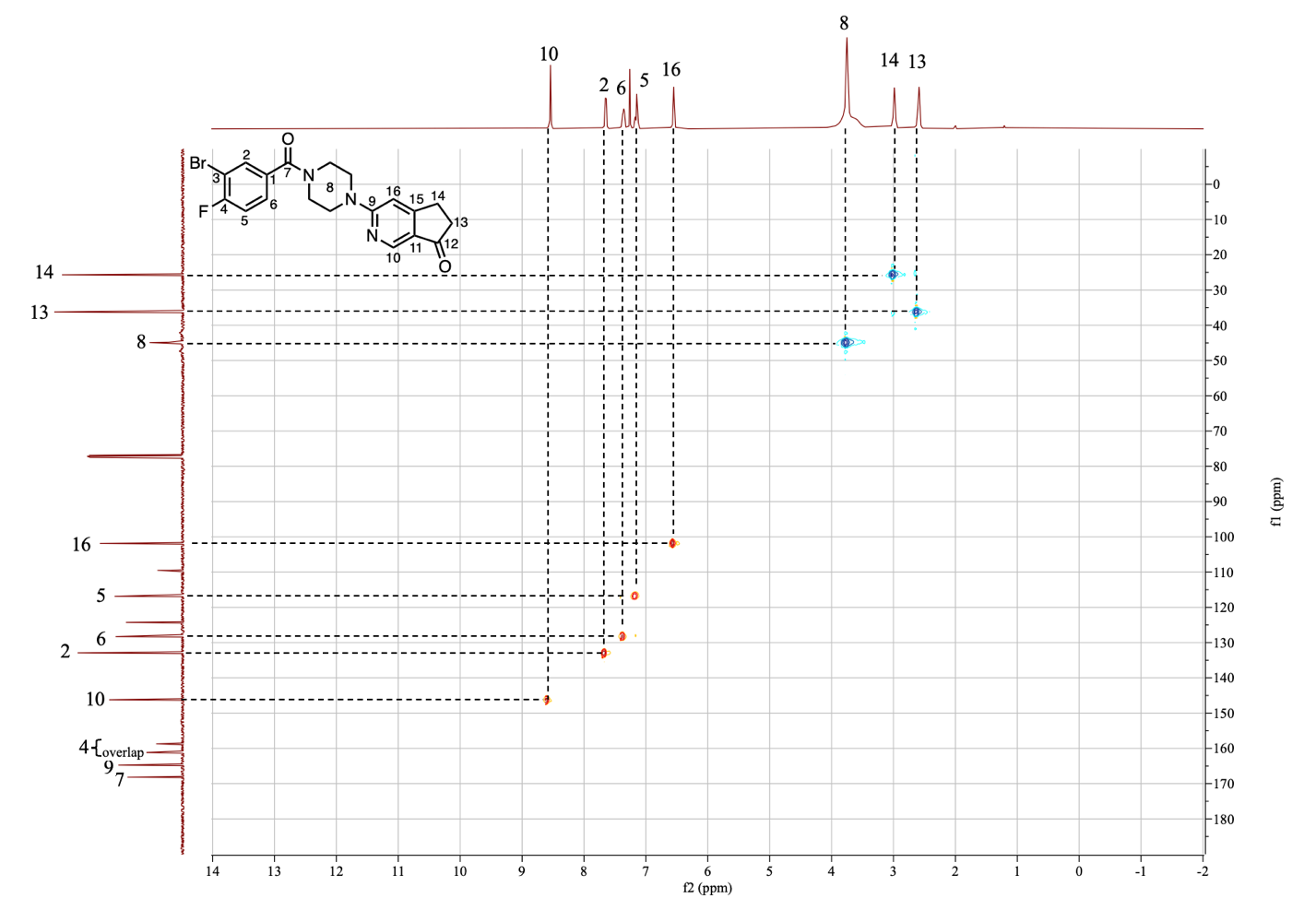

#### **^19^F NMR of C48**

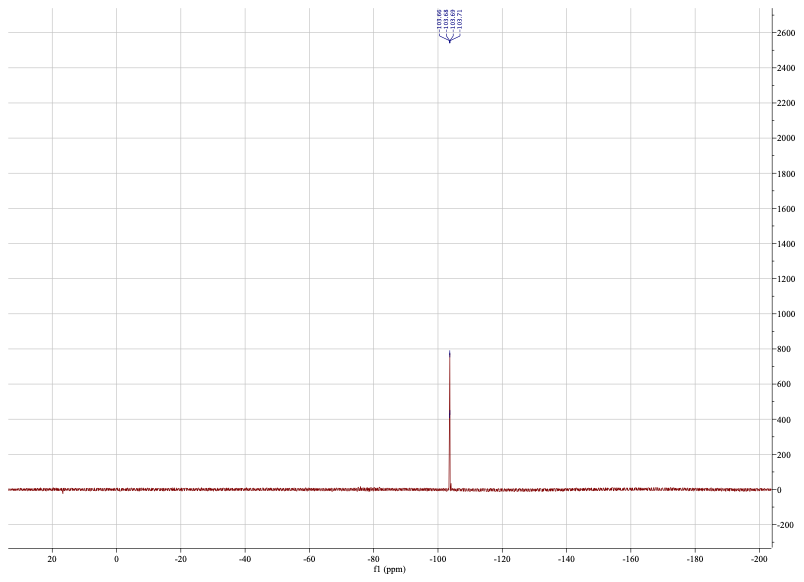

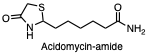

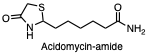

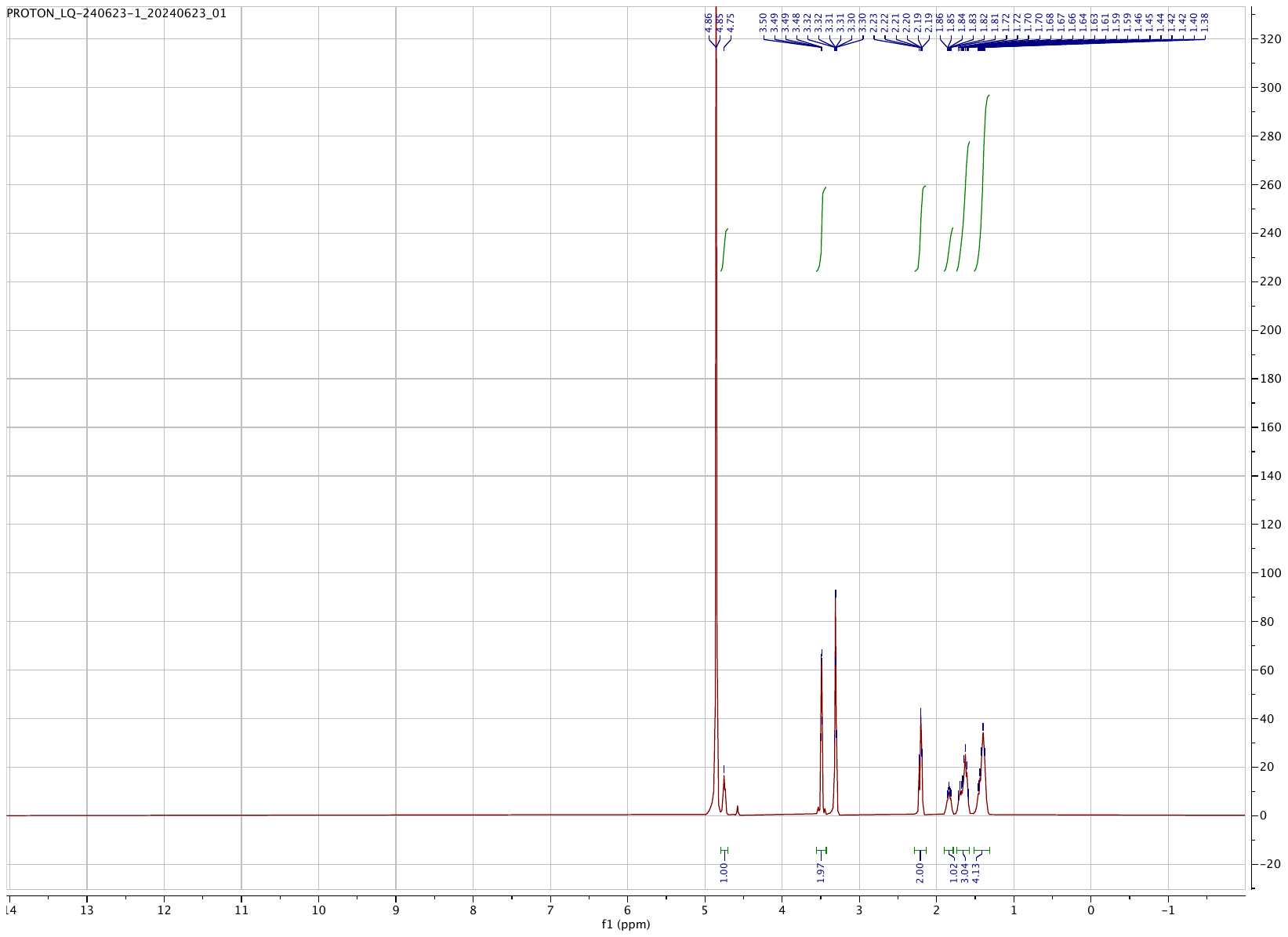

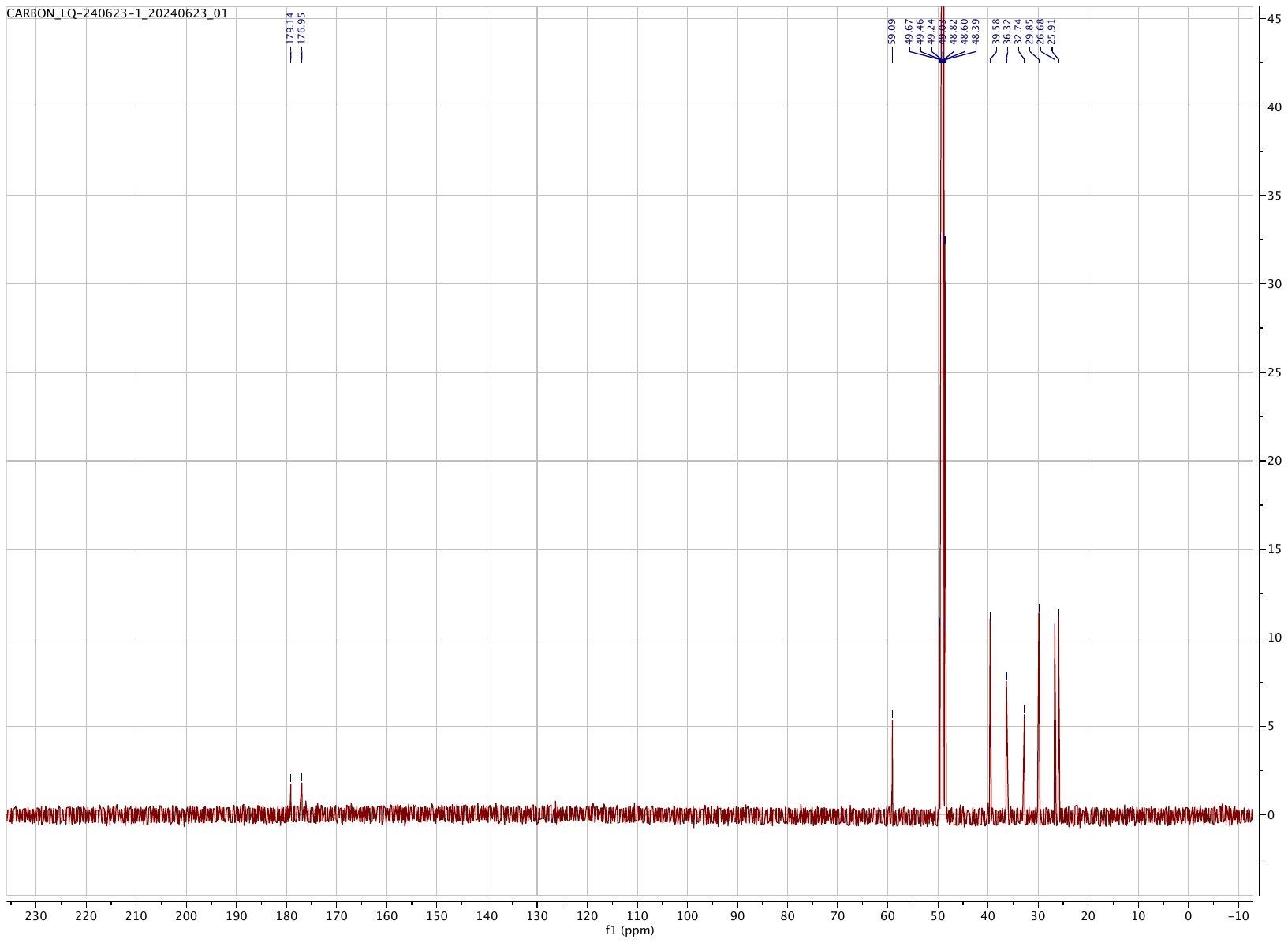

#### **HPLC Trace of C21**

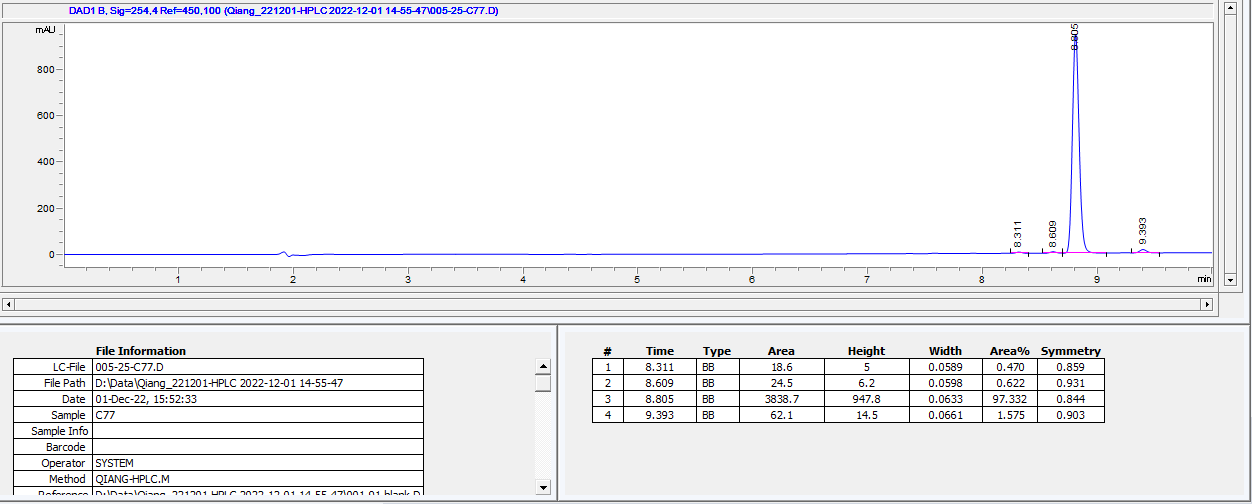

#### **HPLC Trace of C48**

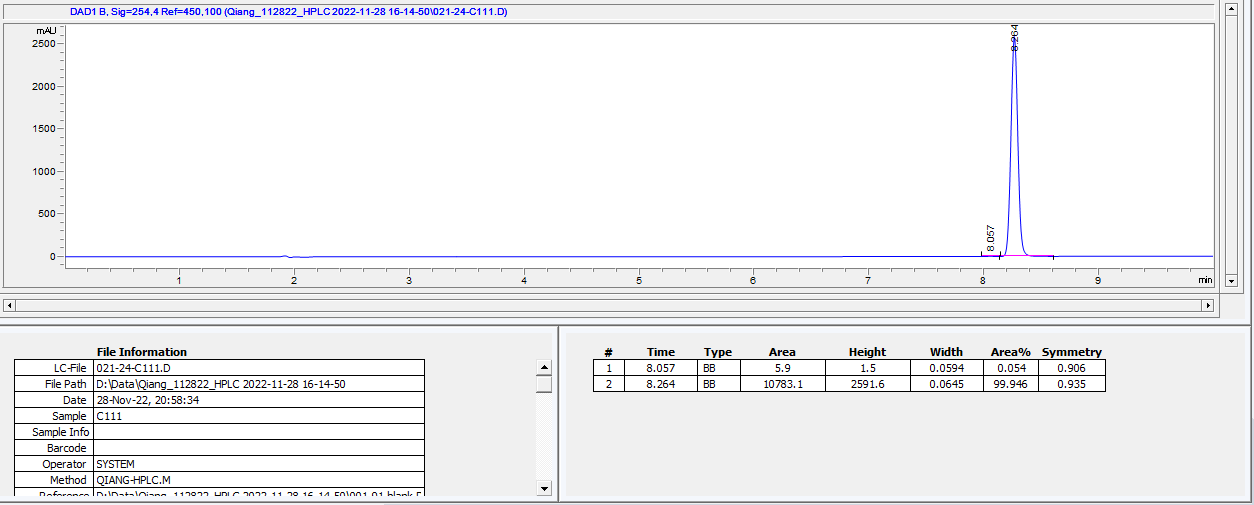

#### **HRMS of C21**

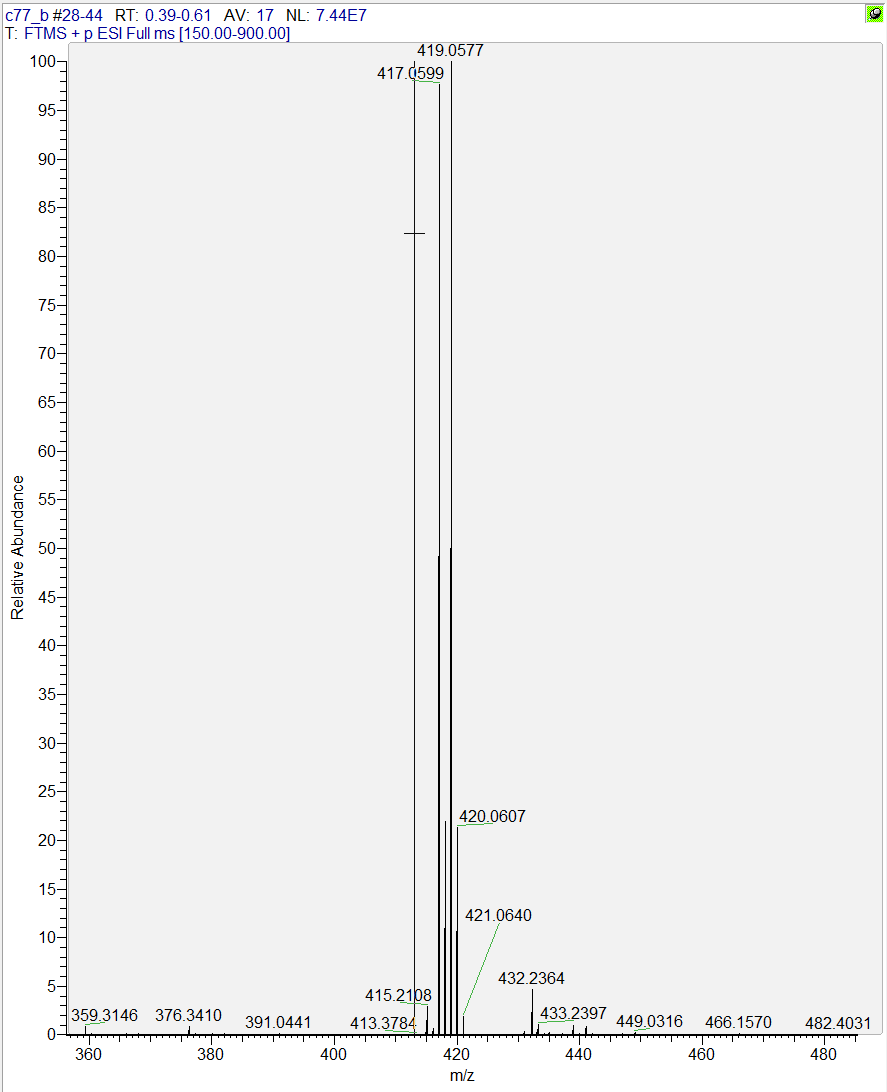

#### **HRMS of compound 10**

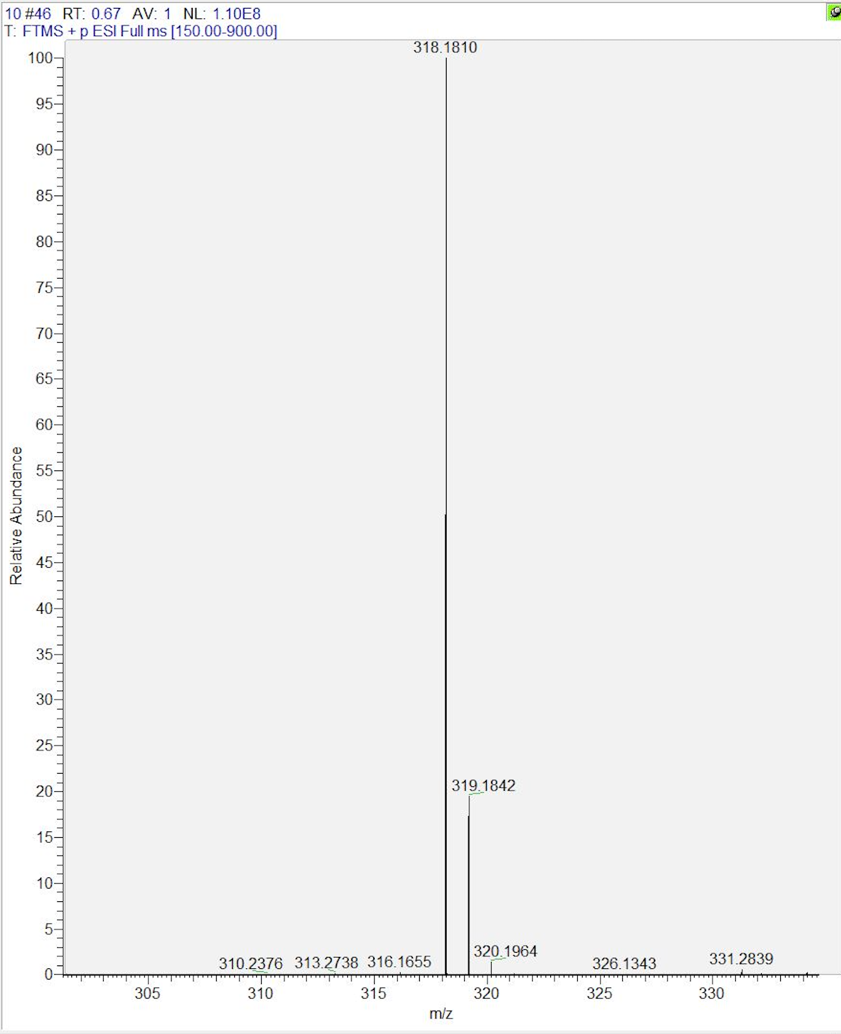

#### **HRMS of C48**

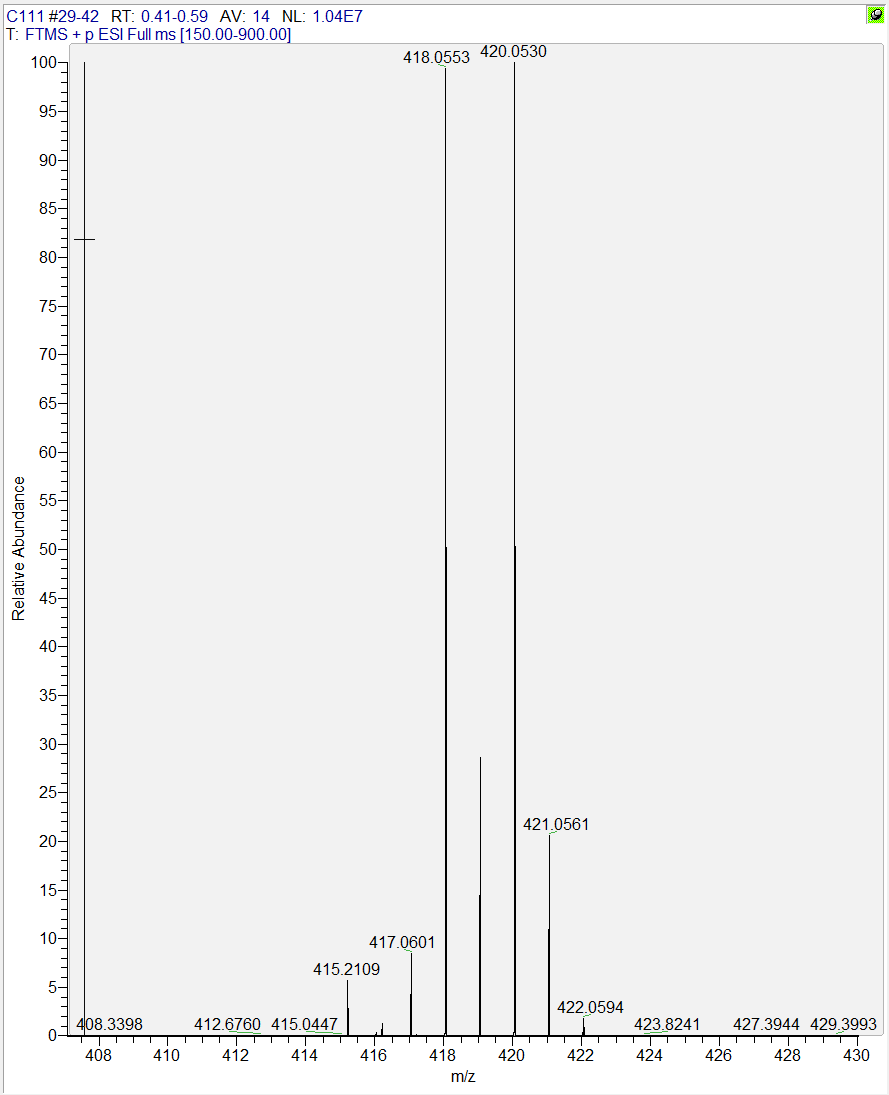

(14) Malherbe, S. T.; Chen, R. Y.; Yu, X.; Smith, B.; Liu, X.; Gao, J.; Diacon, A. H.; Dawson, R.; Tameris, M.; Zhu, H.; Qu, Y.; Jin, H.; Pan, S.; Dodd, L. E.; Wang, J.; Goldfeder, L. C.; Cai, Y.; Arora, K.; Vincent, J.; Narunsky, K.; Serole, K.; Goliath, R. T.; Da Costa, L.; Taliep, A.; Aziz, S.; Daroowala, R.; Thienemann, F.; Mukasa, S.; Court, R.; Sossen, B.; Ahlers, P.; Mendelsohn, S. C.; White, L.; Gouel, A.; Lau, C. Y.; Hassan, S.; Liang, L.; Duan, H.; Moghaddam, G. K.; Paripati, P.; Lahouar, S.; Harris, M.; Wollenberg, K.; Jeffrey, B.; Tartakovsky, M.; Rosenthal, A.; Duvenhage, M.; Armstrong, D. T.; Song, T.; Winter, J.; Gao, Q.; Via, L. E.; Wilkinson, R. J.; Walzl, G.; Barry, C. E., 3rd, PET/CT guided tuberculosis treatment shortening: a randomized trial. *medRxiv* **2024**.

(15) Song, T.; Park, Y.; Shamputa, I. C.; Seo, S.; Lee, S. Y.; Jeon, H. S.; Choi, H.; Lee, M.; Glynne, R. J.; Barnes, S. W.; Walker, J. R.; Batalov, S.; Yusim, K.; Feng, S.; Tung, C. S.; Theiler, J.; Via, L. E.; Boshoff, H. I.; Murakami, K. S.; Korber, B.; Barry, C. E., 3rd; Cho, S. N., Fitness costs of rifampicin resistance in Mycobacterium tuberculosis are amplified under conditions of nutrient starvation and compensated by mutation in the beta' subunit of RNA polymerase. *Mol. Microbiol.* **2014**, *91* (6), 1106-19.

(16) Bang, H.; Park, S.; Hwang, J.; Jin, H.; Cho, E.; Kim, D. Y.; Song, T.; Shamputa, I. C.; Via, L. E.; Barry, C. E.; Cho, S. N.; Lee, H., Improved rapid molecular diagnosis of multidrug-resistant tuberculosis using a new reverse hybridization assay, REBA MTB-MDR. *J. Med. Microbiol.* **2011**, *60* (Pt 10), 1447-1454.

(17) Akusobi, C.; Choudhery, S.; Benghomari, B. S.; Wolf, I. D.; Singhvi, S.; Ioerger, T. R.; Rubin, E. J., Transposon-sequencing across multiple Mycobacterium abscessus isolates reveals significant functional genomic diversity among strains. *mBio.* **2025**, *16* (2), e0337624.

(18) Liu, Q.; Engelhart, C. A.; Wallach, J. B.; Tiwari, D.; Ge, P.; Manna, A.; Panda, S.; McCue, W. M.; Wong, T. Y.; Sharma, S.; Jayasinghe, Y. P.; Fuller, J.; Ronning, D. R.; Bockman, M. R.; Cheung, A.; Dartois, V.; Zimmerman, M. D.; Schnappinger, D.; Aldrich, C. C., Metabolically Stable Adenylation Inhibitors of Biotin Protein Ligase as Antibacterial Agents. *J. Med. Chem.* **2025**, *68* (3), 3065-3087.
